# Inhibition of SARS-CoV-2 main protease by allosteric drug-binding

**DOI:** 10.1101/2020.11.12.378422

**Authors:** Sebastian Günther, Patrick Y. A. Reinke, Yaiza Fernández-García, Julia Lieske, Thomas J. Lane, Helen M. Ginn, Faisal H. M. Koua, Christiane Ehrt, Wiebke Ewert, Dominik Oberthuer, Oleksandr Yefanov, Susanne Meier, Kristina Lorenzen, Boris Krichel, Janine-Denise Kopicki, Luca Gelisio, Wolfgang Brehm, Ilona Dunkel, Brandon Seychell, Henry Gieseler, Brenna Norton-Baker, Beatriz Escudero-Pérez, Martin Domaracky, Sofiane Saouane, Alexandra Tolstikova, Thomas A. White, Anna Hänle, Michael Groessler, Holger Fleckenstein, Fabian Trost, Marina Galchenkova, Yaroslav Gevorkov, Chufeng Li, Salah Awel, Ariana Peck, Miriam Barthelmess, Frank Schlünzen, P. Lourdu Xavier, Nadine Werner, Hina Andaleeb, Najeeb Ullah, Sven Falke, Vasundara Srinivasan, Bruno Alves Franca, Martin Schwinzer, Hévila Brognaro, Cromarte Rogers, Diogo Melo, Joanna I. Zaitseva-Kinneberg, Juraj Knoska, Gisel E. Peña Murillo, Aida Rahmani Mashhour, Filip Guicking, Vincent Hennicke, Pontus Fischer, Johanna Hakanpää, Jan Meyer, Phil Gribbon, Bernhard Ellinger, Maria Kuzikov, Markus Wolf, Andrea R. Beccari, Gleb Bourenkov, David von Stetten, Guillaume Pompidor, Isabel Bento, Saravanan Panneerselvam, Ivars Karpics, Thomas R. Schneider, Maria Marta Garcia Alai, Stephan Niebling, Christian Günther, Christina Schmidt, Robin Schubert, Huijong Han, Juliane Boger, Diana C. F. Monteiro, Linlin Zhang, Xinyuanyuan Sun, Jonathan Pletzer-Zelgert, Jan Wollenhaupt, Christian G. Feiler, Manfred S. Weiss, Eike-Christian Schulz, Pedram Mehrabi, Katarina Karničar, Aleksandra Usenik, Jure Loboda, Henning Tidow, Ashwin Chari, Rolf Hilgenfeld, Charlotte Uetrecht, Russell Cox, Andrea Zaliani, Tobias Beck, Matthias Rarey, Stephan Günther, Dusan Turk, Winfried Hinrichs, Henry N. Chapman, Arwen R. Pearson, Christian Betzel, Alke Meents

**Affiliations:** Center for Free-Electron Laser Science, DESY, Notkestrasse 85, 22607 Hamburg, Germany; Bernhard Nocht Institute for Tropical Medicine, Bernhard-Nocht-Straße 74, 20359 Hamburg, Germany; Diamond Light Source Ltd. Diamond House, Harwell Science and Innovation Campus, Didcot, OX11 0DE, UK; Universität Hamburg, Center for Bioinformatics, Bundesstr. 43, 20146 Hamburg, Germany; Hamburg Centre for Ultrafast Imaging (CUI), Universität Hamburg, Luruper Chaussee 149, 22761 Hamburg, Germany; Universität Hamburg, Institut für Nanostruktur- und Festkörperphysik, Luruper Chaussee 149, 22761 Hamburg, Germany; European XFEL GmbH. Holzkoppel 4, 22869 Schenefeld, Germany; Heinrich Pette Institute, Leibniz Institute for Experimental Virology, Martinistraße 52, 20251 Hamburg, Germany; Max Planck Institute for Molecular Genetics, Ihnestraße 63-73, 14195 Berlin, Germany; Universität Hamburg, Department of Chemistry, Institute of Physical Chemistry, Grindelallee 117, 20146 Hamburg, Germany; Max Planck Institute for the Structure and Dynamics of Matter, Luruper Chaussee 149, 22761 Hamburg, Germany; Deutsches Elektronen Synchrotron (DESY), Photon Science, Notkestrasse 85, 22607, Hamburg, Germany; Vision Systems, Hamburg University of Technology, 21071 Hamburg, Germany; Division of Biology and Biological Engineering, California Institute of Technology, Pasadena, CA 91125, USA; Universität Hamburg, Department of Chemistry, Institute of Biochemistry and Molecular Biology and Laboratory for Structural Biology of Infection and Inflammation, c/o DESY, 22607 Hamburg, Germany; Fraunhofer Institute for Translational Medicine and Pharmacology (ITMP), Schnackenburgallee 114, 22525 Hamburg, Germany; Dompé Farmaceutici SpA, 67100 L’Aquila, Italy; EMBL Outstation Hamburg, c/o DESY, Notkestrasse 85, 22607 Hamburg, Germany; Institute of Molecular Medicine, University of Lübeck, 23562 Lübeck, Germany; Hauptmann Woodward Medical Research Institute, 700 Ellicott Street, Buffalo, NY, 14203, USA; German Center for Infection Research (DZIF), Hamburg-Lübeck-Borstel-Riems Site, University of Lübeck, 23562 Lübeck, Germany; Helmholtz Zentrum Berlin, Macromolecular Crystallography, Albert-Einstein-Str. 15, 12489 Berlin, Germany; Department of Biochemistry & Molecular & Structural Biology, Jozef Stefan Institute, Jamova 39, 1 000 Ljubljana, Slovenia; Centre of excellence for Integrated Approaches in Chemistry and Biology of Proteins (CIPKEBIP), Jamova 39, 1 000 Ljubljana,Slovenia; Universität Hamburg, Department of Chemistry, Institute of Biochemistry and Molecular Biology, Martin-Luther-King-Platz 6, 20146 Hamburg, Germany; Research Group for Structural Biochemistry and Mechanisms, Department of Structural Dynamics, Max Planck Institute for Biophysical Chemistry, Am Fassberg 11, 37077 Göttingen, Germany; Institute for Organic Chemistry and BMWZ, Leibniz University of Hannover, Schneiderberg 38, 30167 Hannover, Germany; Universität Greifswald, Institute of Biochemistry, Felix-Hausdorff-Strasse 4, 17489 Greifswald, Germany; Universität Hamburg, Department of Physics, Luruper Chaussee 149, 22761 Hamburg, Germany

## Abstract

The coronavirus disease (COVID-19) caused by SARS-CoV-2 is creating tremendous health problems and economical challenges for mankind. To date, no effective drug is available to directly treat the disease and prevent virus spreading. In a search for a drug against COVID-19, we have performed a massive X-ray crystallographic screen of two repurposing drug libraries against the SARS-CoV-2 main protease (M^pro^), which is essential for the virus replication and, thus, a potent drug target. In contrast to commonly applied X-ray fragment screening experiments with molecules of low complexity, our screen tested already approved drugs and drugs in clinical trials. From the three-dimensional protein structures, we identified 37 compounds binding to M^pro^. In subsequent cell-based viral reduction assays, one peptidomimetic and five non-peptidic compounds showed antiviral activity at non-toxic concentrations. We identified two allosteric binding sites representing attractive targets for drug development against SARS-CoV-2.

## Main Text

Infection of host cells by SARS-CoV-2 is critically governed by the complex interplay of several molecular factors of both the host and the virus(*1, 2*). Coronaviruses are RNA-viruses with a genome of approximately 30,000 nucleotides. The viral open-reading frames, essential for replication of the virus, are expressed as two overlapping large polyproteins, which must be separated into functional subunits for replication and transcription activity(*1*). This proteolytic cleavage, which is vital for viral reproduction, is primarily accomplished by the main protease (M^pro^), also known as 3C-like protease 3CL^pro^ or nsp5. M^pro^ cleaves the viral polyprotein pp1ab at eleven distinct sites. The core cleavage motif is Leu-Gln↓(Ser/Ala/Gly)(*1*). M^pro^ possesses a chymotrypsin-like fold appended with a C-terminal helical domain, and harbors a catalytic dyad comprised of Cys145 and His41(*1*). The active site is located in a cleft between the two N-terminal domains of the three-domain structure of the monomer, while the C-terminal helical domain is involved in regulation and dimerization of the enzyme, with a dissociation constant of ∼2.5 µM(*1*). Due to its central and vital involvement in virus replication, M^pro^ is recognized as a prime target for antiviral drug discovery and compound screening activities aiming to identify and optimize drugs which can tackle coronavirus infections(*3*). Indeed, a number of recent publications confirm the potential of targeting M^pro^ for inhibition of virus replication(*1, 2*).

A rational approach to the identification of new drugs is structure-based drug design(*4, 5*). The first step is target selection followed by biochemical and biophysical characterization and its structure determination. This knowledge forms the basis for subsequent *in silico* screening of up to millions of potential drug molecules, leading to the identification of potentially binding compounds. The most promising candidates are then subjected to screening *in vitro* for biological activity. Lead structures are derived from common structural features of these biologically active compounds. Further chemical modifications of lead structures can then create a drug candidate that can be tested in animal models and, finally, clinical trials.

In order to speed up this process and find drug candidates against SARS-CoV-2, we performed a massive X-ray crystallographic screen of the virus’ main protease against two repurposing libraries containing in total 5953 unique compounds from the “Fraunhofer IME Repurposing Collection”(*6*) and the “Safe-in-man” library from Dompé Farmaceutici S.p.A. Analysis of the derived electron-density maps revealed 37 structures with bound compounds. Further validation by native mass spectrometry and viral reduction assays led to the identification of six of those compounds showing significant *in vitro* antiviral activity against SARS-CoV-2, including inhibitors binding at allosteric sites.

In contrast to crystallographic fragment-screening experiments that use small molecules of low molecular weight typically below 200 Da, the repurposing libraries are chemically more complex and contain compounds twice the molecular weight (Fig. 1A) and thus likely to bind more specifically and with higher affinity(*7*). Due to the higher molecular weights, we performed co-crystallization experiments instead of compound soaking into native crystals(*8*). Crystals were grown at a physiological pH-value of 7.5.

**Fig. 1.**
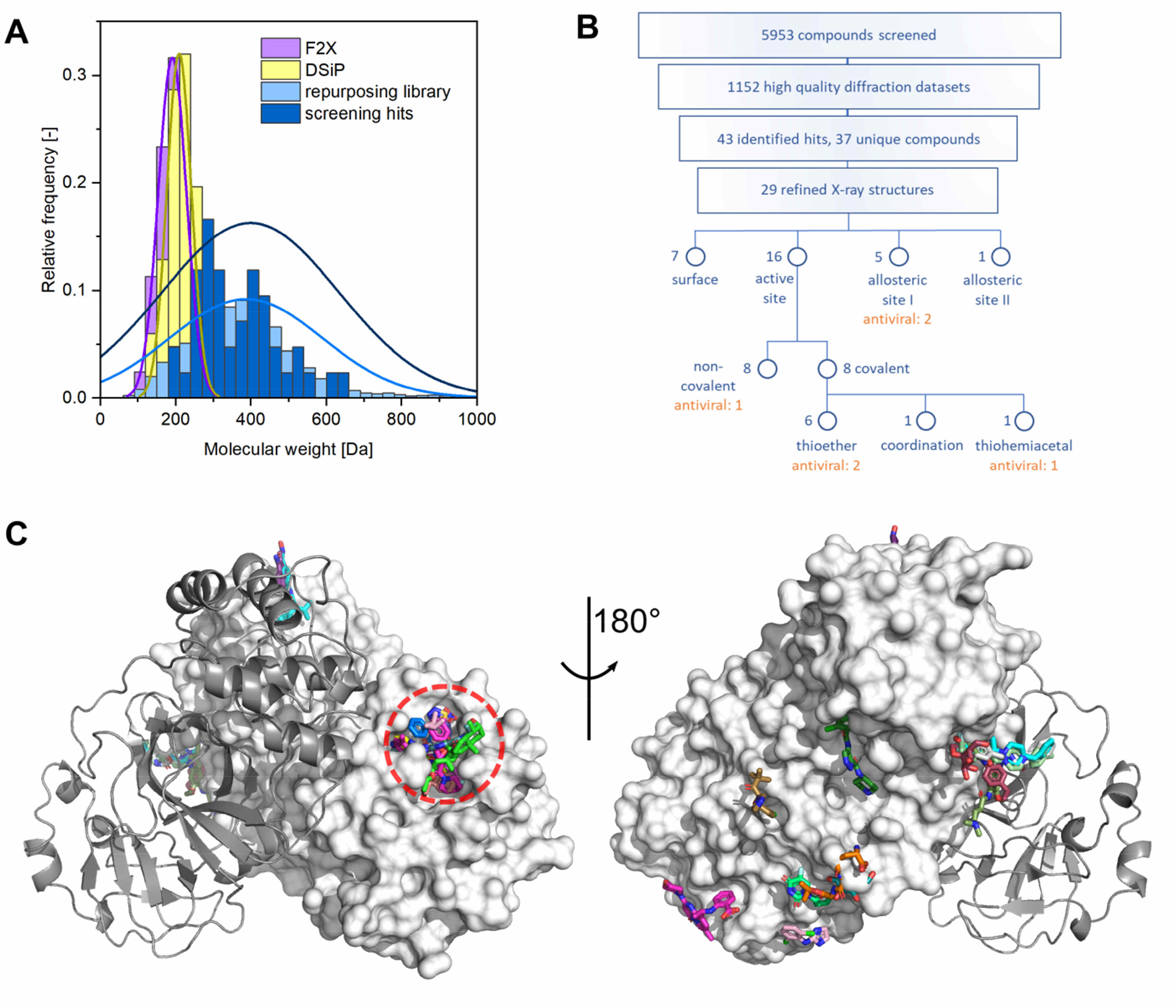
The repurposing libraries reveal compound binding sites distributed across the complete M^pro^ surface. **A**, Normalized histograms of molecular weight distributions of two commonly used fragment screening libraries F2X-Universal(*57*) (median 193.2 Da) and DSiP (a version of the “poised library”(*58*), 211.2 Da), the two combined repurposing libraries used in the present effort (Fraunhofer IMG, 371.3 Da, Dompé “Safe-in-man” 316.3 Da, combined 366.5 Da), and the resulting hits from our X-ray screen (403.6 Da). Normal distributions are indicated by solid lines in corresponding colors. Compounds with a molecular weight above 1000 Da are not shown. **B**, Flow chart with overview of analyzed compounds and identified hits compounds including classification of binding sites. **C**, Cartoon representation of M^pro^ with all unambiguously bound compounds. One protomer of the native dimer is depicted as a cartoon and the other one as surface representation. Left panel, view of the active site of M^pro^(dashed circle), right panel, view of M^pro^ rotated by 180°.

X-ray data collection was performed at beamlines P11, P13 and P14 at the PETRA III storage ring at DESY. In total, datasets from 6288 crystals were collected over a period of four weeks. From the 5953 unique compounds in our screen, we obtained 3089 high-quality diffraction datasets to a resolution better than 2.5 Å. Datasets from 1152 compounds were suitable for subsequent automated structure refinement followed by cluster analysis(*9*) and pan dataset density analysis (PanDDA)(*10*). In total, 43 compounds were found that bound to M^pro^. Seven of these compounds had maleate as a counterion and in these structures maleate was found in the active site but not the compounds themselves, resulting in 37 unique binders. A summary of these, together with additional experimental information, is provided in tables S1 and S2. The binding mode could be unambiguously determined for 29 molecules. The majority of hits were found in the active site of the enzyme. Six of 16 active-site binders covalently bind as thioethers to Cys145, one compound binds covalently as a thiohemiacetal to Cys145, one is coordinated through a zinc ion and eight bind non-covalently. The remaining 13 compounds bind outside the active site at various locations (Fig. 1B).

Out of the 43 hits from our X-ray screen, 39 compounds were tested for their antiviral activity against SARS-CoV-2 in cell assays. Ten compounds reduced viral RNA replication by at least two orders of magnitude in Vero E6 cells (Fig. S1). Further evaluation to determine the effective concentrations that reduced not only vRNA but also SARS-CoV-2 infectious particles by 50% (EC_50_) (Fig. 2) showed that six compounds exhibit either selectivity indexes (SI = CC_50_ / EC_50_) greater than five or hundredfold viral reduction with no cytotoxicity in the tested concentration range (table S3).

**Fig. 2.**
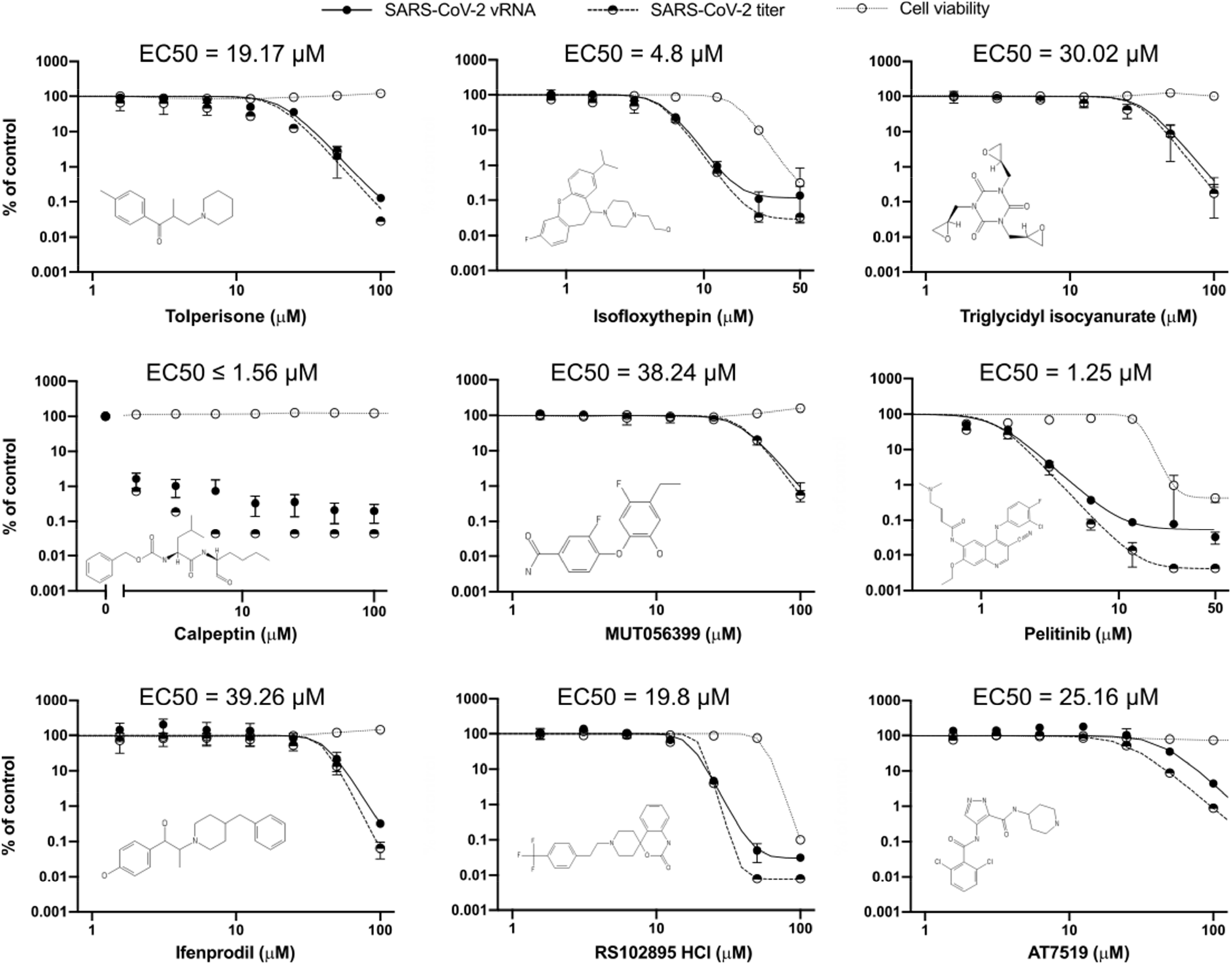
Effect of selected compounds on SARS-CoV-2 replication in Vero E6 cells. The vRNA yield (solid circles), viral titers (half-solid circles), and cell viability (empty circles) were determined by RT-qPCR, immunofocus assays, and the CCK-8 method, respectively. EC_50_ for the viral titers reduction are shown. Values were calculated from three independent replicates in one experiment. Individual data points represent mean ± SD.

In the following we focus on a more detailed description of the most relevant compounds. The compounds are grouped according to their different binding sites. All other compounds are described in more detail in the supplementary text.

**Tolperisone**, HEAT and isofloxythepin bind covalently to the active site. Tolperisone is antivirally active (EC_50_ = 19.17 µM) and shows no cytotoxicity at 100 µ M (Fig. 2), whereas HEAT and isofloxythepin show activity but unfavorable cytotoxicity. For all three compounds only breakdown products are observed in the active site. Tolperisone and HEAT are β-ketoamines, but we only observe the part of the drug containing the activating ketone, while the remaining part with the amine group is missing in the electron-density maps. The breakdown product of the parent drug is observed to bind as Michael-acceptor to the thiol of Cys145. Similarly, the aromatic ring system of both tolperisone (Fig. 3A) and HEAT (Fig. 3B) protrudes into the S1 pocket and forms van der Waals contacts with the backbone of Phe140 and Leu141 and the side chain of Glu166. In addition, the keto group accepts a hydrogen bond from the imidazole side chain of His163. Tolperisone and HEAT bind exclusively in the (S)-configuration. Interestingly, for HEAT, this binding mode was confirmed independently by mass spectrometry (Fig. S2 and table S3). A similar observation has been reported for binding of β-ketoamines to type-1 methionine aminopeptidases, where the parent compound decomposes into an amine and an α,β-unsaturated ketone which subsequently binds to the thiol of the catalytic cysteine(*11*). This is a typical situation for a pro-drug(*12*). Tolperisone is in use as a skeletal muscle relaxant(*13*). Isofloxythepin binds similarly as a fragment to Cys145 (Fig. 3C).

**Fig. 3.**
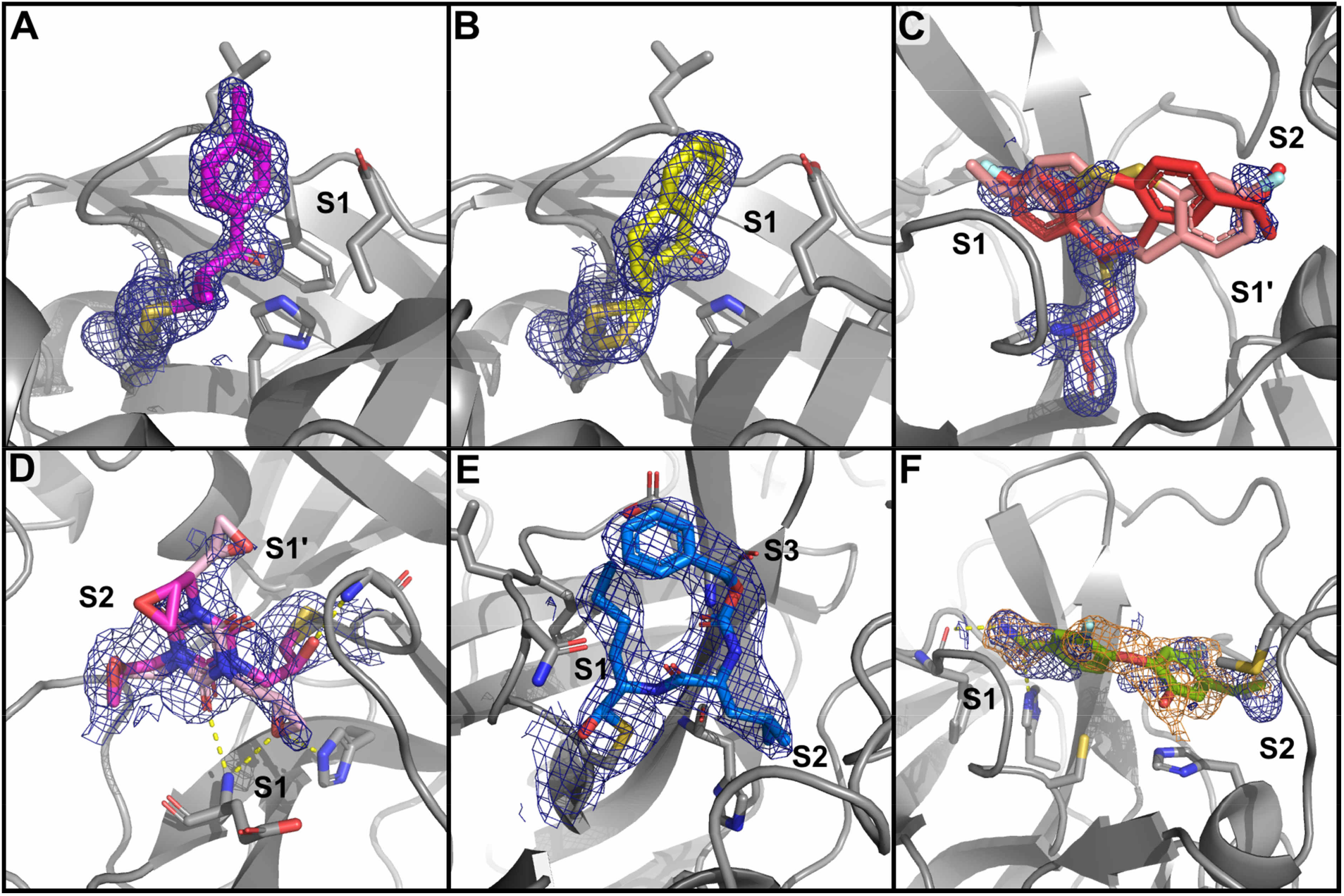
Detailed view of covalent and non-covalent binders in the active site of M^pro^. Bound compounds are depicted as colored sticks while M^pro^ is shown as a grey cartoon representation with selected interacting residues as sticks. Hydrogen bonds are depicted by dashed lines. The blue mesh represents 2Fo-Fc electron-density maps carved at 1.6 Å around the compounds (rmsd = 1 except for **E** and **F**, which are shown at rmsd = 0.7). For **E** the PanDDA event map is additionally shown in orange (rmsd = 1). **A**, tolperisone; **B**, HEAT; **C**, isofloxythepin; **D**, triglycidyl isocyanurate; **E**, calpeptin; **F**, MUT056399. Additional information is provided in table S1.

**Triglycidyl isocyanurate** shows antiviral activity and adopts a covalent and non-covalent binding mode to the M^pro^ active site. In both modes, the compound’s central ring sits on top of the catalytic dyad (His41, Cys145) and its three epoxypropyl substituents reach into subsites S1’, S1 and S2. The non-covalent binding mode is stabilized by hydrogen bonds to the main chain of Gly143 and Gly166, and to the side chain of His163. In the covalently bound form, one oxirane ring is opened by nucleophilic attack of Cys145 forming a thioether (Fig. 3D). The use of epoxides as warheads for inhibition of M^pro^ offers another avenue for covalent inhibitors, whereas epoxysuccinyl warheads have been extensively used in biochemistry, cell biology and later in clinical studies(*14*). Triglycidyl isocyanurate (teroxirone, Henkel’s agent) has been tested as antitumor agent(*15*).

**Calpeptin** shows the highest antiviral activity in the screen, with an EC_50_ value in the lower µM range. It binds covalently via its aldehyde group to Cys145, forming a thiohemiacetal. This peptidomimetic inhibitor occupies substrate pockets S1 to S3, highly similar to inhibitor GC-376(*16, 17*), calpain inhibitors(*18*) and other peptidomimetic inhibitors such as N3(*2*) and the α-ketoamide 13b(*1*). The peptidomimetic backbone forms hydrogen bonds to the main chain of His164 and Glu166, whereas the norleucine side chain is in van der Waals contacts with the backbone of Phe140, Leu141 and Asn142 (Fig. 3E). Calpeptin has known activity against SARS-CoV-2(*16*). The structure is highly similar to leupeptin, which served as positive control in our screen (Fig. S3B). *In silico* docking experiments verified the peptidomimetic compound Calpeptin as a likely M^pro^ binding molecule (table S4).

**MUT056399** is an active-site binding compound without a covalent bond to Cys145 and reduced viral replication. The diphenyl ether core of MUT056399 blocks access to the catalytic site consisting of Cys145 and His41. The terminal carboxamide group occupies pocket S1 and forms hydrogen bonds to the side chain of His163 and the backbone of Phe140 (Fig. 3F). The other part of the molecule reaches deep into pocket S2, which is enlarged by a shift of the side chain of Met49 out of the substrate binding pocket. MUT056399 was developed as an antibacterial agent against multidrug-resistant *Staphylococcus aureus* strains(*19*).

In general, the enzymatic activity of M^pro^ relies on the architecture of the active site, which critically depends on the dimerization of the enzyme and the correct orientation of the subdomains to each other. In addition to the active site, as the most obvious target for drug development, we discovered two allosteric binding sites of M^pro^ which have previously not been reported.

Five compounds of our X-ray screen bind in a hydrophobic pocket in the C-terminal dimerization domain (Fig. 4A-C), located close to the oxyanion hole in pocket S1 of the substrate-binding site. Two of these show antiviral activity in combination with low cytotoxicity (Fig. 2). Another compound with slightly lower antiviral activity binds in between the catalytic and dimerization domains of M^pro^.

**Fig. 4.**
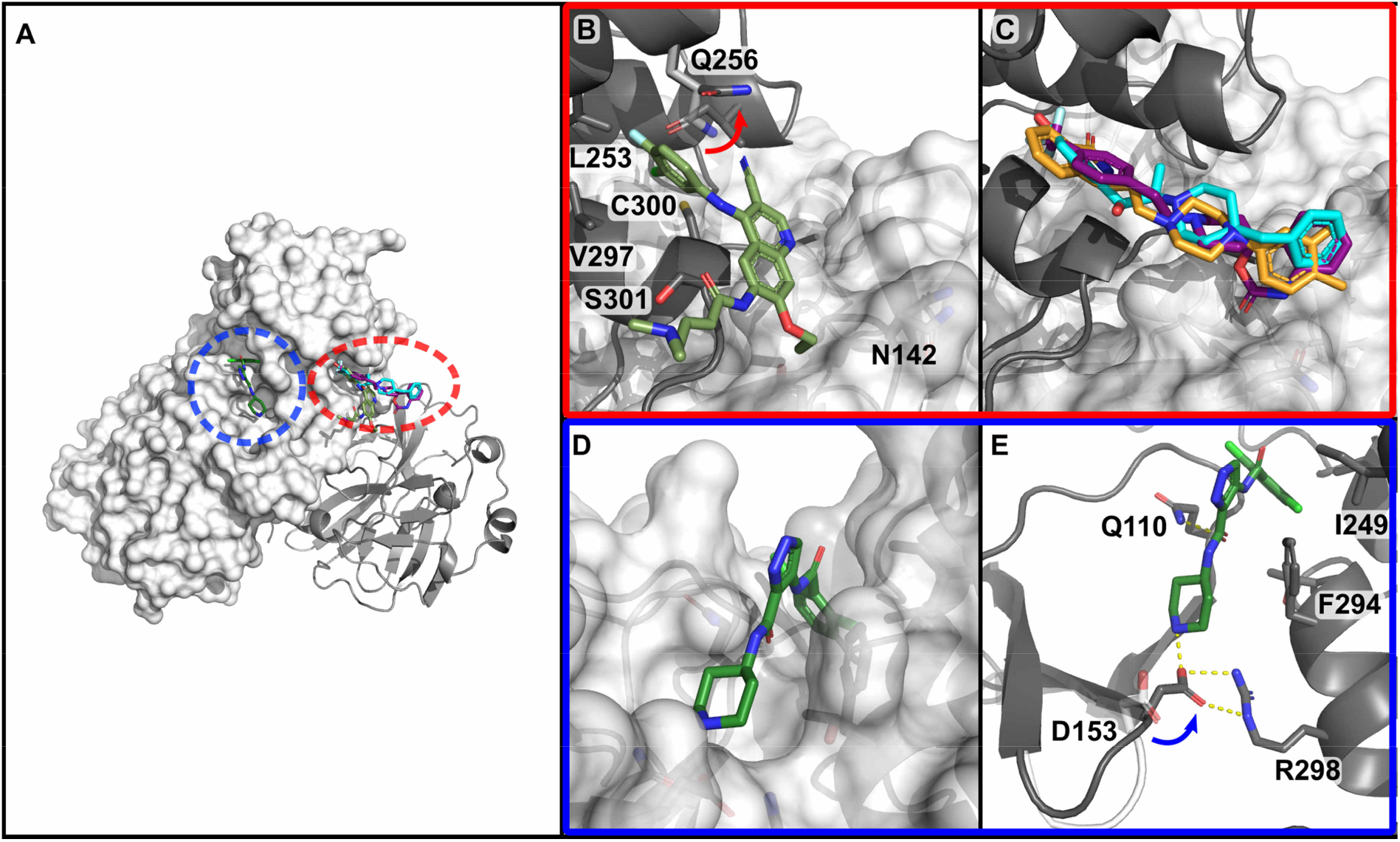
Screening hits at allosteric sites of M^pro^. **A**, View of the allosteric sites of M^pro^. One site is located within the dimerization domain of M^pro^ proximal to the active site (red circle). The other site is located in between the catalytic domains and the dimerization domain in a deep groove (blue). **B**, Close up view of the binding site in the dimerization domain, close to the active site of second protomer in the native dimer. Residues forming the hydrophobic pocket are indicated. Pelitinib (dark green) binds to the C-terminal α-helix at Ser301 and pushes against Asn142 and the β-turn of the pocket S1. Movement of side chain of Gln256 compared to the native M^pro^ structure is indicated by red arrow. **C**, RS-102895 (purple), ifenprodil (cyan) and PD-168568 (orange) occupy the same binding pocket as pelitinib. **D**, AT7519 occupies a deep cleft between the catalytic and dimerization domain of M^pro^. **E**, M^pro^ residues interacting with the compound AT7519 are depicted as sticks, hydrogen bonds and salt bridges are indicated by dashed lines. Loop region containig Asn153 in the native M^pro^ structure is shown in white and movement of this residue is indicate by blue arrow. Additional information is provided in table S1.

Central to the first allosteric binding site is a hydrophobic pocket formed by Ile213, Leu253, Gln256, Val297 and Cys300 within the C-terminal dimerization domain (Fig. 4B). **Pelitinib, ifenprodil**, RS-102895, PD-168568 and tofogliflozin all employ this site by inserting an aromatic moiety into this pocket.

**Pelitinib** shows the second highest antiviral activity in our screen, with an EC_50_ value of 1.25 µM. Its halogenated benzene ring binds to the hydrophobic groove (Fig. 4B). The central 3-cyanoquinoline moiety interacts with the end of the C-terminal helix (Ser301). The ethyl ether substituent pushes against Tyr118 and Asn142 (from loop 141-144 of the S1 pocket) of the opposing protomer within the native dimer. Previous work on M^pro^ of SARS-CoV demonstrated that the integrity of this pocket is crucial for enzyme activity(*20*). Pelitinib is known as an amine-catalyzed Michael acceptor(*21*), developed to bind to a cysteine in the active site of a tyrosine kinase. But from its observed binding position it is impossible for it to reach into the active site and no evidence for covalent binding to Cys145 is found in the electron-density maps. Pelitinib is an irreversible epidermal growth factor receptor inhibitor and developed as an anticancer agent(*22*).

**Ifenprodil**, RS-102895 and PD-168568 all exhibit an elongated structure, consisting of two aromatic ring systems separated by a linker containing a piperidine or piperazine ring (Fig 4C). All three compounds have a distance of at least 12 Å between the terminal aromatic rings. Thus, this binding mode is unlikely to be identified through fragment screening. The hydrophobic pocket in the helical domain is covered by the side chain of Gln256. In our complex structures, this side chain adopts a different conformation. One of the terminal aromatic ring systems is inserted into the hydrophobic groove in the dimerization domain. The linker moiety stretches across the native dimer interface and the second aromatic ring is positioned close to Asn142, adjacent to the active site loop where residues 141-144 contribute to the pocket S1. In particular, in the case of RS-102895, two hydrogen bonds are formed to the side and main chains of Asn142. In contrast to ifenprodil, RS-102895 and PD-168568 do not exhibit selective antiviral activity (SI<5). All three compounds are GPCR antagonists. Ifenprodil antagonizes N-methyl-D-aspartate receptors(*23*), RS-102895 inhibits C-C chemokine receptor 2(*24*), and PD-168568 dopamine-receptors(*25*). Of note, ifenprodil is currently in phase IIb/III clinical trials for the treatment of COVID-19 based on the observation that it reduces mortality of lethal infection of H5N1 influenza in mice, likely through reduced inflammatory cytokine expression(*26*). Our crystal structure and antiviral tests suggest an additional mode of action beyond this anti-inflammatory effect.

The second allosteric site is formed by the deep groove between the catalytic domains and the dimerization domain (Fig. 4A and D). AT7519 is the only compound in our screen that we identified bound to this site. The chlorinated benzene ring is engaged in various van der Waals interactions to loop 107-110, Val202, and Pro293 (Fig. 4E). The central pyrazole has van der Waals contacts to Ile249, Phe294 and its adjacent carbonyl group forms a hydrogen bond to the side chain of Gln110. The terminal piperidine forms hydrogen bonds to the carboxylate of Asp153. This results in a displacement of loop 153-155, slightly narrowing the binding groove. The Cα-atom of Tyr154 moves by 2.8 Å, accompanied by a conformational change of Asp153. This allows hydrogen bonding to the compound and the formation of a salt-bridge to Arg298. In turn, Arg298 is crucial for dimerization(*27*). The mutation Arg298Ala causes a reorientation of the dimerization domain relative to catalytic domain, leading to changes in the oxyanion hole and destabilization of the S1 pocket by the N-terminus. AT7519 was evaluated for treatment of human cancers(*28*) and shows weak antiviral activity but a poor selectivity index against SARS-CoV-2 (Fig. 2).

Our X-ray screen revealed six compounds with previously unreported antiviral activity against SARS-CoV-2. Two of them, calpeptin and pelitinib, show strong antiviral activity combined with low cytotoxicity and are suitable for preclinical evaluation. The remaining compounds are valuable lead structures for further drug development. A general advantage of using drug-repurposing libraries for such a screening is the proven bioactivity of the compounds and key properties such as cell-permeability are usually known(*29*).

The most active compound, calpeptin binds in the active site in the same way as other members of the large class of peptide-based inhibitors that bind as thiohemi-acetals or -ketals to M^pro^ (*30*). However, in addition to this peptidomimetic inhibitor, we discovered several non-peptidic inhibitors. Those compounds binding to the active site of M^pro^ contained new Michael acceptors based on β-ketoamines (tolperisone and HEAT). These lead to the formation of thioethers and have not previously been described as prodrugs for viral proteases. We also identified a non-covalent binder, MUT056399, blocking the active site. Besides this common orthosteric inhibition, we discovered compounds that inhibit the enzyme through binding at two previously unreported allosteric sites of M^pro^.

The first allosteric site (dimerization domain) is in direct vicinity of the S1 pocket of the adjacent monomer within the native dimer. The potential for antiviral inhibition through this site is demonstrated by ifenprodil and pelitinib. A comparison of coronavirus M^pro^ sequences shows that the compound binding residues of this allosteric site are conserved (Fig. S4). Consequently, potential drugs targeting this allosteric binding site can be assumed to be robust against mutational variations and might also be effective against other coronaviruses.

The potential of the second allosteric site, connecting the dimerization and catalytic domain, as a druggable target is demonstrated by the observed weak antiviral activity of AT7519. Recently, the potential of allosteric inhibition of M^pro^ through modulation of its dimerization has been demonstrated by mass spectrometry(*31*).

## Materials and Methods

### Protein production and purification

The protein was overexpressed in *E. coli* and purified for subsequent crystallization according to previously published protocols and plasmid constructs(*1*). Lysis was carried out in 20 mM HEPES buffer supplemented with 150 mM NaCl using ultrasound for cell disruption. After separation of the cell fragments and the dissolved protein, a subsequent nickel NTA column was used to extract the M^pro^-histidine-tag fusion. The cleavage of the histidine tag was achieved by a 3C protease during an overnight dialysis step. The histidine tag and the 3C protease were removed using a nickel NTA column, and as a final step a gel filtration was performed with an S200 Superdex column.

### Crystallization experiments

Co-crystallization with the compounds was achieved mixing 0.23 μL of protein solution (6.25 mg/mL) in 20 mM HEPES buffer (pH 7.8) containing 1 mM DTT/TCEP (respectively), 1 mM EDTA, and 150 mM NaCl with 0.22 μL of reservoir solution consisting of 100 mM MIB, pH 7.5, containing 25% w/w PEG 1500 and 5% (v/v) DMSO, and 0.05 μL of a micro-seed crystal suspension using an Oryx4 pipetting robot (Douglas Instruments). This growth solution was equilibrated by sitting drop vapor diffusion against 40 μL reservoir solution. Prior to crystallization 125 nL droplets of 10 mM compound solutions from the two libraries in DMSO were applied to the wells of SwissCI 96-well plates (2-well or 3-well low profile, respectively) and subsequently dried in vacuum. Taking the crystallization drop volume into account this resulted in a final compound concentration of 2.5 mM and a molar ratio of ∼13.6 of compound to protein. To obtain well-diffracting crystals in a reproducible way micro-seeding was applied for crystal growth(*32*). Crystals appeared within a few hours and reached their final size (∼200×100×10 µm^3^) after 2 - 3 days. Crystals were manually harvested and flash-frozen in liquid nitrogen for subsequent X-ray diffraction data collection. We aimed at harvesting two crystals per crystallization condition as a compromise between through-put and increasing the probability to collect data from well diffracting crystals.

### Data collection

Data collection was performed at beamlines P11, P13 and P14 at the PETRA III storage ring at DESY in Hamburg within a period of four weeks. Exclusive use of DESY beamline P11 was generously granted by the DESY directorate for the project.

### Data processing and structure refinement

An automatic data processing and structure refinement pipeline “xia2pipe” as written specifically to support this project. Raw diffraction images from the PETRA III beamlines were processed using three crystallographic integration software packages: XDS(*33*), autoPROC(*34*) followed by staraniso(*35*), and DIALS via xia2(*36, 37*). Diffraction data quality indicators for all datasets and the 43 hits are summarized in Fig. S5. In total, 7857 unique crystals were harvested and frozen, of which 7258 were studied by X-ray diffraction at PETRA-III. Of these, 5934 produced diffraction data consistent with a protein lattice and were labeled as “successful” experiments. In some cases, multiple datasets were collected on a single crystal, so in total 8304 diffraction experiments were conducted with 6831 successful protein diffraction datasets obtained. As processed by DIALS, these 6831 datasets had an average resolution of 2.12 Å (criterion: CC1/2 > 0.5), CC1/2 of 0.97, and Wilson B of 27.8 Å^2^ (Fig. S5). Crystallographic data of all structures submitted to the PDB are summarized in table S2.

For clustering and hit identification, all datasets were integrated and merged to a resolution of 1.7 Å. In order to reduce the influence of noise for lower resolution datasets, the following processing was applied to standardize the Wilson plot for each dataset: the datasets were split into equally sized bins, each covering 1000 reflections, and a linear fit was applied to the logarithm of the average intensities in each shell. The residual between the data and the Wilson fit was calculated, considering sequentially one additional bin from low to high resolution until the residual exceeded 10%, if applicable. The intensities in all higher resolution bins beyond this point were scaled to fit the calculated Wilson B factor.

The results of each dataset were then automatically refined using Phenix(*38*). Refinement began by choosing one of two manually refined starting models (differing in their unit cell, table S2), selecting the starting model with the closest unit cell parameters, then proceeding in four steps: (a) rigid body and ADP refinement, (b) simulated annealing, ADP, and reciprocal space refinement, (c) real-space refinement, and (d) a final round of reciprocal space refinement as well as TLS refinement, with each residue pre-set as a TLS group. This procedure was hand-tuned on 5 test datasets; the procedure and parameters were manually adjusted to minimize Rfree until deemed satisfactory for the continuation of the project. All processing and refinement results were logged in a database, which enabled comparison between methods and improvement over time. All code and parameters needed to reproduce this pipeline are available online(*39*).

### Hitfinding: cluster4x and PanDDA Analysis

The resulting model structure Cα positions were then ingested into cluster4x(*40*), which briefly (a) computes a correlation coefficient between each structure over the position of all C_α_ atoms, (b) performs PCA the resulting correlation matrix, (c) presents 3 chosen principal components to a human, who then manually annotates clusters. Clusters were ordered chronologically and separated into groups of 1500 and subsequently clustered into groups of approximately 60-120 datasets based on a combination of reciprocal and Cα-atom differences using cluster4x. In an earlier version of the software, structure factor amplitudes were used for clustering instead of refined Cα positions, and both methods were applied for hitfinding. The resulting clusters were then analyzed via PanDDA(*10*) using default parameters. The resulting PanDDA analyses were manually inspected for hits which were recorded.

### Manual structure refinement

Identified hits were further refined by alternating rounds of refinement using refmac(*41*), phenix.refine(*38*) or MAIN(*42*), interspersed with manual model building in COOT(*43*).

### *In sillico* screening of compound libraries

To enable a preselection of potentially promising compounds to support the experimental X-ray screening effort and to get an idea about the most promising compounds, we pursued a virtual screening workflow consisting of the selection of a representative ensemble of binding site conformations, non-covalent molecular docking and rescoring. We performed this study with 5,575 compounds of the Fraunhofer IME Repurposing Collection. UNICON(*44*) was applied to prepare the library compounds. To consider binding site flexibility, we used multiple receptor structures. We applied SIENA(*45*) to extract five representative binding site conformations for the active site of M^pro^. We chose the structures with the PDB IDs 5RFH, 5RFO, 6W63, 6Y2G and 6YB7 The SIENA-derived aligned structures were used and the proteins were preprocessed using Protoss(*46*) to determine protonation states, tautomeric forms, and hydrogen orientations. The binding site was defined based on the active site ligand of the structure with the PDB ID 6Y2G (ligand ID O6K). A 12.5 Å radius of all ligand atoms was chosen as binding site definition. The new docking and scoring method JAMDA was applied with default settings for the five selected binding sites(*47*). Subsequently, HYDE(*48*) was used for a rescoring of all predicted poses of the library compounds. The 200 highest ranked compounds of all 5,575 compounds according to the HYDE score were extracted. For 70 of these compounds, well-diffracting crystals were obtained in the X-ray screening. Intriguingly, only calpeptin, a known cysteine protease inhibitor, could be co-crystallized and was found on rank 3 (table S5).

### Mass Spectrometry

M^pro^ was prepared for native MS measurements by buffer-exchange into ESI compatible solutions (250 μM, 300 mM NH4OAc, 1 mM DTT, pH 7.5) by five cycles of centrifugal filtration (Vivaspin 500 columns, 30,000 MWCO, Sartorius). Inhibitors were dissolved to 1 mM in DMSO. Then Inhibitors and M^pro^ were mixed to final concentrations of 50 µM and 10 µM, respectively, and incubated for 16 h at 4 °C. For putative covalent ligands, compounds were incubated at 1 mM with 100 µM M^pro^ in 20 mM Tris, 150 mM NaCl, 1 mM TCEP, pH 7.8, for 16 h prior to buffer exchange. Buffer exchange was carried out as described above and samples were diluted tenfold prior to native MS measurements. All samples were prepared in triplicate. Nano ESI capillaries were pulled in-house from borosilicate capillaries (1.2 mm outer diameter, 0.68 mm inner diameter, filament, World Precision Instruments) with a micropipette puller (P-1000, Sutter instruments) using a squared box filament (2.5 × 2.5 mm^2^, Sutter Instruments) in a two-step program. Subsequently capillaries were gold-coated using a sputter coater (CCU-010, safematic) with 5.0 × 10-2 mbar, 30.0 mA, 100 s, 3 runs to vacuum limit 3.0 × 10-2 mbar argon. Native MS was performed using an electrospray quadrupole time-of-flight (ESI-Q-TOF) instrument (Q-TOF2, Micromass/Waters, MS Vision) modified for higher masses(*49*). Samples were ionized in positive ion mode with voltages of 1300 V applied at the capillary and of 130 V at the cone. The pressure in the source region was kept at 10 mbar throughout all native MS experiments. For desolvation and dissociation, the pressure in the collision cell was adjusted to 1.5 × 10^−2^ mbar argon. Native-like spectra were obtained at an accelerating voltage of 30 V. To calibrate raw data, CsI (25 mg/ml) spectra were acquired. Calibration and data analysis were carried out with MassLynx 4.1 (Waters) software. In order to determine each inhibitor binding to M^pro^, peak intensities of zero, one or two bound ligands were analyzed from three independently recorded mass spectra at 30 V acceleration voltage. Results are shown in table S4.

### Antiviral assays

#### Compounds

All compounds were diluted to a 50 mM concentration in 100% DMSO and stored at -80°C.

#### Cytotoxicity assays

Vero E6 cells (ATCC CRL-1586) were seeded at 3.5 × 10^4^ cells/well in 96-well plates. After 24 h, the cell culture media was changed and 2-fold serial dilutions of the compounds were added. Cell viability under 42 h compound treatment was determined via the Cell Counting Kit-8 (CCK-8, Sigma-Aldrich #96992) following the manufacturer’s instructions.

#### Antiviral activity assays

Vero E6 cells (ATCC CRL-1586) seeded at 3.5 × 10^4^ cells/well in 96-well plates were pretreated 24 h later with twofold serial dilutions of the compounds. After 1 h incubation with the compounds, SARS-CoV-2 (strain SARS-CoV-2/human/DEU/HH-1/2020) was subsequently added at a MOI of 0.01 and allowed absorption for 1 h. The viral inoculum was removed, cells were washed with PBS without Mg^2+^ / Ca^2+^ and fresh media containing the compounds (final DMSO concentration 0.5% (v/v)) was added to the cells. Cell culture supernatant was harvest 42 hpi and stored at -80°C. Viral RNA was purified from the cell culture supernatant using the QIAamp Viral RNA Mini Kit (QIAGEN #52906) in accordance with the manufacturer’s instructions. Quantification of vRNA was carried out by the interpolation of RT-qPCR (RealStar SARS-CoV-2 RT-PCR Kit, Altona Diagnostics #821005) results onto a standard curve generated with serial dilutions of a template of known concentration. Titers of infectious virus particles were measured via immunofocus assay. Briefly, Vero E6 cells (ATCC CRL-1586) seeded at 3.5 × 10^4^ cells/well in 96-well plates were inoculated with 50 µl of serial tenfold dilutions of cell culture supernatant from treated cells. The inoculum was removed after 1 h and replaced by a 1.5% methylcellulose-DMEM-5% FBS overlay. Following incubation for 24 h, cells were inactivated and fixed with 4.5% formaldehyde. Infected cells were detected using an antibody against SARS-CoV-2 NP (ThermoFischer, PA5-81794). Foci were counted using an AID ELISpot reader from Mabtech. The cytotoxic concentrations that reduced cell growth by 50% (CC_50_) and the effective concentrations that reduced infectious particles or vRNA by 50% (EC_50_) were calculated by fitting the data to the sigmoidal function using GraphPad Prism version 8.00 (GraphPad Software, La Jolla California USA, www.graphpad.com).

## Supplementary Text

In the following, we discuss those compounds that did not show significant antiviral activity but for which we could determine the binding pose based on the crystal structures.

### Active site, covalent

**Isofloxythepin** binds as breakdown product (Fig. 3C). Here, the piperazine group is not found in the crystal structure but the dibenzothiepine moiety is observed in the active site, bound as a thioether to Cys145. The tricyclic system stretches from the S1 across to the S1’ pocket. According to the electron-density maps, two orientations of the molecule are possible, with either the fluorine or the isopropyl group placed inside the S1 pocket. Degradation of the drug with piperazine as the leaving group has been previously reported(*50*) and was confirmed by mass spectrometry (Fig. S2). Isofloxythepin is an antagonist of dopamine receptors D1 and D222 and has been tested as a neuroleptic in phase II clinical trials.

**Leupeptin** is a well-known cysteine protease inhibitor and was therefore included in our screening effort as a positive control(*51*). Structurally, it is highly similar to calpeptin. Indeed this peptidomimetic inhibitor also forms a thiohemiacetal and occupies the substrate pocket, much like calpeptin (Fig. S3B and 3E). The binding mode is identical to the recently released room-temperature structure of M^pro^ with leupeptin (PDB-ID 6XCH).

**Maleate** was observed covalently bound in seven structures during hit finding. In all cases maleate served as the counter ion of the applied compound. In these crystal structures the maleate, rather than the applied compound, forms a thioether with the thiol of Cys145, modifying it to succinyl-cysteine. The thiol of Cys145 undergoes a Michael-type nucleophilic attack on the C2 of maleate. A similar adduct has been described for maleate isomerase(*52*) as an intermediate structure in the isomerization reaction. The covalent adduct is further stabilized by hydrogen bonds to the backbone amide of Gly143 and Cys145 to the carboxylate group (C1) of succinate. The terminal carboxylate (C4) is positioned by hydrogen bonds to the side chain of Asn142 and a water-bridged hydrogen bond to the side chain of His163 (Fig. S3A).

**TH-302 (Evofosfamide)** is covalently linked to Cys145 through nucleophilic substitution of the bromine, leading to thioether formation (Fig. S3C). The other bromine-alkane chain occupies the S1 pocket while the nitro-imidazole stretches into pocket S2. The substitution of chlorine or hydroxyl for bromines in TH-302 has been demonstrated in cell culture(*53*). Our mass spectrometry analysis suggested the loss of a bromine atom (Fig. S2C).

**Triglycidyl isocyanurate** shows antiviral activity and adopts a covalent and non-covalent two binding modes to the M^pro^ active site, one covalent and one non-covalent. In both modes, the compound’s central ring sits on top of the catalytic dyad (His41, Cys145) and its three epoxypropyl substituents reach into subsites S1’, S1 and S2. The non-covalent binding mode is stabilized by hydrogen bonds to the main chain of Gly143 and Gly166, and to the side chain of His163. In the covalently bound form, one oxirane ring is opened by nucleophilic attack of Cys145 forming a thioether (Fig. 3D). The use of epoxides as warheads for inhibition of M^pro^ offers another avenue for covalent inhibitors, whereas epoxysuccinyl warheads have been extensively used in biochemistry, cell biology and later in clinical studies23. Triglycidyl isocyanurate (teroxirone, Henkel’s agent) has been tested as antitumor agent24.

**Zinc pyrithione** was already demonstrated to have inhibitory activity against SARS-CoV-1 M^pro^ (*54*).The pyrithione chelates the Zn^2+^ ion which coordinates the thiolate and imidazole of the catalytic dyad residues Cys145 and His41 (Fig. S3D). The remaining part of the ionophore protrudes out of the active site. This tetrahedral binding mode of zinc has previously been described for other zinc-coordinating compounds in complex with HCoV-229E M^pro^(*55*). Interestingly, antiviral effects against a range of corona- and non-coronaviruses have already been ascribed to zinc pyrithione, although its effect had been attributed to inhibition of RNA-dependent polymerase(*56*). Zinc pyrithione exhibits both antifungal and antimicrobial properties and is known in treatment of seborrheic dermatitis.

### Active site, non-covalent

**Adrafinil** mainly binds mainly through van der Waals interactions to M^pro^. In particular, its two phenyl rings are inserted into pockets S1’ and S2 (Fig. S3E). A hydrogen bond is formed between the backbone amide of Cys145 and the hydroxylamine group. The side chain of Met49 is wedged between the two phenyl rings.

**Fusidic acid** interacts with M^pro^ mainly through hydrophobic interactions, especially through the alkene chain within pocket S2 and the tetracyclic moiety packing against Ser46 (Fig. S3F). Moreover, the carboxylate group forms indirect hydrogen bonds, mediated via two water molecules, to the main chain of Thr26, Gly143 and Cys145. In addition, the same carboxylate group forms a hydrogen bond to an imidazole molecule from the crystallization conditions. This imidazole occupies pocket S1’ and mediates hydrogen bonds to the backbone of His41 and Cys44. These indirect interactions offer opportunities for optimization of compounds binding to M^pro^. Fusidic acid is a well-known bacteriostatic compound, with a steroid core structure.

**LSN-2463359** binds mainly to M^pro^ by interaction of the pyridine ring with the S1 pocket (Fig. S3G). Besides van der Waals interactions with the β-turn Phe140-Ser144, contributing to the pocket, it also forms a hydrogen bond to the side chain of His163.

**SEN1269** binds only to the active site of one protomer in the native dimer. This causes a break in the crystallographic symmetry, leading to a different crystallographic space group (table S2). The central pyrazine ring forms a hydrogen bond to Gln189 (Fig. S3H). The terminal dimethylaniline moiety sits deep in pocket S2 which is enlarged by an outwards movement of the short α-helix Ser46-Leu50 by 1.7 Å (Ser46 Cα-atom) compared to the native structure. This includes a complete reorientation of the side chain of Met49 which now points outside of the S2 pocket. Additionally, the C-terminus of a crystallographic neighboring M^pro^ protomer is trapped between SEN1269 and part of the S1 pocket, including a hydrogen bond between Asn142 and the backbone amide of Phe305 and Gln306 of the C-terminus.

**Tretazicar** binds at the active site entrance at pocket S3/S4 (Fig. S3I). The amide group forms hydrogen bonds to the backbone carbonyl of Glu166, the adjacent nitro group forms hydrogen bonds to the side chain of Gln192 and the backbone amide of Thr190.

**UNC2327** binds to active site of M^pro^ by stacking its benzothiadiazole ring against the loop Glu166-Pro168 that forms the shallow pocket S3 (Fig. S3J). This is stabilized by a hydrogen bond between the benzothiadiazole and the main chain carbonyl of Glu166. The piperine ring and adjacent carbonyl are inserted into pocket S1’ and interact with Thr25 and His41.

### Covalent binder to Cys156

#### Aurothioglucose

In the crystal structure of the aurothioglucose complex, the strong nucleophile Cys145 becomes oxidized to a sulfinic acid. The initial reaction is the disproportionation of Aurothioglucose into Au(0) and a disulfide dimer of thioglucose. This is followed by a cascade of redox reactions of thioglucose, its disulfide and sulfenic acid. A disulfide linkage to thioglucose is only observed at Cys156 on the surface of M^pro^ (Fig. S3K). Here the thioglucose moiety is located between Lys100 and Lys102.

**Glutathione isopropyl ester** binds to the surface-exposed Cys156 via a disulfide linkage (Fig. S3L). Additionally, the ester forms a hydrogen bond to the backbone amide of Tyr101, while the amine of the other arm of the molecule is interacting with the side chain amine of Lys102.

### Surface pockets

**AR-42** binds with its phenyl ring to a small hydrophobic pocket in the dimerization domain formed by residues Gly275, Met276, Leu286 and Leu287 (Fig. S3M). Additionally, the central amide forms a hydrogen bond to the backbone carbonyl of Leu272.

**AZD6482** binds to a pocket on the back of the catalytic domain, away from the native dimer interface (Fig. S3N). The nitrobenzene ring is inserted in a pocket formed by His80, Lys88, Leu89 and Lys90. The central aromatic system and morpholine ring lie flat on the surface of M^pro^. Furthermore, Asn63 forms a hydrogen bond to the keto-group in the pyrimidine ring.

**Climbazole** binds in a shallow surface pocket, wedged between two crystallographic symmetry-related molecules (Fig. S3O). Only van der Waals interactions are observed. One monomer contributes with residues Phe103, Val104, Arg105 and Glu178 to this binding site, while the other monomer contributes Asn228, Asn231, Leu232, Met235 and Pro241.

**Clonidine** also sits in between two crystallographic, symmetry-related molecules and binds through van der Waals interactions (Fig. S3P). Here one protomer mainly forms the binding site, by contributing Asp33, Aps34 and Ala94. The other protomer contributes Lys236, Tyr237 and Asn238. The amine ring of clonidine forms a loose ring stacking interaction to Tyr237, while a hydrogen bond between the backbone carbonyl of Lys236 and the ring connecting amine of clonidine is formed. The side chain of Lys236 is flipped to the side to make room for the chlorine containing ring system.

**Ipidacrine** is in contact with two different M^pro^ protomers (Fig. S3Q). The tricylic ring system is packed against a surface loop, including residues Pro96 and Lys97 as well as Lys12. It also interacts with the end of an α-helix including residues Gln273, Asn274 and Gly275.

**Tegafur** binds to a in a shallow surface pocket generated by residues Asp33, Pro99, Lys100 and Tyr101. The main interaction is through π-stacking of the aromatic ring of Tyr223. The side chain of Lys100 flips away and generates space for the compound (Fig. S3R).

### Allosteric site I

**Tofogliflozin** binds to the same hydrophobic pocket as pelitinib, ifenprodil, RS-102895, and PD-168568 but no antiviral activity was observed at 100 µM, the highest concentration tested. In contrast to the previous four compounds, it does not reach across to the opposing protomer in the native dimer. Its main interaction with M^pro^ is via its isobenzofuran moiety that occupies the hydrophobic pocket (Fig. S3S).

## Acknowledgements

We acknowledge DESY (Hamburg, Germany), a member of the Helmholtz Association HGF, for the provision of experimental facilities. Parts of this research were carried out at PETRA III at beamline P11. Further MX data was collected at beamline P13 and P14 operated by EMBL. We thank the DESY machine group, in particular Mario Wunderlich, Kim Heuck, Arne Brinkmann, Olaf Goldbeck, Jürgen Haar, Torsten Schulz, Gunnar Priebe, Maximilian Holz, Björn Lemcke, Klaus Knaack, Oliver Seebauer, Philipp Willanzheimer, Rolf Jonas, and Nicole Engling. We thank Thomas Dietrich, Simon Geile, Heshmat Noei, and Tim Pakendorf from DESY, and Bianca Di Fabrizio and Sebastian Kühn from BNI for assistance. This research was supported in part through the Maxwell computational resources operated at Deutsches Elektronen-Synchrotron (DESY), Hamburg, Germany. We acknowledge the use of the XBI biological sample preparation laboratory, enabled by the XBI User Consortium. **Funding**: We acknowledge financial support through the EXSCALATE4CoV EU-H2020 Emergency Project (https://www.exscalate4cov.eu), by the Cluster of Excellence ‘Advanced Imaging of Matter’ of the Deutsche Forschungsgemeinschaft (DFG) - EXC 2056 - project ID 390715994, the Federal Ministry of Education and Research (BMBF) via project 05K19GU4, 05K16GUA and 05K20FL1 (“STOP CORONA”), the Joachim-Herz-Stiftung Hamburg via the project Infecto-Physics. CE and MR acknowledge the financial support by Grant-No. HIDSS-0002 DASHH (Data Science in Hamburg - HELMHOLTZ Graduate School for the Structure of Matter). The work of JPZ and MR was supported by the BMBF (SFX2: Hochdurchsatz Serielle-Femtosekunden Kristallographie@EU XFEL (WP2) - Compound Target Screenings of essential SARS-CoV-2 Enzymes and selected Human Corona processing Enzymes FKZ: 05K19GU4). DT is supported by Slovenian Research Agency (ARRS; research program P1-0048, Infrastructural program IO-0048 – both awarded to D.T.). B.S. was supported by an Exploration Grant from the Boehringer Ingelheim Foundation. R.C. is supported by DFG grants INST 187/621-1 and INST 187/686-1.

## Author contributions

SeG, PR, YFG, WB, PG, ARB, RC, DT, AZ, HNC, ARP, CB, AM designed research. SeG, PR, TJL, WH, HNC, ARP, CB, AM wrote manuscript. SeG, PR, JL, FK, SM, WB, ID, BS, HGie, BNB, MB, PLX, NW, HA, NU, SF, BAF, MS, HB, JK, GEPM, ARM, FG, VH, PF, MW, ECS, PM, HT, TB participated in sample preparation. PR performed crystallization experiments. SeG, PR, JL, TJL, OY, SS, AT, MGr, HF, FT, MGa, YG, CFL, SA, AP, GB, DVS, GP, TRS, IB, SP performed X-ray data collection. TJL, HGin, DO, OY, LG, MD, TAW,FS, CR, DM, JZD, IK, CS, RS, HUH, DCFM contributed to X-ray data management. SeG, PR, JL, TJL, HGin, FHMK, WE, DO, AH, VS, JH, JM, JB, JW, CF, MSW, AC, DT, WH, AM performed X-ray data analysis. KL, BK, CU, RC performed and analyzed MS experiments. YFG, BEP, StG performed and analyzed antiviral activity assays. PG, BE, MK, MGA, SN, CG, LZ, XS, KK, AU, JL, RH, performed and analyzed ligand binding studies and protein activity assays. CE, JPZ, MR performed computational binding studies.

## Data and materials availability

The coordinates and structure factors for all described crystal structures of SARS-CoV-2 M^pro^ in complex with compounds are deposited in the PDB with accession codes 6YNQ, 6YT8, 6YVF, 6YZ6, 7A1U, 7ABU, 7ADW, 7AF0, 7AGA, 7AHA, 7AK4, 7AKU, 7AMJ, 7ANS, 7AOL, 7AP6, 7APH, 7AQE, 7AQI, 7AQJ, 7AR5, 7AR6, 7ARF, 7AVD, 7AWR, 7AWS, 7AWU, 7AWW, 7AX6, 7AXM, 7AXO, and 7AY7. Code used in this analysis has been previously published(*40*). The code for forcing adherence to the Wilson distribution is included in the same repository under a GPLv3 license. Any other relevant data are available from the corresponding authors upon reasonable request.

**Correspondence and requests for data and materials should be addressed to**.

---

**Fig. S1.**
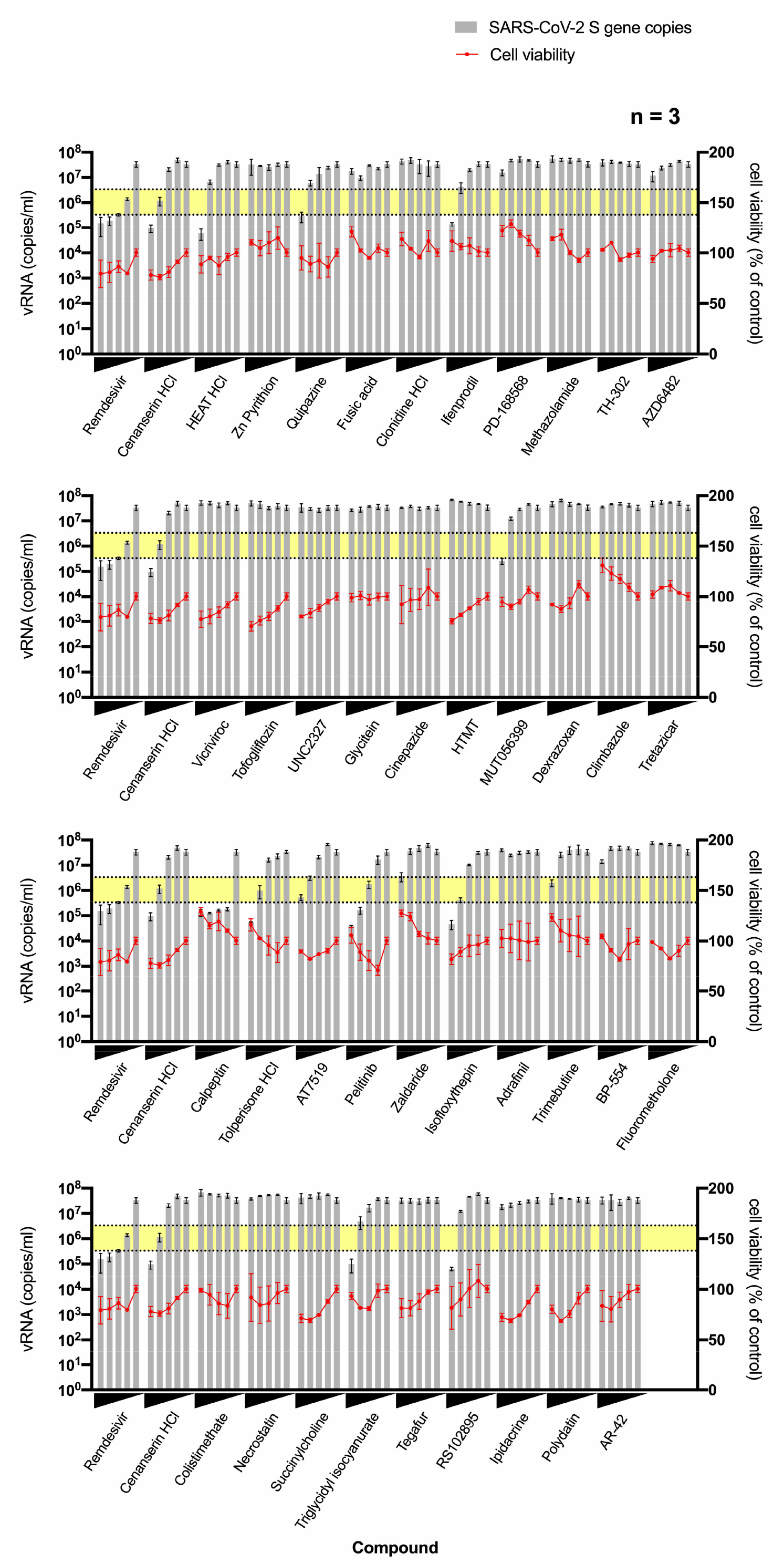
X-ray hit compounds were tested in a non-toxic range for inhibition of SARS-CoV-2 replication in Vero E6 cells. The vRNA yield (gray bars) and cell viability (red circles) were determined by RT-qPCR and the CCK-8 method, respectively. All data are mean ± standard deviation. Upper and lower boundaries of yellow bars represent one and two log reduction in vRNA level. Twofold serial dilutions of compounds were used to treat cells for 42 hours, where 100 µM was used as the highest concentration for all compounds except remdesivir (10 µM), cenanserin HCl (125 µM, HEAT HCl (25 µM), Zn pyrithion (1 µM), pelitinib (12.5 µM), zaldaride (50 µM), isofloxythepin (25 µM) and RS-102895 HCl (50 µM). Control is DMSO without compound.

**Fig. S2.**
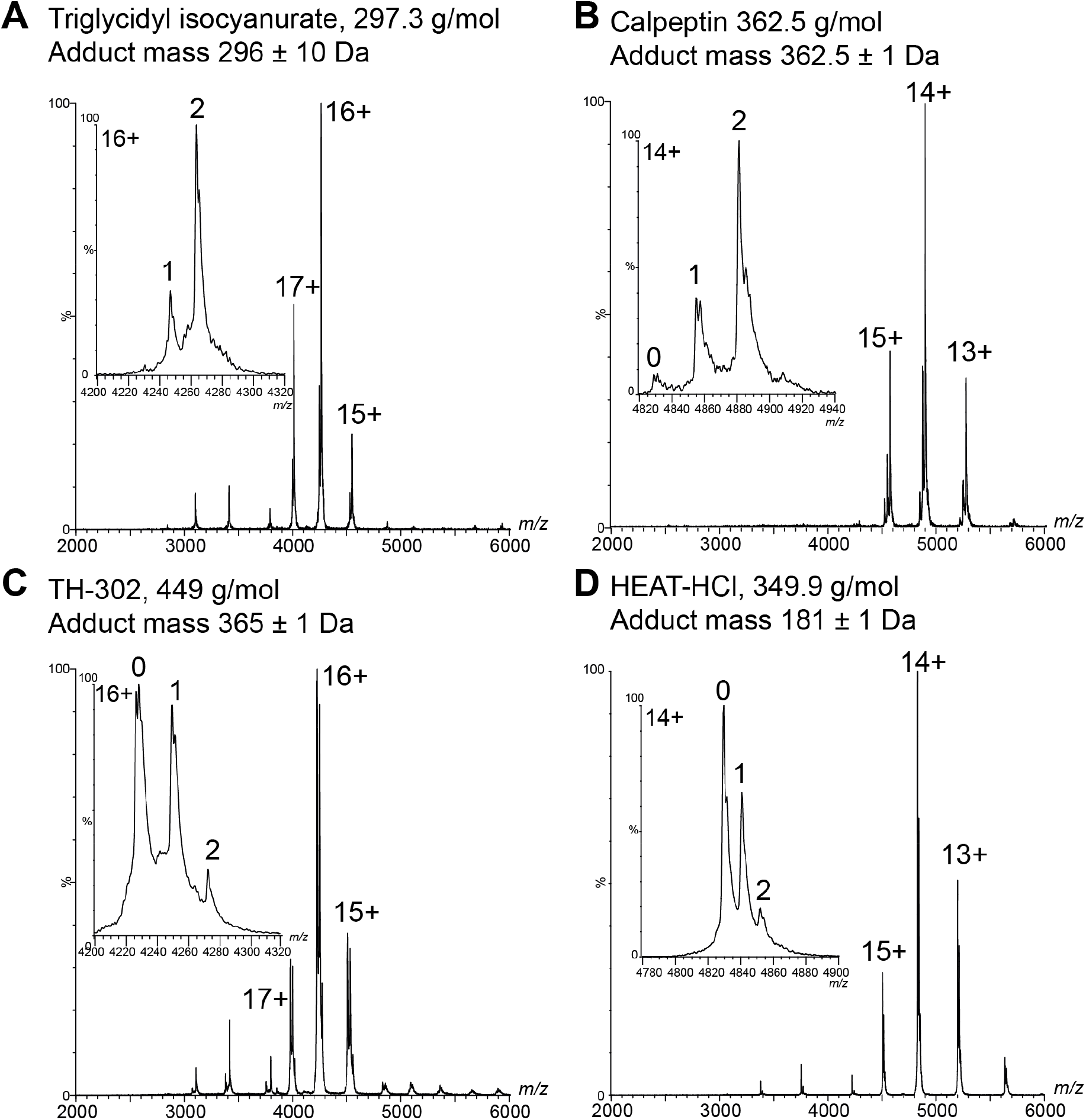
Binding of compounds confirmed by native mass-spectrometry. Main mass spectra of M^pro^ with compounds. **(A)**, Triglycidyl isocyanurate, **(B)** calpeptin, **(C)** TH-302 and **(D)** HEAT-HCl. Insets depict main charge state signals with native M^pro^ (0) binding to one (1) or two (2) compounds, exhibiting the molecular mass of the complete compound (A and B) or a fragment (C and D). Mass spectra were recorded after the inhibitor was washed out (A and C) or in presence of fivefold excess of compound (B and D). Average compound masses are given and charged states are labelled.

**Fig. S3.**
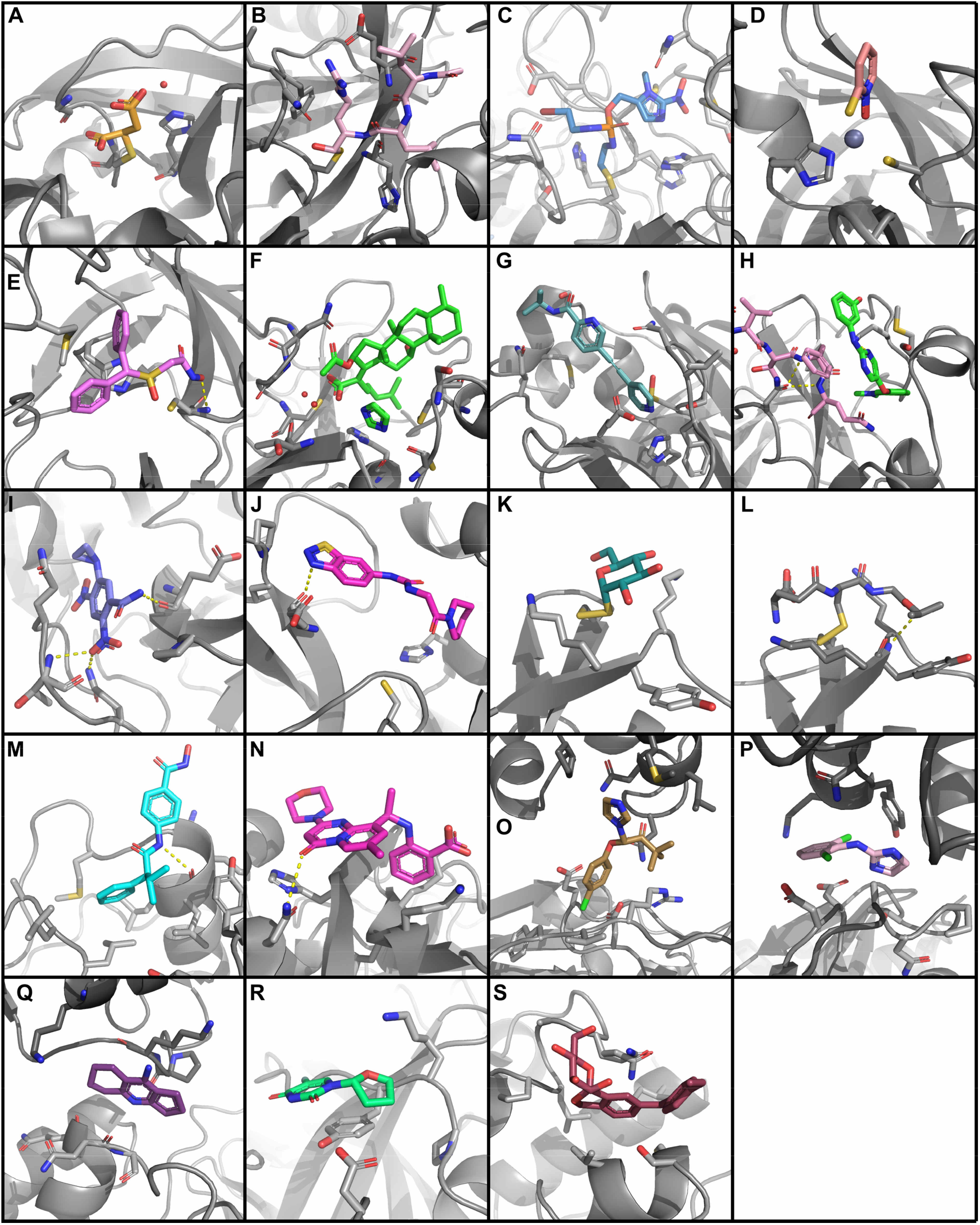
The structures of inactive compounds. Compounds are depicted as colored sticks. M^pro^ is shown as a grey cartoon model with residues important for ligand binding shown as stick models and hydrogen bonds are indicated by dashed lines. Ligands binding covalently to the active site residue Cys145: **A**, maleate. **B**, leupeptin. **C**, TH-302. **D**, zinc pyrithione. Ligands binding non-covalently to the active site: **E**, adrafinil. **F**, fusidic acid. **G**, LSN-2463359. **H**, SEN1269 (C-terminus of neighboring M^pro^ protomer shown as pink stick model). **I**, tretazicar. **J**, UNC2327. Covalent binders to Cys156: **K**, aurothioglucose. **L**, glutathione isopropylester. Other surface pockets: **M**, AR-42. **N**, AZD6482. **O**, climbazole. **P**, clonidine. **Q**, ipidacrine. **R**, tegafur. Allosteric binding site: **S**, tofogliflozin.

**Fig. S4.**
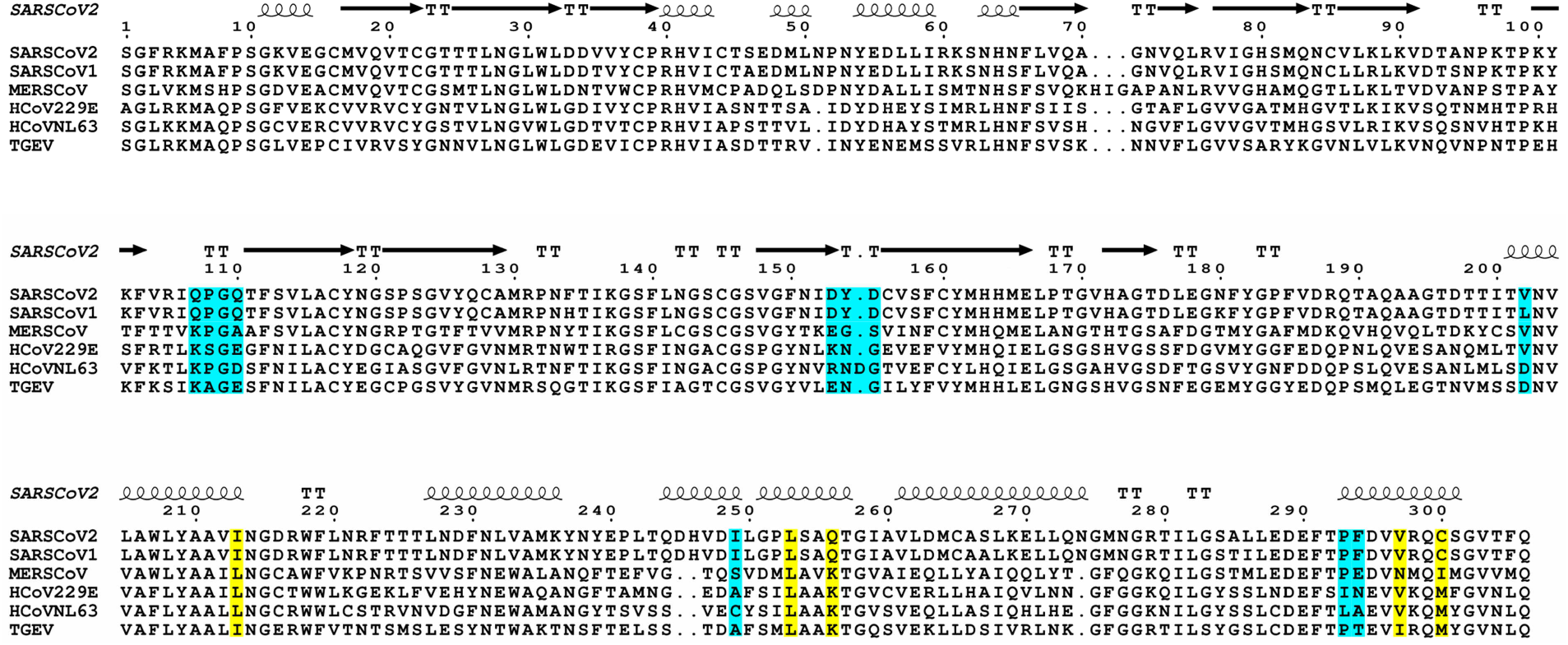
Sequence comparison of M^pro^ of α-(HCoV229E, HCovNL63, TGEV) and β-(SARS-CoV-1&2, MERS-CoV) coronaviruses. SARS-CoV-2 M^pro^-residues interacting with compounds in allosteric site 1 (blue) and 2 (yellow) are indicated in colored boxes.

**Fig. S5.**
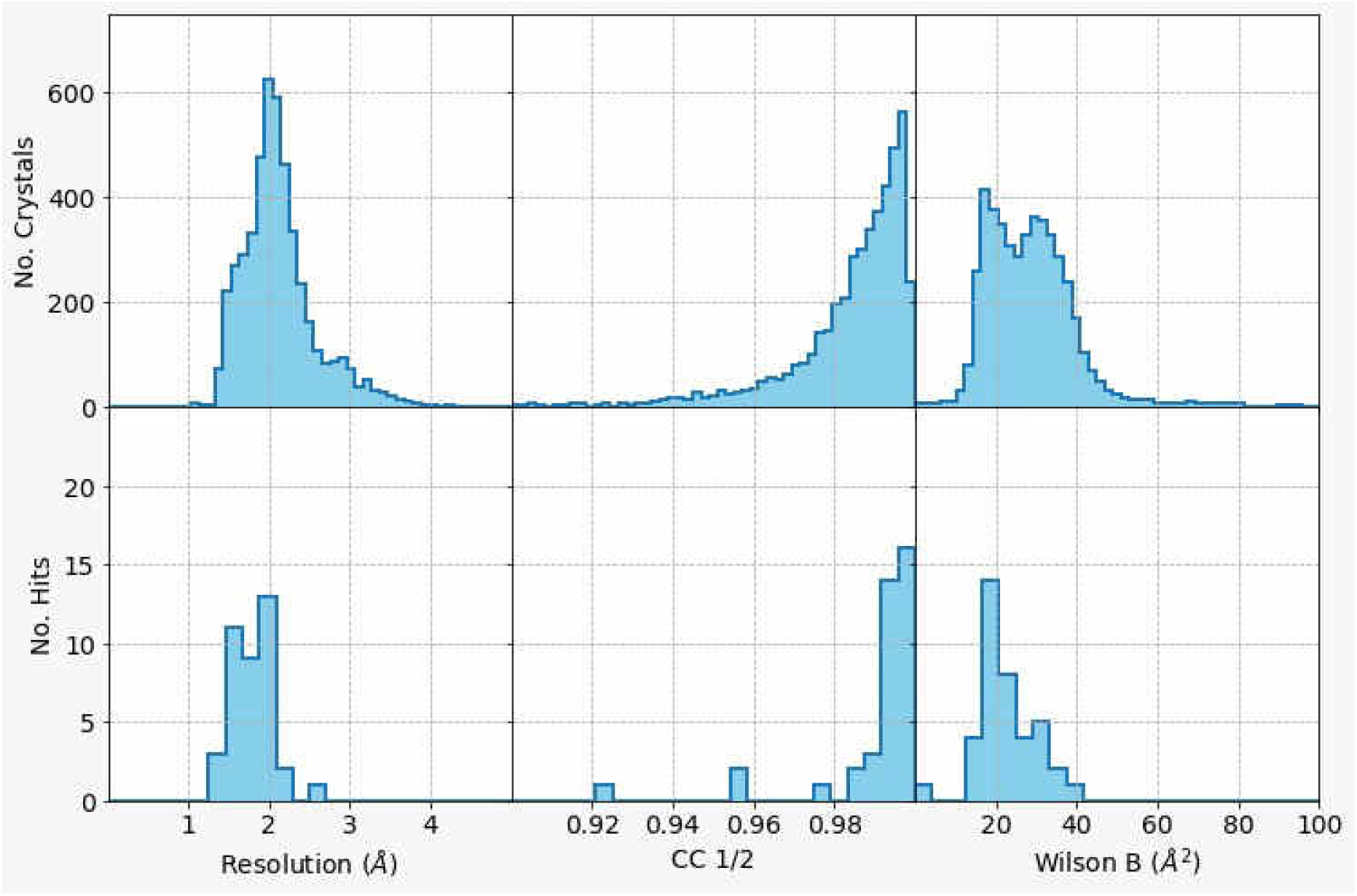
Distribution of data quality indicators of all collected X-ray diffraction datasets (upper panel) and of datasets with identified compound (lower panel): diffraction resolution (left), CC1/2 of the datasets (middle), and Wilson B-factor (right).

**Table S1.**

Comprehensive summary sheets of hit compounds showing electron-density maps, compound interactions with M^pro^, detailed compound information, biochemical and cell-based antiviral reduction data.

Table provided as separate file.

**Table S2.**

Summary of X-ray crystallographic data processing and refinement statistics. Table provided as separate file. *In vitro* antiviral activity, cytotoxicity and selectivity of selected compounds against SARS-CoV-2. EC_50_-half-maximal effective concentration; CC_50_-half-maximal cytotoxic concentration; n.a.-not available. Viral titers, vRNA yield and cell viability were determined by RT-qPCR, immunofocus assays, and the CCK-8 method, respectively. Values were calculated from three independent replicates in one experiment.

**Table S3.**
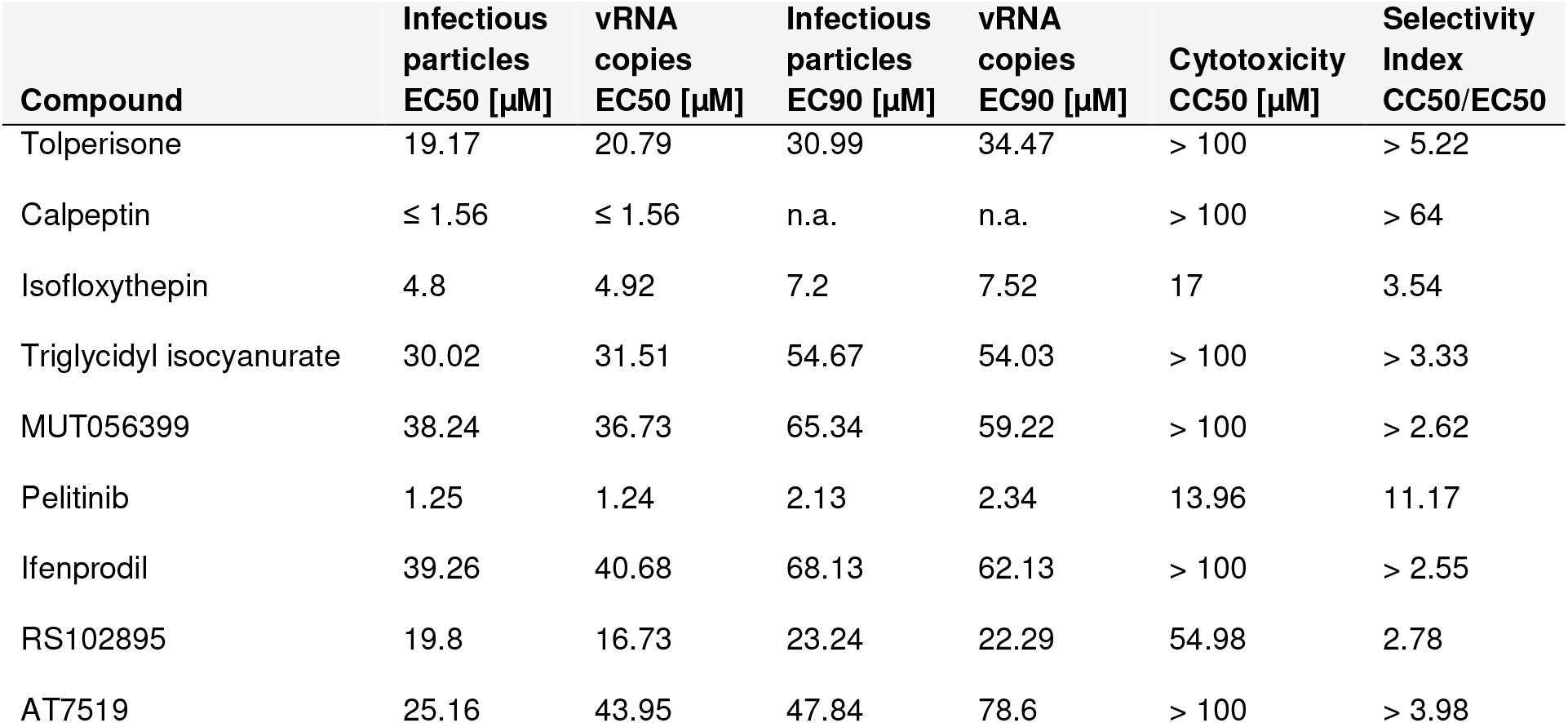
*In vitro* antiviral activity, cytotoxicity and selectivity of selected compounds against SARSCoV-2. EC_50_- half-maximal effective concentration; CC_50_- half-maximal cytotoxic concentration; n.a.- not available. Viral titers, vRNA yield and cell viability were determined by RT-qPCR, immunofocus assays, and the CCK-8 method, respectively. Values were calculated from three independent replicates in one experiment.

| Compound | Infectious particles<br>EC50 [μM] | vRNA copies<br>EC50 [μM] | Infectious particles<br>EC90 [μM] | vRNA copies<br>EC90 [μM] | Cytotoxicity<br>CC50 [μM] | Selectivity<br>Index<br>CC50/EC50 |
| --- | --- | --- | --- | --- | --- | --- |
| Tolperisone | 19.17 | 20.79 | 30.99 | 34.47 | > 100 | > 5.22 |
| Calpeptin | ≤ 1.56 | ≤ 1.56 | n.a. | n.a. | > 100 | > 64 |
| Isofloxythepin | 4.8 | 4.92 | 7.2 | 7.52 | 17 | 3.54 |
| Triglycidyl isocyanurate | 30.02 | 31.51 | 54.67 | 54.03 | > 100 | > 3.33 |
| MUT056399 | 38.24 | 36.73 | 65.34 | 59.22 | > 100 | > 2.62 |
| Pelitinib | 1.25 | 1.24 | 2.13 | 2.34 | 13.96 | 11.17 |
| Ifenprodil | 39.26 | 40.68 | 68.13 | 62.13 | > 100 | > 2.55 |
| RS102895 | 19.8 | 16.73 | 23.24 | 22.29 | 54.98 | 2.78 |
| AT7519 | 25.16 | 43.95 | 47.84 | 78.6 | > 100 | > 3.98 |

**Table S4.** Native MS verified binding of compounds to M^pro^. The table shows compounds and their molecular weight. Mass spectra of compounds and M^pro^ (final conc. 50 µ M and 10 µM) were analyzed by converting peak intensities into intensity fractions for zero, one and two ligands (0/1/2 ligands in %) bound per M^pro^ dimer. Mass of the fragmented compounds is given when the observed mass is deviating from the expected mass. Type and mass of counterion is indicated if applicable.

| Compound (counter ion) | Compound/counter ion mass [Da] | Intensity fraction (0/1/2 compounds per M<sup>pro</sup> dimer) [%] | Mass of fragmented compounds [Da] |
| --- | --- | --- | --- |
| Calpeptin | 362.5 | 5 / 27 / 68 |  |
| Triglycidyl isocyanurate | 297.3 | 8 / 38 / 53 |  |
| Zinc Pyrithione | 317.7 | 11 / 34 / 55 | 128 |
| HEAT (HCl) | 295.4 / 35 | 39 / 37 / 24 | 181 |
| HTMT (dimaleate) | 382.4 / 2*116.1 | 55 / 24 / 21 | 131 |
| Dexrazoxan | 268.3 | 56 / 29 / 15 |  |
| Adrafinil | 289.35 | 58 / 28 / 14 |  |
| TH-302 | 449.04 | 59 / 30 / 10 | 365 |
| Ifenprodil (hemitartrate) | 325.2 / 74.0 | 61 / 24 / 15 | 126 |
| AZD6482 | 408.5 | 62 / 29 / 9 |  |
| Glutathione-isopropyl-ester | 349.41 | 62 / 28 / 10 | 126/188 |
| AT7519 | 382.24 | 65 / 28 / 7 |  |
| AL-8697 | 402.41 | 67 / 24 / 8 |  |
| Cinepazide (maleate) | 417.5 / 116.1 | 72 / 21 / 7 |  |
| UNC2327 | 319.4 | 76 / 24 / 0 |  |
| Fusidic acid | 516.7 | 78 / 18 / 5 |  |
| AR-42 | 312.4 | 78 / 18 / 4 | 580 |
| PD-168568 (HCl)<sub>2</sub> | 440.41 / 2*35 | 81 / 16 / 3 | 350 |
| Tofogliflozin (hydrate) | 404.45 | 82 / 18 / 0 | 380 |
| MUT056399 | 293.27 | 83 / 17 / 0 |  |
| Colistimethate (Na) | 1735.82 / 23 | 83 / 15 / 2 |  |
| Vicriviroc (maleate) | 533.6 / 116.1 | 88 / 12 / 0 | 535 |
| Pelitinib | 467.92 | 88 / 12 / 0 |  |

**Table S5.**

The highest ranked 200 compounds of the virtual screening. The names and HYDE scores of the top ranked molecules are given. The yellow background highlights compounds for which high-quality X-ray data was obtained in the X-ray screening. The green background highlights compounds that were detected in the active site in the X-ray screen. Compounds highlighted in light green show a similar binding mode to the fragment with the PDB ligand ID K0G in complex with M^pro^ (PDB ID 5R83). Compounds highlighted in light yellow were reported as being active in other screening studies. Table provided as separate file.

