## Supplemental Table S1 for "Inhibition of SARS-CoV-2 main protease by allosteric drug-binding"

#### Hit#1: Leupeptin bound to SARS-CoV-2 MPro, PDB: 6YZ6

**Synonyms:** Leupeptin, Ac-Leu-Leu-Arg-H, Ac-Leu-Leu-Arginal

**PubChem CID:** 72429

**CAS:** 55123-66-5

**Molecular weigth:** 426.6 g/mol

**Binding type:** covalent, hemithioacetal

**Binding location:** Cystein 145

**Screen compound ID:** - (positive control)

**Original compound:**

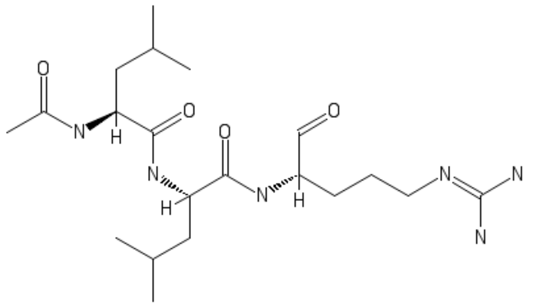

Lig-Plot interaction network

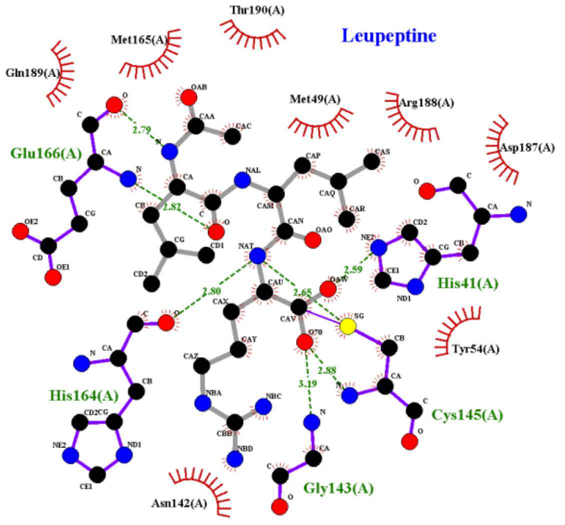


**Isomeric smiles:**
CC(C)C[C@@H](C(=O)N[C@@H](CC(C)C)C(=O)N[C@@H](CCCN=C(N)N)C=O)NC(=O)C

**Refinement 2Fo-Fc map**, 1.2 σ-level

note

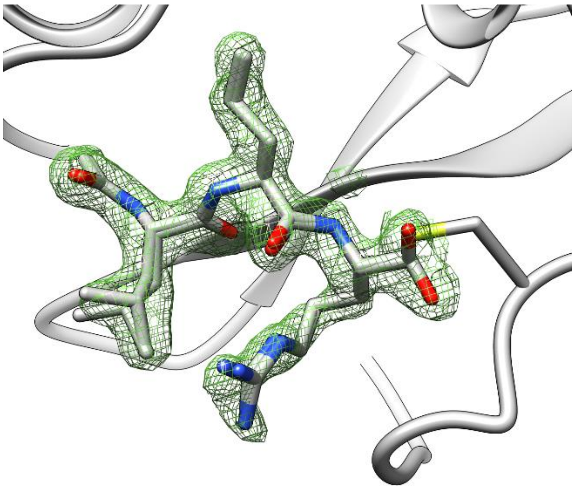

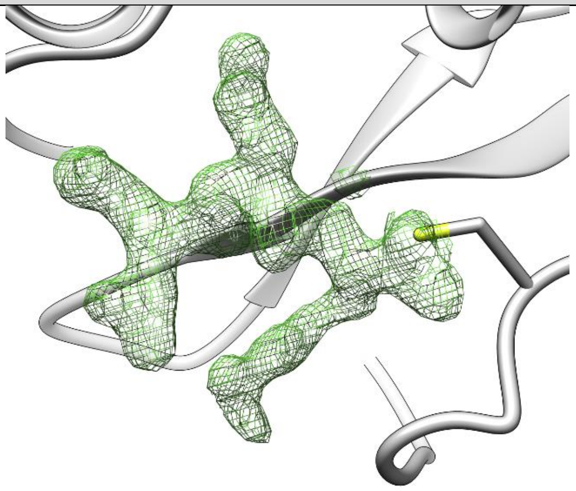

| | |
|---|---|
| **PDB structure code:** 6YZ6 | **PDB Ligand code:** AR7 |
| **Resolution:** 1.7 Å | **$R_{work}$/$R_{free}$:** 0.17/0.22 |
| **Occupancy:** 0.73, 0.21 | **Ligand RSCC/RSR:** 0.86/0.19, 0.86/0.19 |
| **Biochemistry & Cell biology** | |
| **Anti viral activity $EC_{50}$:** N.a. | **vRNA yield $EC_{50}$:** N.a. |
| **Anti viral activity $EC_{90}$:** N.a. | **vRNA yield $EC_{90}$:** N.a. |
| **Cytotoxicity $CC_{50}$:** N.a. | **SI CC50/EC50:** N.a. |
| **Native MS adduct saturation:** N.a. | **Native MS adduct size:** N.a. |

#### Hit#2: Calpeptin bound to SARS-CoV-2 MPro, PDB: 7AKU

| | |
|---|---|
| **Synonyms:** Calpeptin, N-Cbz-leu-nleu-al, Benzylcarbonyl-leu-nleu-H | Lig-Plot interaction network |
| **PubChem CID:** 73364 | |
| **CAS:** 117591-20-5 | |
| **Molecular weigth:** 362.5 g/mol | |
| **Binding type:** covalent, hemithioacetal | |
| **Binding location:** Cystein 145 | |
| **Screen compound ID:** SPE_K26134695 | |
| **Original compound:** | |

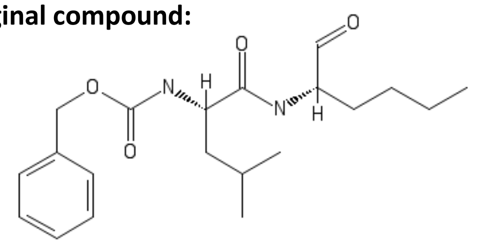

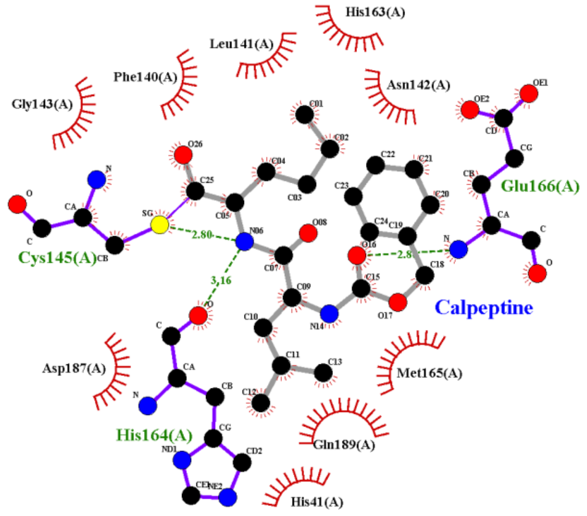


**Isomeric smiles:** CCCC[C@@H](C=O)NC(=O)[C@H](CC(C)C)NC(=O)OCC1=CC=CC=C1

**Refinement 2Fo-Fc map**, 1.4 σ-level

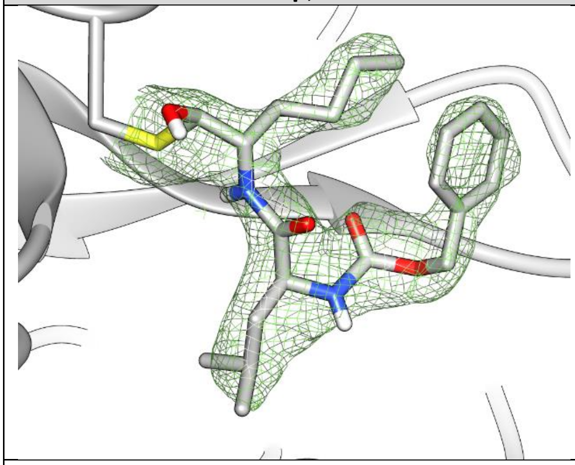

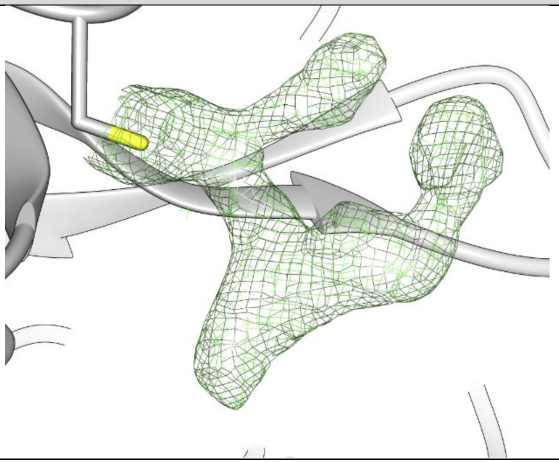

| | |
|---|---|
| **PDB structure code:** 7AKU | **PDB Ligand code:** RN2 |
| **Resolution:** 2.5 Å | $\mathbf{R_{work}/R_{free}}$: 0.19/0.24 |
| **Occupancy:** 1.0 | **Ligand RSCC/RSR:** 0.94/0.16 |
| **Biochemistry & Cell biology** | |
| **Anti viral activity $EC_{50}$:** N.a. | **vRNA yield $EC_{50}$:** N.a. |
| **Anti viral activity $EC_{90}$:** N.a. | **vRNA yield $EC_{90}$:** N.a. |
| **Cytotoxicity $CC_{50}$:** N.a. | **SI CC50/EC50:** N.a. |
| **Native MS adduct saturation:** 82% (5 / 27 68) | **Native MS adduct size:** 362.5 Da |

#### Hit#3: HEAT bound to SARS-CoV-2 MPro, PDB: 6YNQ

**Synonyms:** HEAT, BE-2254, 2-(beta-(4-Hydroxyphenyl)ethylaminomethyl)tetralone

**PubChem CID:** 34772

**CAS:** 40077-13-2

**Molecular weigth:** 295.4 g/mol

**Binding type:** covalent, thioether

**Binding location:** Cystein 145

**Screen compound ID:** SPE_A24429032

**Original compound:**

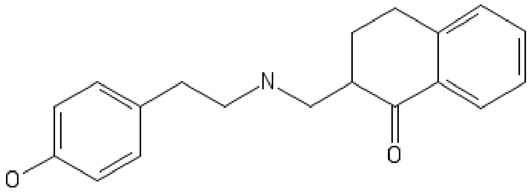


Lig-Plot interaction network

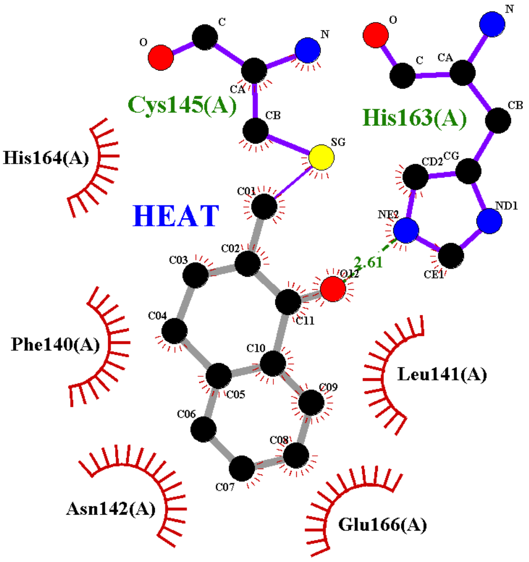


**Isomeric smiles:** C1CC2=CC=CC=C2C(=O)C1CNCCC3=CC=C(C=C3)O

**Refinement 2Fo-Fc map**, 1.4 σ-level

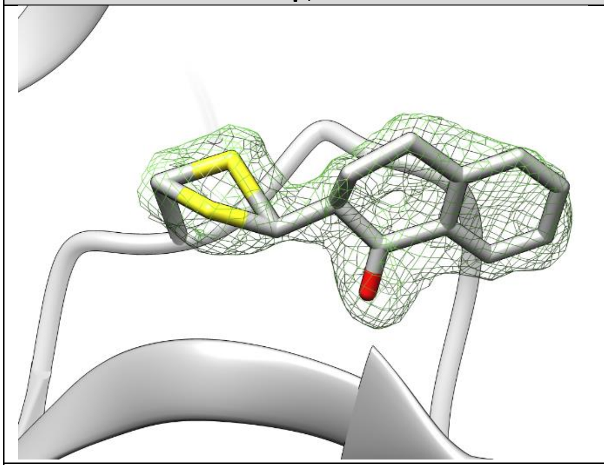

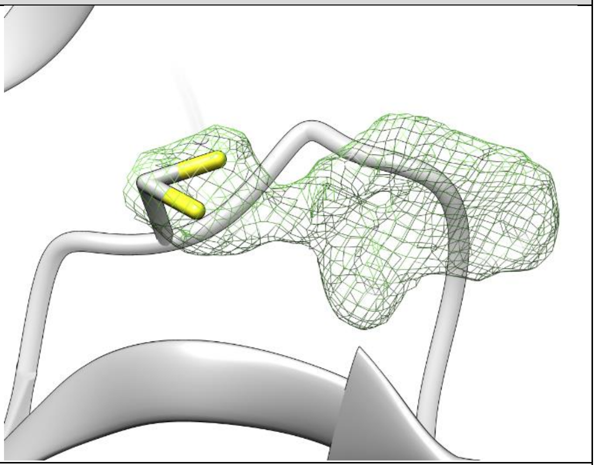

| | |
|---|---|
| **PDB structure code:** 6YNQ | **PDB Ligand code:** P6N |
| **Resolution:** 1.8 Å | **$R_{work}$/$R_{free}$:** 0.19/0.23 |
| **Occupancy:** 1.0 | **Ligand RSCC/RSR:** 0.92/0.11 |
| **Biochemistry & Cell biology** | |
| **Anti viral activity $EC_{50}$:** N.a. | **vRNA yield $EC_{50}$:** N.a. |
| **Anti viral activity $EC_{90}$:** N.a. | **vRNA yield $EC_{90}$:** N.a. |
| **Cytotoxicity $CC_{50}$:** N.a. | **SI CC50/EC50:** N.a. |
| **Native MS adduct saturation:** 43% (39/37/24) | **Native MS adduct size:** 181 Da |

### Hit#4: Tolperisone HCl bound to SARS-CoV-2 MPro, PDB: 7ADW

**Synonyms:** Tolperisone, Mydeton, Mydocalm, Mideton

**PubChem CID:** 5511

**CAS:** 728-88-1

**Molecular weigth:** 245.36 g/mol

**Binding type:** covalent, thioether

**Binding location:** Cystein 145

**Screen compound ID:** SPE_A27732521

**Original compound:**

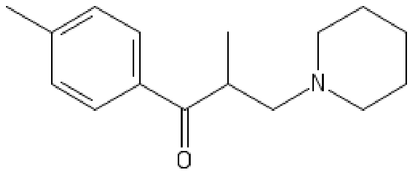


Lig-Plot interaction network

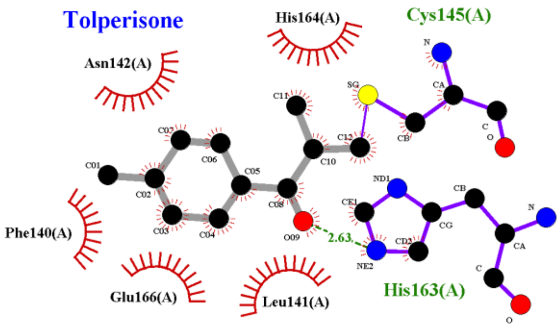


**Isomeric smiles:** CC1=CC=C(C=C1)C(=O)C(C)CN2CCCCC2

**Refinement 2Fo-Fc map**, 1.0 σ-level

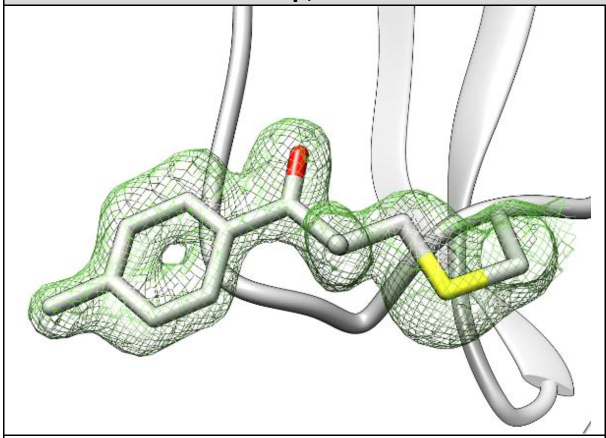

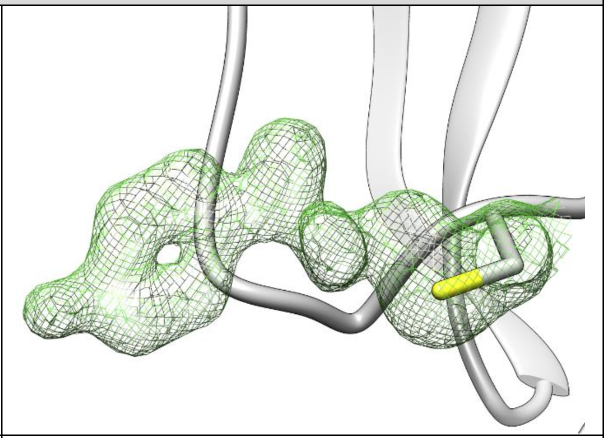

| | |
|---|---|
| **PDB structure code:** 7ADW | **PDB Ligand code:** R7Q |
| **Resolution:** 1.62 Å | **$R_{work}$/$R_{free}$:** 0.20/0.23 |
| **Occupancy:** 1.0 | **Ligand RSCC/RSR:** 0.87/0.11 |

**Biochemistry & Cell biology**

| | |
|---|---|
| **Anti viral activity $EC_{50}$:** 17.16 ± 1.76 µM | **vRNA yield $EC_{50}$:** 22.42 ± 2.25 µM |
| **Anti viral activity $EC_{90}$:** 29.68 ± 2.32 µM | **vRNA yield $EC_{90}$:** 34.65 ± 3.76 µM |
| **Cytotoxicity $CC_{50}$:** > 100 µM | **SI CC50/EC50:** > 5.83 |
| **Native MS adduct saturation:** - | **Native MS adduct size:** N.a. |

#### Hit#5: Maleate (Quipazine) bound to SARS-CoV-2 MPro, PDB: 7AHA

**Synonyms:** Maleic acid, cis-butenedioic acid, 110-16-7, Toxilic acid, Maleinic acid

**PubChem CID:** 444266 (Maleate) / 5011 (Quip.)

**CAS:** 110-16-7 (Maleate) / 5786-68-5

**Molecular weigth:** 116.07 g/mol (Maleate)

**Binding type:** covalent, thioether

**Binding location:** Cystein 145

**Screen compound ID:** SPE_K77925998

**Original compound:**

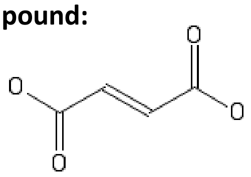

Lig-Plot interaction network

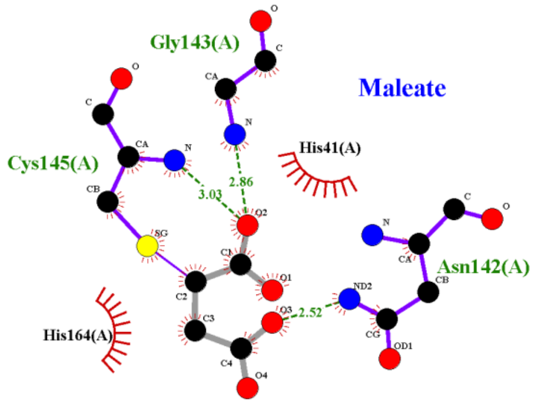


**Isomeric smiles:** C(=C\C(=O)[O-])\C(=O)[O-] (Maleate)

**Refinement 2Fo-Fc map**, 0.8 σ-level

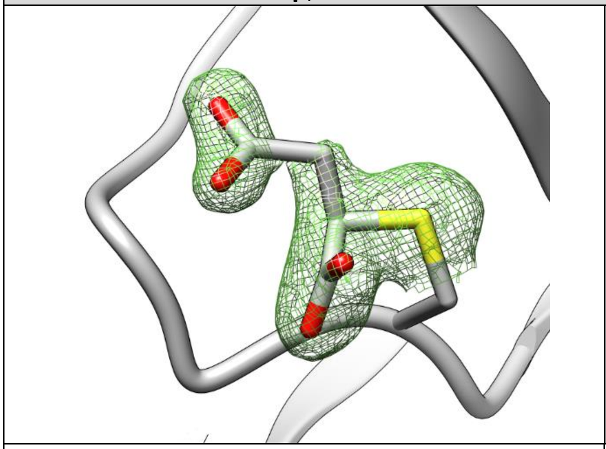

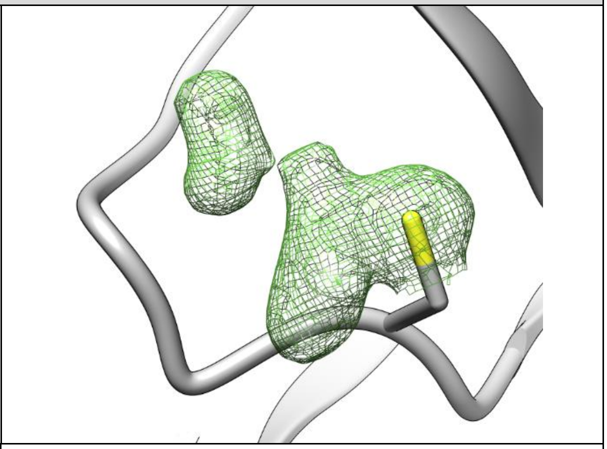

| | |
|---|---|
| **PDB structure code:** 7AHA | **PDB Ligand code:** SIN |
| **Resolution:** 1.68 Å | **$R_{work}$/$R_{free}$:** 0.17/0.20 |
| **Occupancy:** 1.0 | **Ligand RSCC/RSR:** 0.89/0.16 |
| **Biochemistry & Cell biology** | |
| **Anti viral activity $EC_{50}$:** N.a. | **vRNA yield $EC_{50}$:** N.a. |
| **Anti viral activity $EC_{90}$:** N.a. | **vRNA yield $EC_{90}$:** N.a. |
| **Cytotoxicity $CC_{50}$:** N.a. | **SI CC50/EC50:** N.a. |
| **Native MS adduct saturation:** - | **Native MS adduct size:** N.a. |

, Deutsches Elektronen Synchrotron, Notkestraße 85, 22607 Hamburg

#### Hit#6: Isofloxythepin bound to SARS-CoV-2 MPro, PDB: 7AY7

**Synonyms:** Isofloxythepin, EINECS 275-028-1, 70931-18-9

**PubChem CID:** 115193

**CAS:** 70931-18-9

**Molecular weigth:** 400.6 g/mol

**Binding type:** covalent, thioether

**Binding location:** Cystein 145

**Screen compound ID:** DOM_SIM_130

**Original compound:**

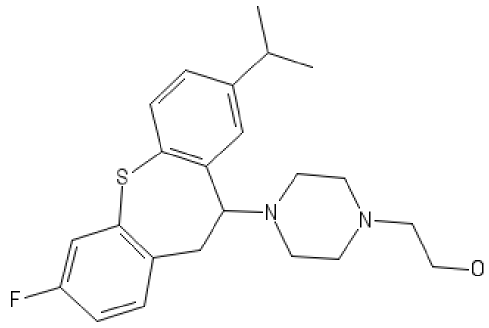


Lig-Plot interaction network

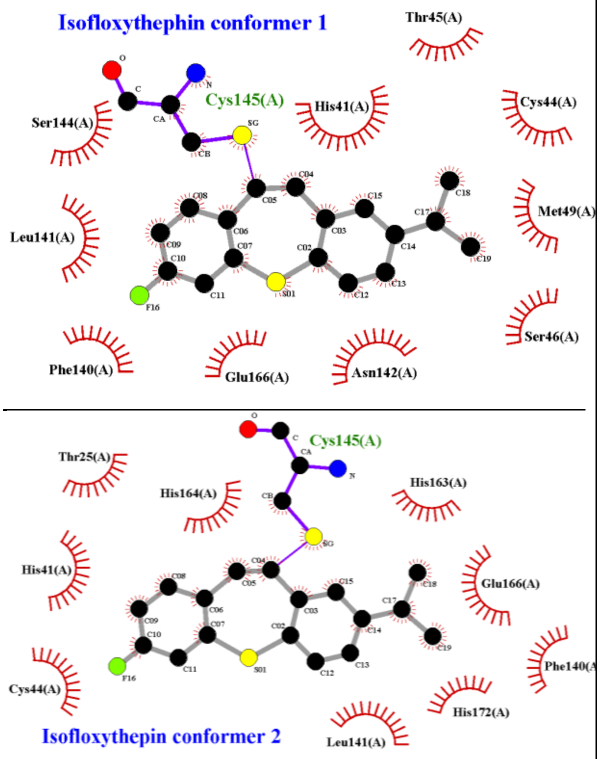


**Isomeric smiles:** CC(C)C1=CC2=C(C=C1)SC4=C(CC2N3CCN(CC3)CCO)C=CC(=C4)F

**Refinement 2Fo-Fc map**, 0.7 σ-level, **PanDDA event map BDC 0.15** , 1.4 σ-level

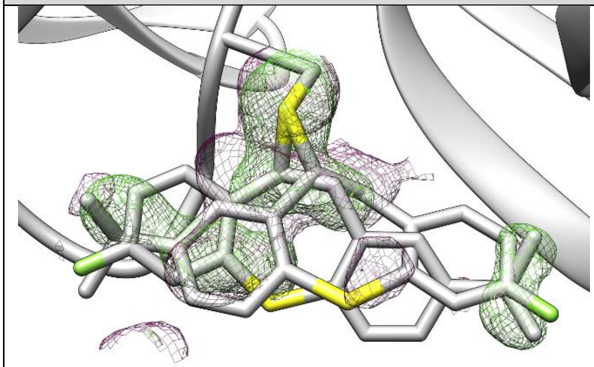

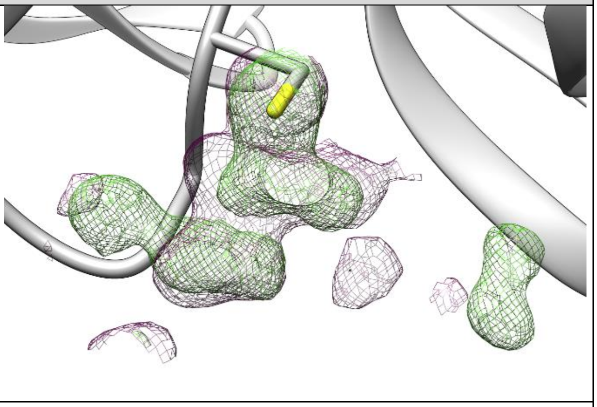

2Fo-Fc map, PanDDA event map

| | |
|---|---|
| **PDB structure code:** 7AY7 | **PDB Ligand code:** S8T |
| **Resolution:** 1.55 Å | **$R_{work}$/$R_{free}$:** 0.17/0.20 |
| **Occupancy:** 0.30, 0.38 | **Ligand RSCC/RSR:** 0.60/0.51, 0.72/43 |

**Biochemistry & Cell biology**

| | |
|---|---|
| **Anti viral activity $EC_{50}$:** 4.82 ± 0.33 µM | **vRNA yield $EC_{50}$:** 4.94 ± 0.58 µM |
| **Anti viral activity $EC_{90}$:** 7.21 ± 0.31 µM | **vRNA yield $EC_{90}$:** 7.52 ± 0.56 µM |
| **Cytotoxicity $CC_{50}$:** 14.49 ± 3.57 µM | **SI CC50/EC50:** > 4.04 |
| **Native MS adduct saturation:** - | **Native MS adduct size:** N.a. |

#### Hit#7: Triglycidyl isocyanurate bound to SARS-CoV-2 MPro, PDB: 7AQJ

**Synonyms:** Triglycidyl isocyanurate, 2451-62-9, 1,3,5-Triglycidyl isocyanurate, Teroxirone

**PubChem CID:** 17142

**CAS:** 6990-06-3

**Molecular weigth:** 297.26 g/mol

**Binding type:** covalent, thioether

**Binding location:** Cystein 145

**Screen compound ID:** SPE_A06935312

**Original compound:**

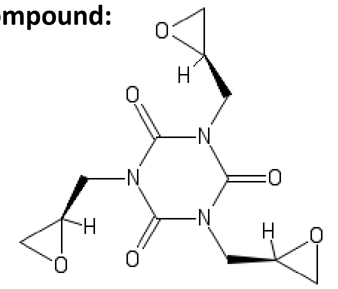

Lig-Plot interaction network

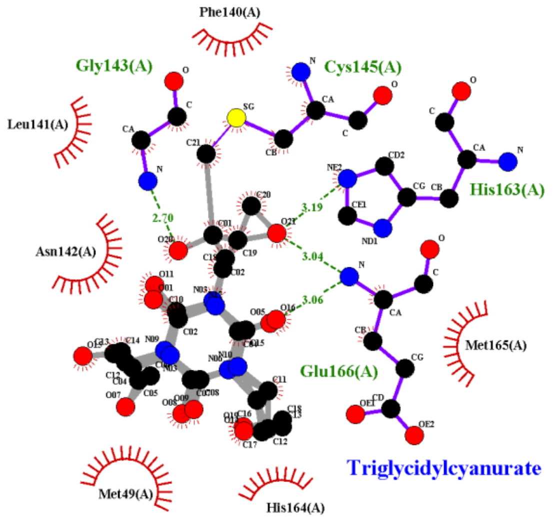


**Isomeric smiles:** C1[C@H](O1)CN2C(=O)N(C(=O)N(C2=O)C[C@@H]3CO3)C[C@@H]4CO4

**Refinement 2Fo-Fc map**, 0.7 σ-level

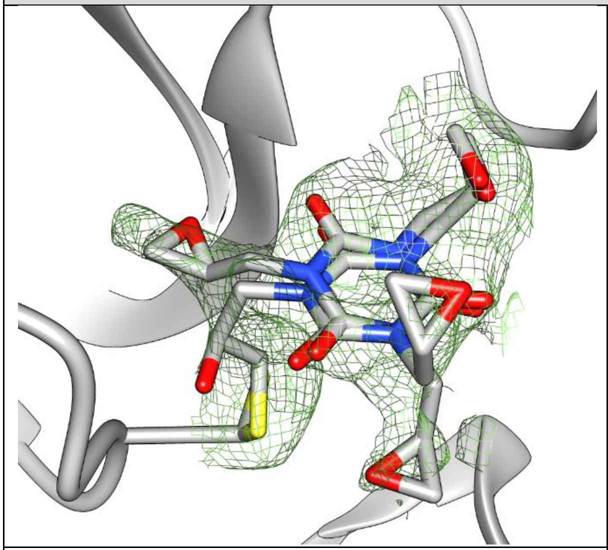

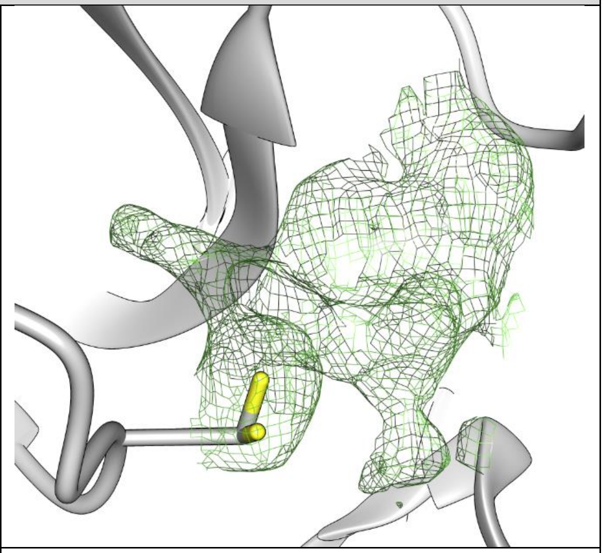

| | |
|---|---|
| **PDB structure code:** 7AQJ | **PDB Ligand code:** RV8, S7H |
| **Resolution:** 2.59 Å | **$R_{work}$/$R_{free}$:** 0.22/0.26 |
| **Occupancy:** 0.4 / 0.47 | **Ligand RSCC/RSR:** 0.78/0.27, 0.84/0.22 |

**Biochemistry & Cell biology**

| | |
|---|---|
| **Anti viral activity $EC_{50}$:** 35.68 ± 10.87 µM | **vRNA yield $EC_{50}$:** 37 ± 5.35 µM |
| **Anti viral activity $EC_{90}$:** 48.08 ± 9.57 µM | **vRNA yield $EC_{90}$:** 50.25 ± 4.14 µM |
| **Cytotoxicity $CC_{50}$:** > 100 µM | **SI CC50/EC50:** > 2.8 |
| **Native MS adduct saturation:** 72% (8/38/53) | **Native MS adduct size:** 297 Da |

### Hit#8: MUT056399 acid bound to SARS-CoV-2 MPro, PDB: 7AP6

| | |
|---|---|
| **Synonyms:** MUT056399, 1269055-85-7, FAB-001, 4-(4-ethyl-5-fluoro-2-hydroxyphenoxy)-3-fluorobenzamide<br>**PubChem CID:** 44208849<br>**CAS:** 1269055-85-7<br>**Molecular weigth:** 293.26 g/mol<br>**Binding type:** non-covalent<br>**Binding location:** active site pocket<br>**Screen compound ID:** SPE_K72078047<br>**Original compound:** 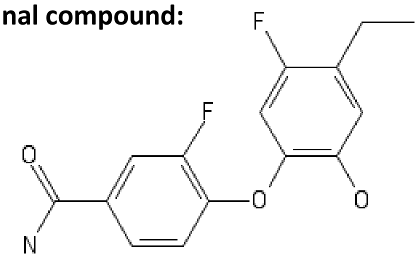 | Lig-Plot interaction network 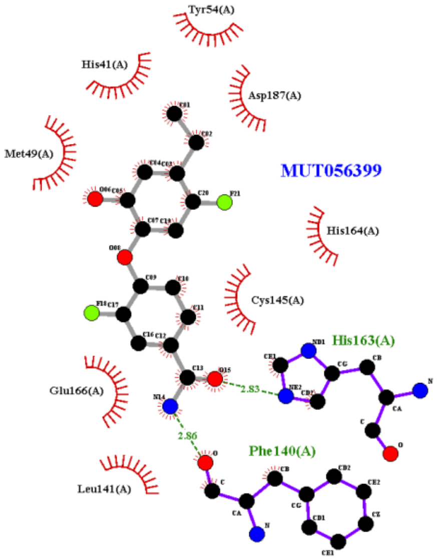  |
| **Isomeric smiles:** CCC1=CC(=C(C=C1F)OC2=C(C=C(C=C2)C(=O)N)F)O | |
| **Refinement 2Fo-Fc map**, 0.7 σ-level, **PanDDA event map BDC 0.18** , 1.4 σ-level | |
| 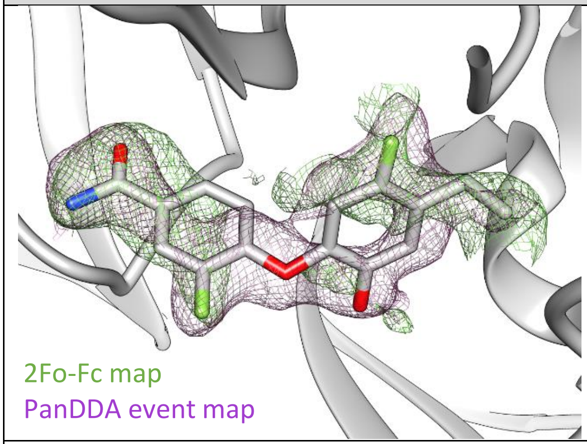  | 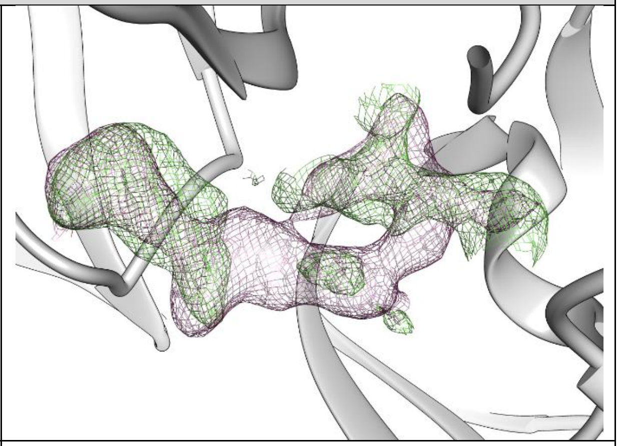 |
| **PDB structure code:** 7AP6 | **PDB Ligand code:** RQN |
| **Resolution:** 1.78 Å | **$R_{work}$/$R_{free}$:** 0.20/0.23 |
| **Occupancy:** 0.44 | **Ligand RSCC/RSR:** 0.64/0.54 |
| **Biochemistry & Cell biology** | |
| **Anti viral activity $EC_{50}$:** 43.27 ± 0.59 µM | **vRNA yield $EC_{50}$:** 42.18 ± 3.86 µM |
| **Anti viral activity $EC_{90}$:** 54.05 ± 0.48 µM | **vRNA yield $EC_{90}$:** 54.27 ± 2.27 µM |
| **Cytotoxicity $CC_{50}$:** > 100 µM | **SI CC50/EC50:** > 2.31 |
| **Native MS adduct saturation:** 6% (83/17/0) | **Native MS adduct size:** 293 Da |

, Deutsches Elektronen Synchrotron, Notkestraße 85, 22607 Hamburg

#### Hit#9: Pelitinib bound to SARS-CoV-2 MPro, PDB: 7AXM

**Synonyms:** Pelitinib, 257933-82-7, EKB-569

**PubChem CID:** 6445562

**CAS:** 257933-82-7

**Molecular weigth:** 467.9 g/mol

**Binding type:** non-covalent

**Binding location:** allosteric site I

**Screen compound ID:** DOM_SIM_123

**Original compound:**

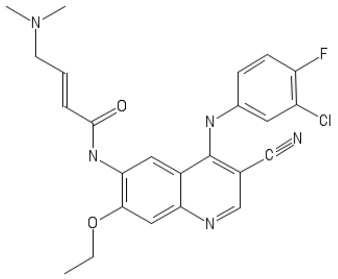

Lig-Plot interaction network

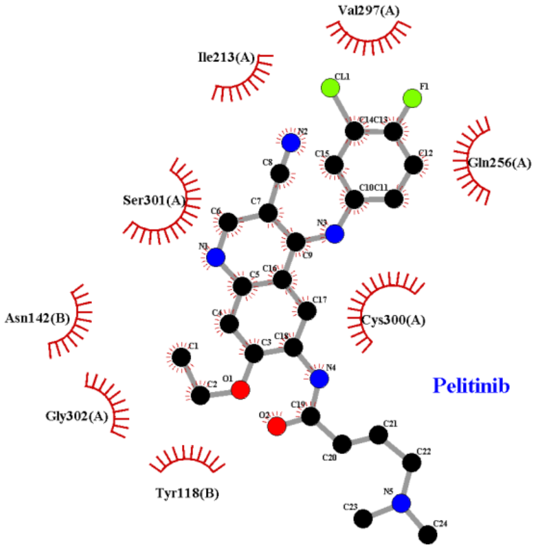


**Isomeric smiles:** CCOC1=C(C=C2C(=C1)N=CC(=C2NC3=CC(=C(C=C3)F)Cl)C#N)NC(=O)/C=C/CN(C)C

**Refinement 2Fo-Fc map**, 0.6 σ-level, **PanDDA event map BDC 0.18** , 1.0 σ-level

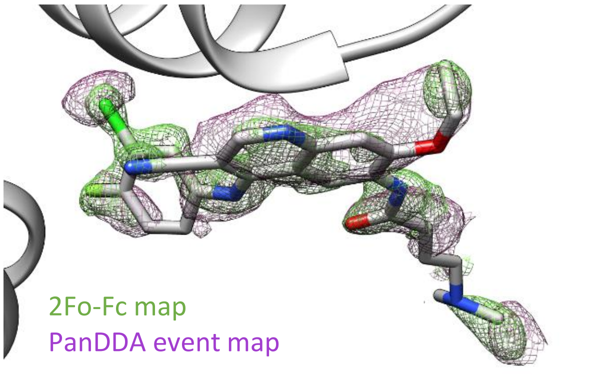


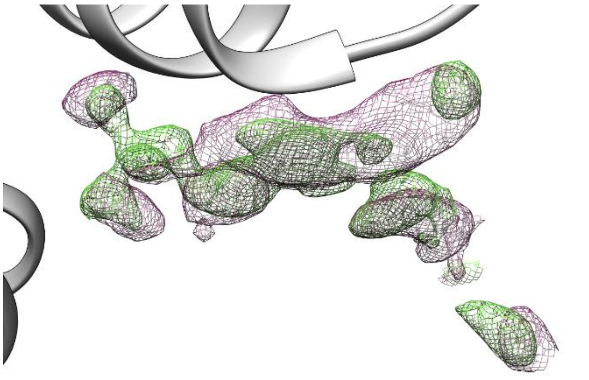

| | |
|---|---|
| **PDB structure code:** 7AXM | **PDB Ligand code:** 93J |
| **Resolution:** 1.4 Å | **$R_{work}$/$R_{free}$:** 0.14/0.20 |
| **Occupancy:** 0.61 | **Ligand RSCC/RSR:** 0.61/0.28 |

**Biochemistry & Cell biology**

| | |
|---|---|
| **Anti viral activity $EC_{50}$:** 1.25 ± 0.02 µM | **vRNA yield $EC_{50}$:** 1.21 ± 0.1 µM |
| **Anti viral activity $EC_{90}$:** 2.13 ± 0.12 µM | **vRNA yield $EC_{90}$:** 2.32 ± 0.1 µM |
| **Cytotoxicity $CC_{50}$:** > 100 µM | **SI CC50/EC50:** 11.17 |
| **Native MS adduct saturation:** 12% (88/12/0) | **Native MS adduct size:** 467 Da |

, Deutsches Elektronen Synchrotron, Notkestraße 85, 22607 Hamburg

#### Hit#10: Ifenprodil bound to SARS-CoV-2 MPro, PDB: 7AQI

**Synonyms:** Ifenprodil, 23210-56-2, ifenprodil tartrate, Vadilex, Dilvax

**PubChem CID:** 3689

**CAS:** 23210-56-2

**Molecular weigth:** 325.4 g/mol

**Binding type:** non-covalent

**Binding location:** allosteric site I

**Screen compound ID:** SPE_A06935312

**Original compound:**

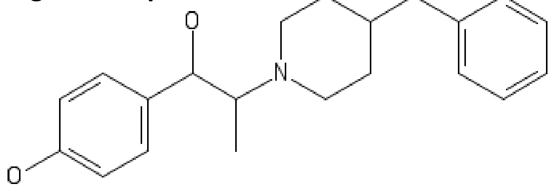


Lig-Plot interaction network

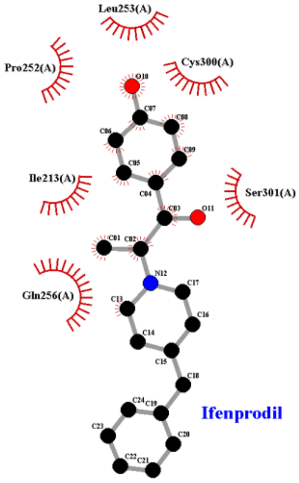


**Isomeric smiles:** CC(C(C1=CC=C(C=C1)O)O)N2CCC(CC2)CC3=CC=CC=C3

**Refinement 2Fo-Fc map**, 0.7 σ-level

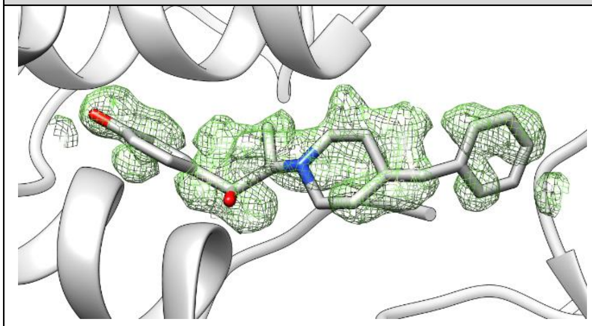

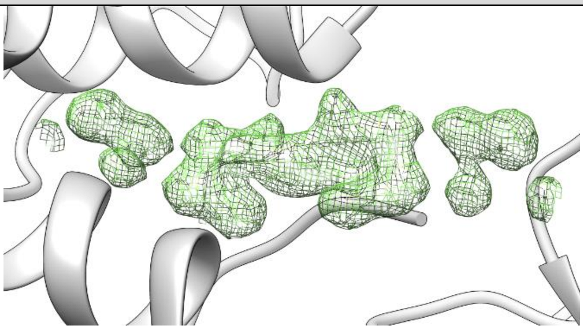

| | |
|---|---|
| **PDB structure code:** 7AQI | **PDB Ligand code:** QEL |
| **Resolution:** 1.7 Å | **$R_{work}$/$R_{free}$:** 0.23/0.26 |
| **Occupancy:** 0.5 + 0.5 | **Ligand RSCC/RSR:** 0.80/0.19 |
| **Biochemistry & Cell biology** | |
| **Anti viral activity $EC_{50}$:** 43.37 ± 2.99 µM | **vRNA yield $EC_{50}$:** 44.96 ± 5.52 µM |
| **Anti viral activity $EC_{90}$:** 51.25 ± 2.34 µM | **vRNA yield $EC_{90}$:** 53.73 ± 1.74 µM |
| **Cytotoxicity $CC_{50}$:** > 100 µM | **SI CC50/EC50:** > 2.31 |
| **Native MS adduct saturation:** 27% (61/24/15) | **Native MS adduct size:** 126 Da |

, Deutsches Elektronen Synchrotron, Notkestraße 85, 22607 Hamburg

### Hit#11: RS102895 bound to SARS-CoV-2 MPro, PDB: 7ABU

**Synonyms:** RS102895, GTPL779, CHEMBL1593104, SCHEMBL9972649

**PubChem CID:** 10000456

**CAS:** 1173022-16-6

**Molecular weigth:** 390.4 g/mol

**Binding type:** non-covalent

**Binding location:** allosteric site I

**Screen compound ID:** SPE_K83063356

**Original compound:**

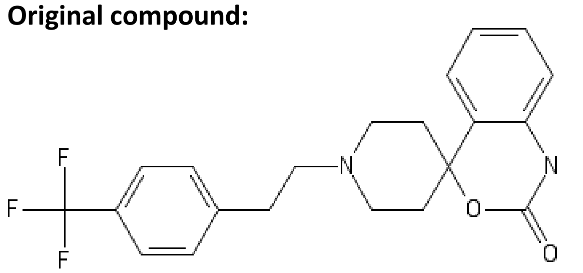


Lig-Plot interaction network

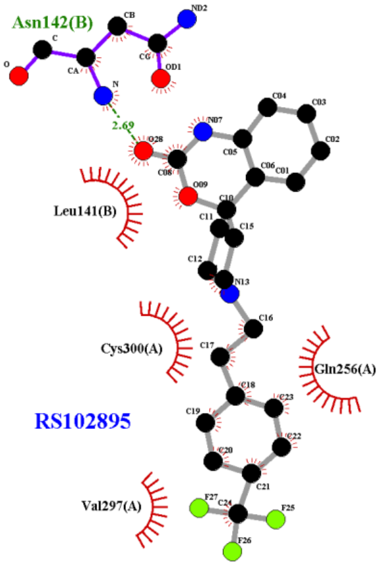


**Isomeric smiles:** C1CN(CCC12C3=CC=CC=C3NC(=O)O2)CCC4=CC=C(C=C4)C(F)(F)F

**Refinement 2Fo-Fc map**, 0.5 σ-level

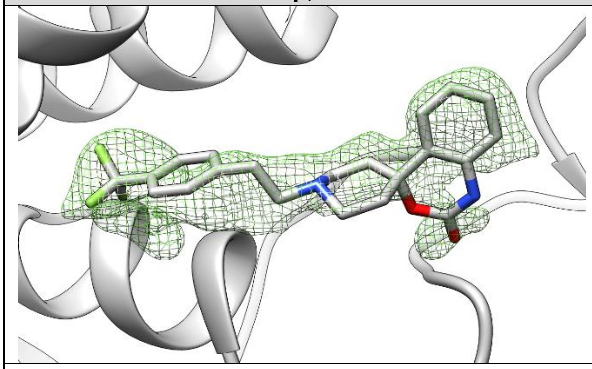

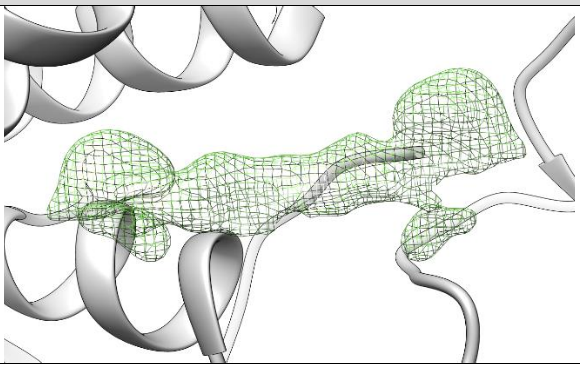

| | |
|---|---|
| **PDB structure code:** 6ABU | **PDB Ligand code:** R6Q |
| **Resolution:** 1.6 Å | **$R_{work}$/$R_{free}$:** 0.18/0.22 |
| **Occupancy:** 0.48 + 0.48 | **Ligand RSCC/RSR:** 0.74/0.22 |
| **Biochemistry & Cell biology** | |
| **Anti viral activity $EC_{50}$:** 19.92 ± 0.21 µM | **vRNA yield $EC_{50}$:** 16.76 ± 3.65 µM |
| **Anti viral activity $EC_{90}$:** 23.29 ± 0.15 µM | **vRNA yield $EC_{90}$:** 22.3 ± 1.38 µM |
| **Cytotoxicity $CC_{50}$:** 51.5 ± 0.62 µM | **SI CC50/EC50:** 2.59 |
| **Native MS adduct saturation:** 3% (94/6/0) | **Native MS adduct size:** 390 Da |

### Hit#12: AT7519 bound to SARS-CoV-2 MPro, PDB: 7AGA

**Synonyms:** AT7519, 844442-38-2

**PubChem CID:** 11338033

**CAS:** 844442-38-2

**Molecular weigth:** 382.2 g/mol

**Binding type:** non-covalent

**Binding location:** allosteric site II

**Screen compound ID:** DOM_SIM_132

**Original compound:**

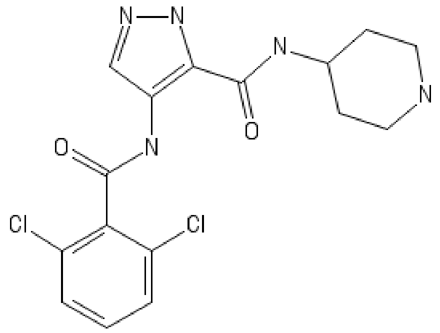


Lig-Plot interaction network

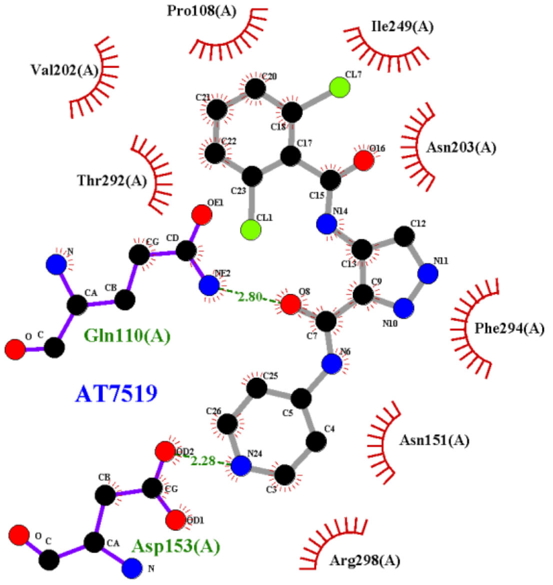


**Isomeric smiles:** C1CNCCC1NC(=O)C2=C(C=NN2)NC(=O)C3=C(C=CC=C3Cl)Cl

**Refinement 2Fo-Fc map**, 0.6 σ-level

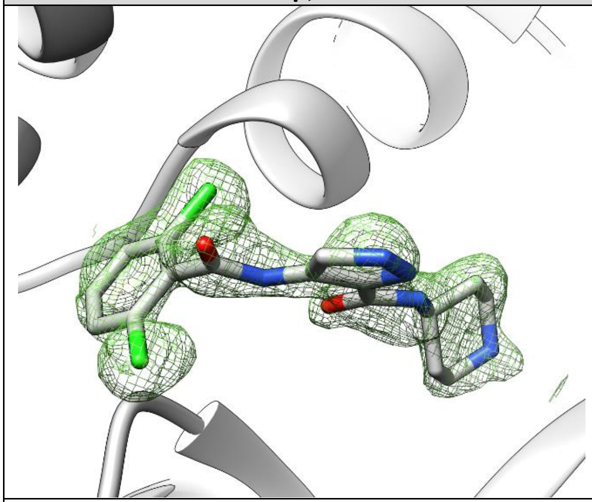

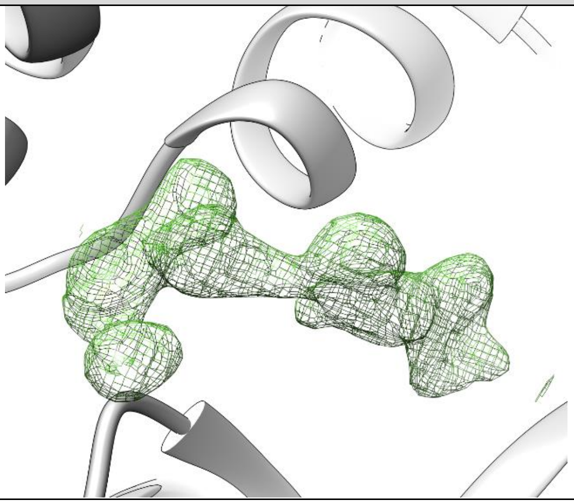

| | |
|---|---|
| **PDB structure code:** 7AGA | **PDB Ligand code:** LZE |
| **Resolution:** 1.68 Å | **$R_{work}$/$R_{free}$:** 0.19/0.23 |
| **Occupancy:** 0.72 | **Ligand RSCC/RSR:** 0.80/0.16 |

#### Biochemistry & Cell biology

| | |
|---|---|
| **Anti viral activity $EC_{50}$:** 27.5 ± 2.83 µM | **vRNA yield $EC_{50}$:** 43.66 ± 6.32 µM |
| **Anti viral activity $EC_{90}$:** 45.47 ± 2.64 µM | **vRNA yield $EC_{90}$:** 74.93 ± 5.39 µM |
| **Cytotoxicity $CC_{50}$:** > 100 µM | **SI CC50/EC50:** > 3.64 |
| **Native MS adduct saturation:** 21% (65/28/7) | **Native MS adduct size:** 382 Da |

### Hit#13: Zinc pyrithion bound to SARS-CoV-2 MPro, PDB: 6YT8

**Synonyms:** Zinc pyrithione, Zinc Omadine,

**PubChem CID:** 26041

**CAS:** 13463-41-7

**Molecular weigth:** 317.7 g/mol

**Binding type:** Covalent, complex bond

**Binding location:** Cystein 145, Histidine 41

**Screen compound ID:** SPE_K16136380

**Original compound:**

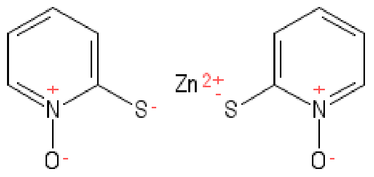


Lig-Plot interaction network

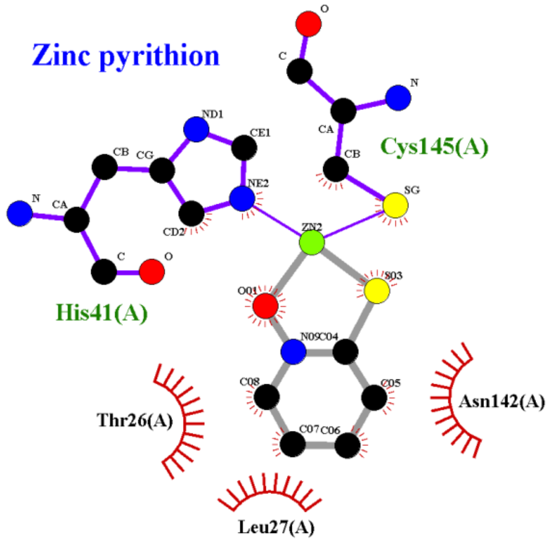


**Isomeric smiles:** C1=CC=[N+](C(=C1)[S-])[O-].C1=CC=[N+](C(=C1)[S-])[O-].[Zn+2]

**Refinement 2Fo-Fc map**, 1.4 σ-level

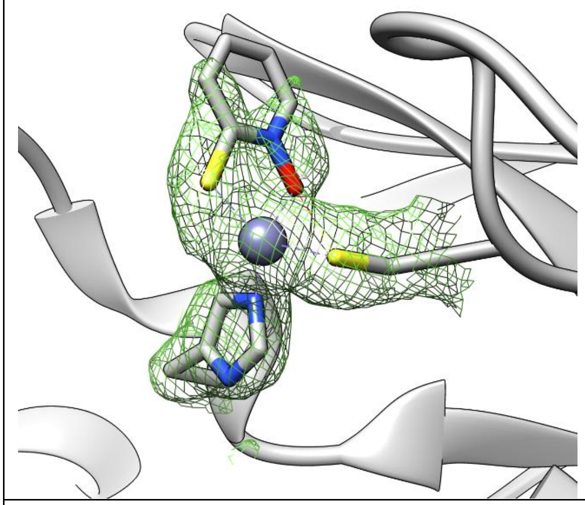

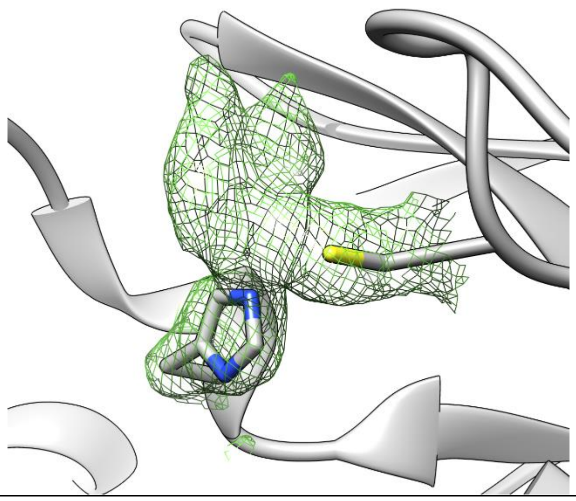

| | |
|---|---|
| **PDB structure code:** 6YT8 | **PDB Ligand code:** PK8 |
| **Resolution:** 2.05 Å | **$R_{work}$/$R_{free}$:** 0.20/0.23 |
| **Occupancy:** 0.83 | **Ligand RSCC/RSR:** 0.96/0.16 |

**Biochemistry & Cell biology**

| | |
|---|---|
| **Anti viral activity $EC_{50}$:** N.a. | **vRNA yield $EC_{50}$:** N.a. |
| **Anti viral activity $EC_{90}$:** N.a. | **vRNA yield $EC_{90}$:** N.a. |
| **Cytotoxicity $CC_{50}$:** N.a. | **SI CC50/EC50:** N.a. |
| **Native MS adduct saturation:** 72% (11/34/55) | **Native MS adduct size:** 128 Da |

, Deutsches Elektronen Synchrotron, Notkestraße 85, 22607 Hamburg

### Hit#14: TH-302 bound to SARS-CoV-2 MPro, PDB: 7AWS

**Synonyms:** Evofosfamide, TH-302, 918633-87-1, TH302

**PubChem CID:** 11984561

**CAS:** 918633-87-1

**Molecular weigth:** 449.04 g/mol

**Binding type:** covalent, thioether

**Binding location:** Cystein 145

**Screen compound ID:** SPE_K20958582

**Original compound:**

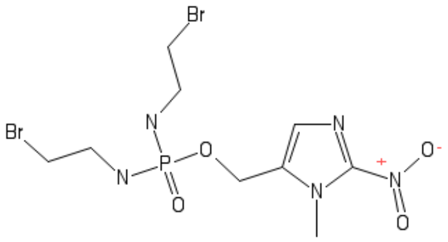

Lig-Plot interaction network

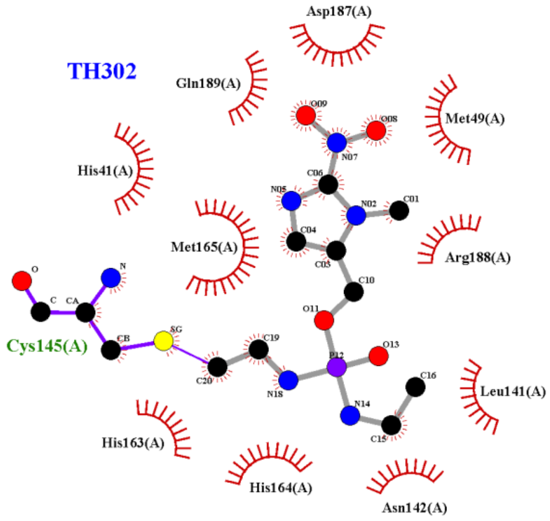


**Isomeric smiles:** CN1C(=CN=C1[N+](=O)[O-])COP(=O)(NCCBr)NCCBr

**Refinement 2Fo-Fc map**, 1.0 σ-level, **PanDDA event map BDC 0.18** , 2.0 σ-level

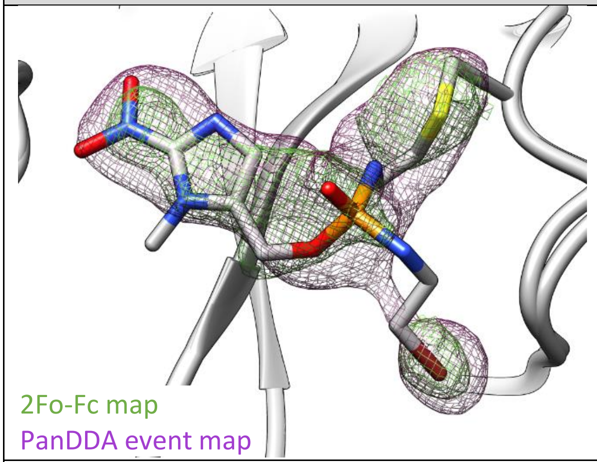


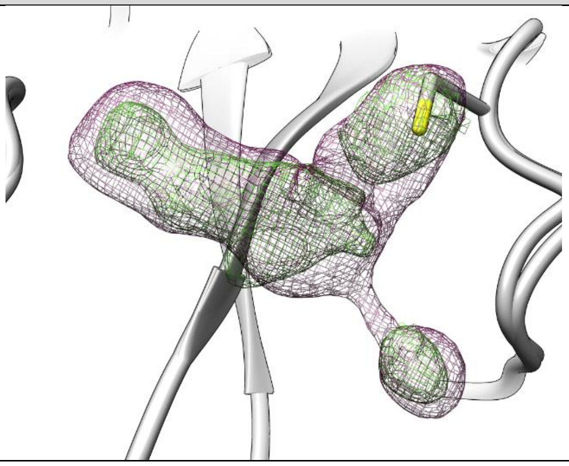

| | |
|---|---|
| **PDB structure code:** 7AWS | **PDB Ligand code:** S8E |
| **Resolution:** 1.81 Å | **$R_{work}$/$R_{free}$:** 0.20/0.24 |
| **Occupancy:** 0.56 | **Ligand RSCC/RSR:** 0.64/0.27 |

**Biochemistry & Cell biology**

| | |
|---|---|
| **Anti viral activity $EC_{50}$:** N.a. | **vRNA yield $EC_{50}$:** N.a. |
| **Anti viral activity $EC_{90}$:** N.a. | **vRNA yield $EC_{90}$:** N.a. |
| **Cytotoxicity $CC_{50}$:** N.a. | **SI CC50/EC50:** N.a. |
| **Native MS adduct saturation:** 25% (59/30/7) | **Native MS adduct size:** 365 Da |

| **Hit#15: Methazolamide bound to SARS-CoV-2 MPro, not deposited** | |
|---|---|
| **Synonyms:** Methazolamide, Neptazane, Methenamide, Neptazaneat<br>**PubChem CID:** 4100<br>**CAS:** 554-57-4<br>**Molecular weigth:** 236.3 g/mol<br>**Binding type:** binding mode undefined<br>**Binding location:** active site pocket<br>**Screen compound ID:** SPE_K13356952<br>**Original compound:** 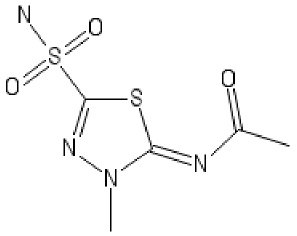 | Lig-Plot interaction network 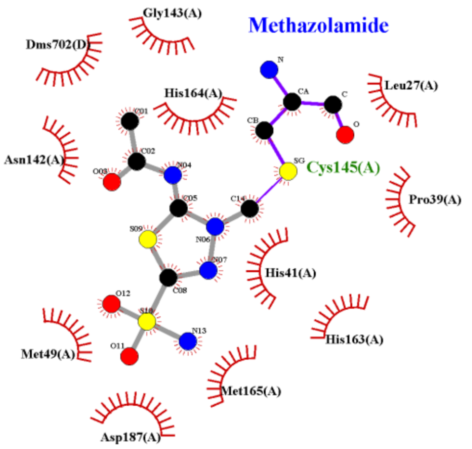 |
| **Isomeric smiles:** CC(=O)N=C1N(N=C(S1)S(=O)(=O)N)C | |
| **PanDDA event map BDC 0.18**, 1.2 σ-level | |
| 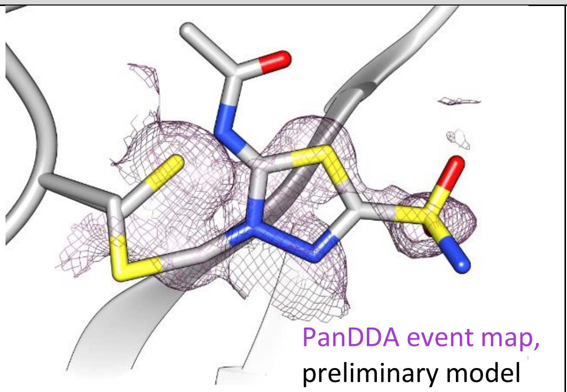 | 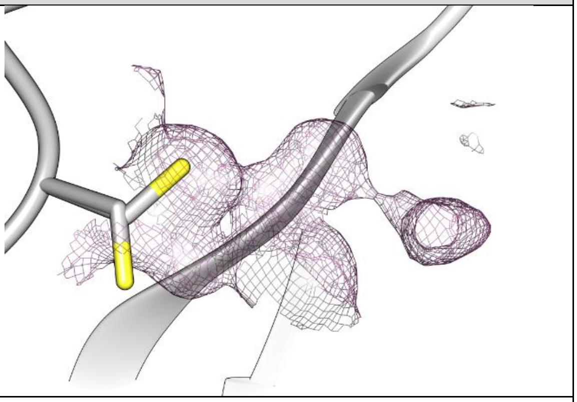 |
| **PDB structure code:** - | **PDB Ligand code:** - |
| **Resolution ohh my god Resolution:** 1.65 Å | **$R_{work}$/$R_{free}$:** - |
| **Occupancy:** - | **Ligand RSCC/RSR:** - |
| **Biochemistry & Cell biology** | |
| **Anti viral activity $EC_{50}$:** N.a. | **vRNA yield $EC_{50}$:** N.a. |
| **Anti viral activity $EC_{90}$:** N.a. | **vRNA yield $EC_{90}$:** N.a. |
| **Cytotoxicity $CC_{50}$:** N.a. | **SI CC50/EC50:** N.a. |
| **Native MS adduct saturation:** - | **Native MS adduct size:** N.a. |

#### Hit#16: Fusidic acid bound to SARS-CoV-2 MPro, PDB: 7A1U

| | |
|---|---|
| **Synonyms:** Fusidic acid, Fusidine, Ramycin, Fucithalmic | Lig-Plot interaction network |
| **PubChem CID:** 3000226 | |
| **CAS:** 6990-06-3 | |
| **Molecular weigth:** 516.7 g/mol | |
| **Binding type:** non-covalent | |
| **Binding location:** active site pocket | |
| **Screen compound ID:** SPE_A06935312 | |
| **Original compound:** | |

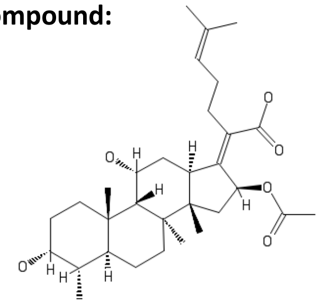

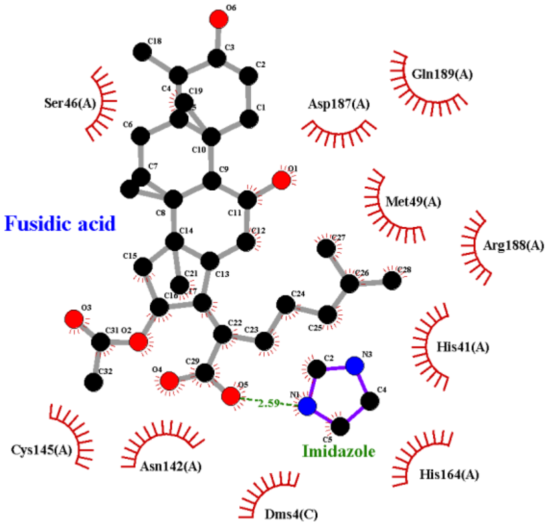


**Isomeric smiles:**
C[C@H]1[C@@H]2CC[C@]3([C@H]([C@]2(CC[C@H]1O)C)[C@@H](C[C@@H]\4[C@@]3(C[C@@H](/C4=C(/CCC=C(C)C)\C(=O)O)OC(=O)C)C)O)C

**Refinement 2Fo-Fc map**, 0.7 σ-level

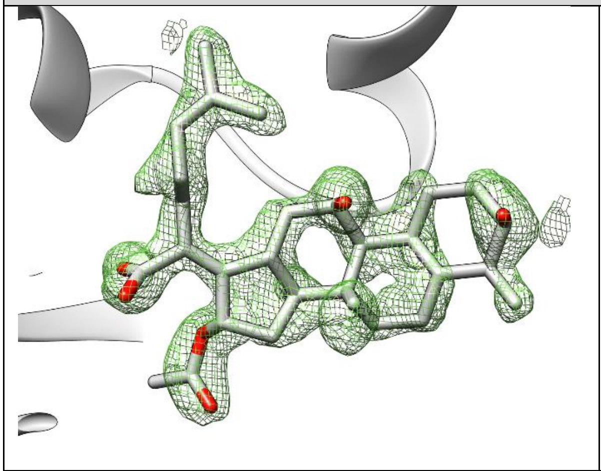

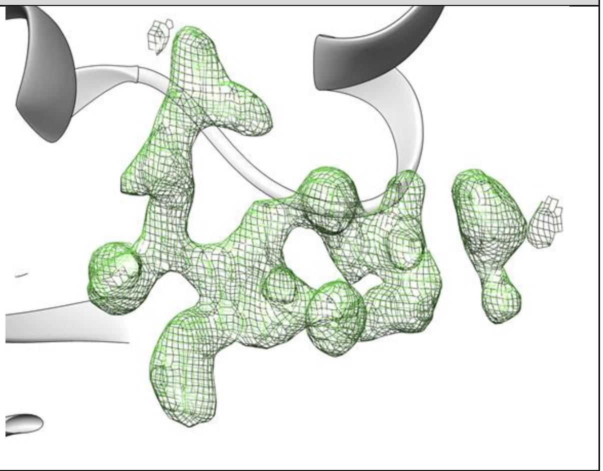

| | |
|---|---|
| **PDB structure code:** 7A1U | **PDB Ligand code:** FUA |
| **Resolution:** 1.8 Å | **$R_{work}$/$R_{free}$:** 0.19/0.23 |
| **Occupancy:** 0.8 | **Ligand RSCC/RSR:** 0.69/0.18 |
| **Biochemistry & Cell biology** | |
| **Anti viral activity $EC_{50}$:** N.a. | **vRNA yield $EC_{50}$:** N.a. |
| **Anti viral activity $EC_{90}$:** N.a. | **vRNA yield $EC_{90}$:** N.a. |
| **Cytotoxicity $CC_{50}$:** N.a. | **SI CC50/EC50:** N.a. |
| **Native MS adduct saturation:** 14% (78/18/5) | **Native MS adduct size:** 516 Da |

#### Hit#17: UNC-2327 bound to SARS-CoV-2 MPro, PDB: 7AQE

**Synonyms:** UNC-2327**,** CHEMBL2325198, BDBM50427787

**PubChem CID:** 71583615

**CAS:** 5057

**Molecular weigth:** 319.38 g/mol

**Binding type:** non-covalent

**Binding location:** active site pocket

**Screen compound ID:** SPE_K60635008

**Interaction:**

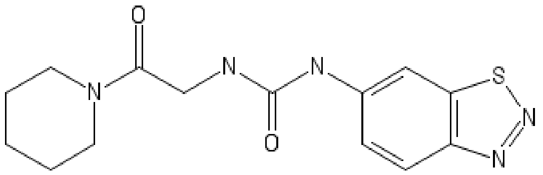

Lig-Plot interaction network

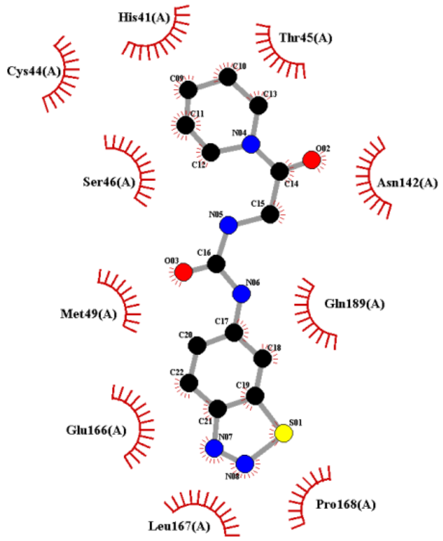


**Isomeric smiles:** C1CCN(CC1)C(=O)CNC(=O)NC2=CC3=C(C=C2)N=NS3

**Refinement 2Fo-Fc map,** 0.7 σ-level, **PanDDA event map BDC 0.18** , 1.2 σ-level

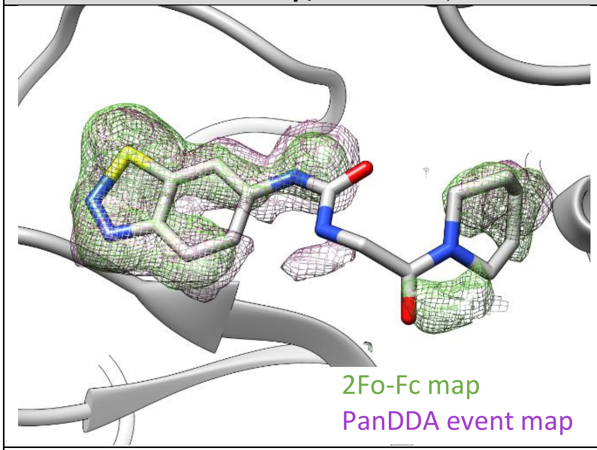


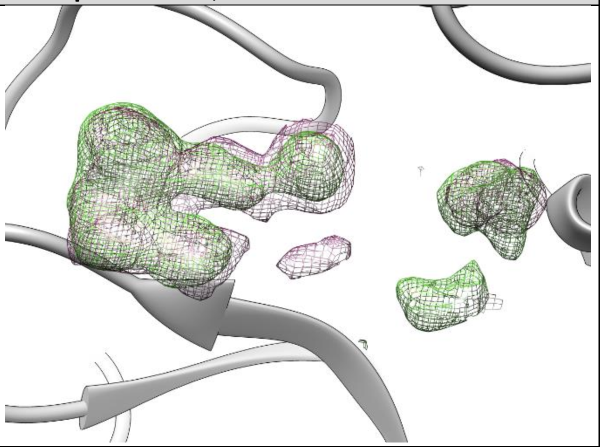

| | |
|---|---|
| **PDB structure code:** 7AQE | **PDB Ligand code:** RV5 |
| **Resolution:** 1.5 Å | $\mathbf{R_{work}/R_{free}}$: 0.20/0.24 |
| **Occupancy:** 0.82 | **Ligand RSCC/RSR:** 0.67/0.34 |

##### Biochemistry & Cell biology

| | |
|---|---|
| **Anti viral activity** $\mathbf{EC_{50}}$: N.a. | **vRNA yield** $\mathbf{EC_{50}}$: N.a. |
| **Anti viral activity** $\mathbf{EC_{90}}$: N.a. | **vRNA yield** $\mathbf{EC_{90}}$: N.a. |
| **Cytotoxicity** $\mathbf{CC_{50}}$: N.a. | **SI CC50/EC50:** N.a. |
| **Native MS adduct saturation:** 12% (76/24/0) | **Native MS adduct size:** 319 Da |

#### Hit#18: Glycitein bound to SARS-CoV-2 MPro, not deposited

| | |
|---|---|
| **Synonyms:** Glycitein, 40957-83-3, 7,4'-Dihydroxy-6-methoxyisoflavone | Lig-Plot interaction network |
| **PubChem CID:** 5317750 | |
| **CAS:** 956128-01-1 | |
| **Molecular weigth:** 284.26 g/mol | |
| **Binding type:** binding mode undefined | |
| **Binding location:** active site pocket | |
| **Screen compound ID:** SPE_K78303961 | |
| **Original compound:** | |

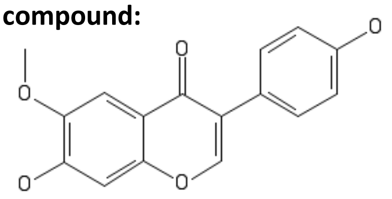

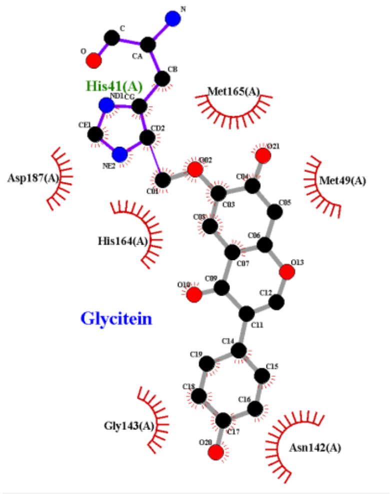


**Isomeric smiles:** COC1=C(C=C2C(=C1)C(=O)C(=CO2)C3=CC=C(C=C3)O)O

**Pandda event-map**, 1.0 σ-level

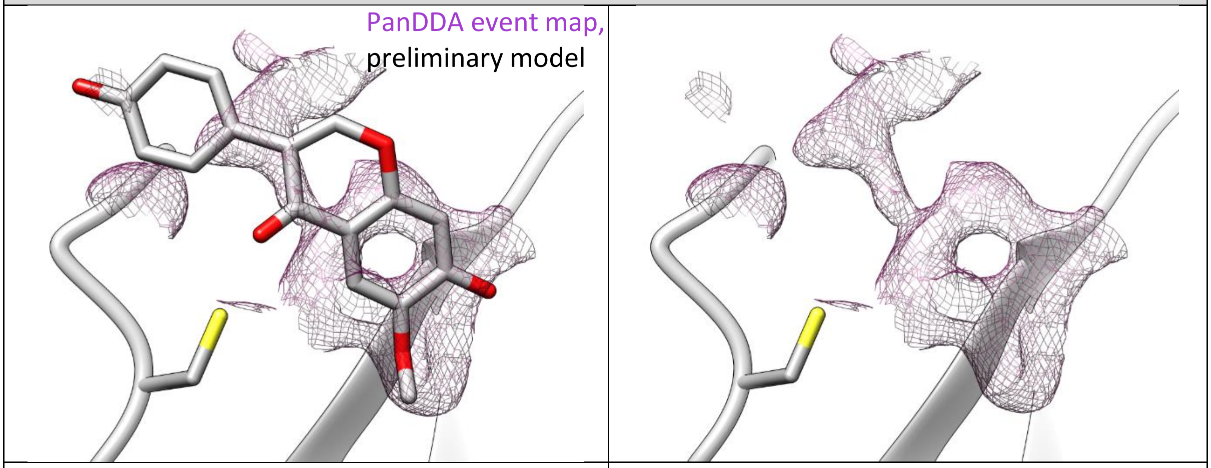


| | |
|---|---|
| **PDB structure code:** - | **PDB Ligand code:** - |
| **Resolution:** 1.7 Å | **$R_{work}$/$R_{free}$:** - |
| **Occupancy:** - | **Ligand RSCC/RSR:** - |
| **Biochemistry & Cell biology** | |
| **Anti viral activity $EC_{50}$:** N.a. | **vRNA yield $EC_{50}$:** N.a. |
| **Anti viral activity $EC_{90}$:** N.a. | **vRNA yield $EC_{90}$:** N.a. |
| **Cytotoxicity $CC_{50}$:** N.a. | **SI CC50/EC50:** N.a. |
| **Native MS adduct saturation:** - | **Native MS adduct size:** N.a. |

#### Hit#19: Bromebric-acid bound to SARS-CoV-2 MPro, not deposited

**Synonyms:** Bromebric acid, Bromebrinsaeure, Acido bromebrico, Acidum bromebricum

**PubChem CID:** 5358572

**CAS:** 21739-91-3

**Molecular weigth:** 285.09 g/mol

**Binding type:** non-covalent

**Binding location:** active site pocket

**Screen compound ID:** SPE_K17874929

**Interaction:**

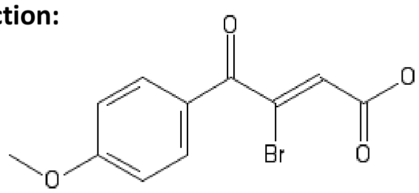


Lig-Plot interaction network

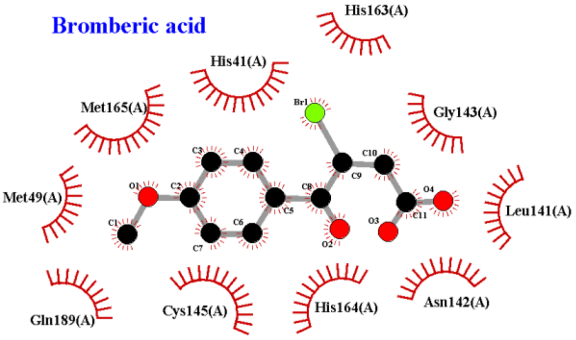


**Isomeric smiles:** COC1=CC=C(C=C1)C(=O)/C(=C\C(=O)O)/Br

**PanDDA event map BDC 0.16** , 1.2 σ-level

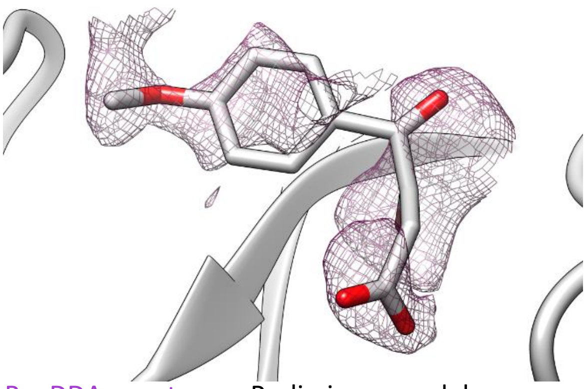

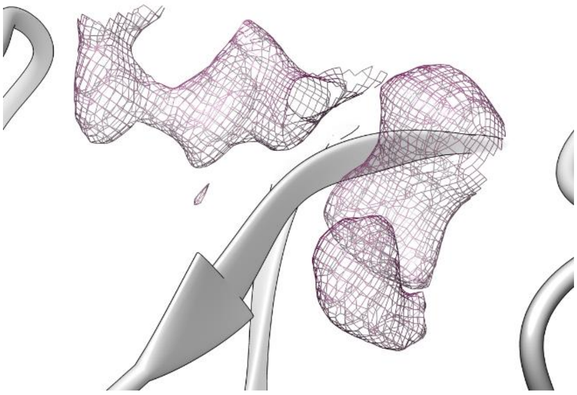

PanDDA event map, Preliminary model

| | |
|---|---|
| **PDB structure code:** - | **PDB Ligand code:** - |
| **Resolution:** 1.7 Å | **$R_{work}$/$R_{free}$:** - |
| **Occupancy:** - | **Ligand RSCC/RSR:** - |
| **Biochemistry & Cell biology** | |
| **Anti viral activity $EC_{50}$:** N.a. | **vRNA yield $EC_{50}$:** N.a. |
| **Anti viral activity $EC_{90}$:** N.a. | **vRNA yield $EC_{90}$:** N.a. |
| **Cytotoxicity $CC_{50}$:** N.a. | **SI CC50/EC50:** N.a. |
| **Native MS adduct saturation:** N.a. | **Native MS adduct size:** N.a. |

#### Hit#20: Aurothioglucose bound to SARS-CoV-2 MPro, PDB: 7ARF

**Synonyms:** Aurothioglucose, Gold thioglucose, Solganal

**PubChem CID:** 454937

**CAS:** 12192-57-3

**Molecular weigth:** 392.18 g/mol

**Binding type:** Sulfinic acid Cys145, covalent disulfide Cys156

**Binding location:** active site Cys145, surface Cys 156

**Screen compound ID:** SPE_A89825407

**Original compound:**

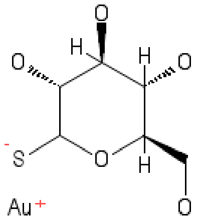


Lig-Plot interaction network

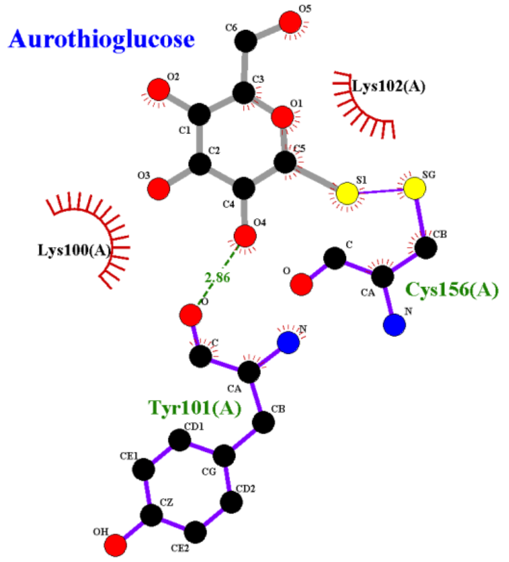


**Isomeric smiles:** C([C@@H]1[C@H]([C@@H]([C@H](C(O1)[S-])O)O)O)O.[Au+]

**Refinement 2Fo-Fc map**, 1.2 σ-level

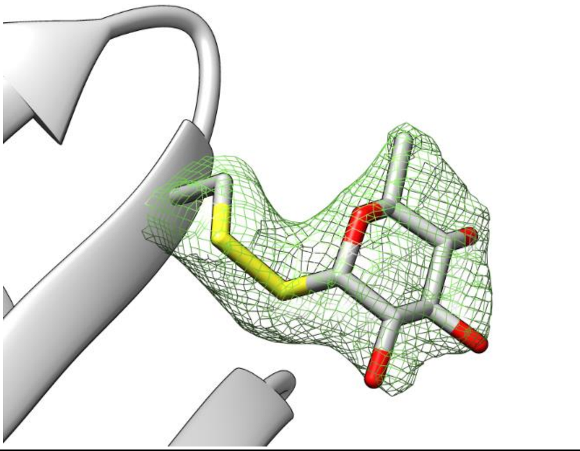

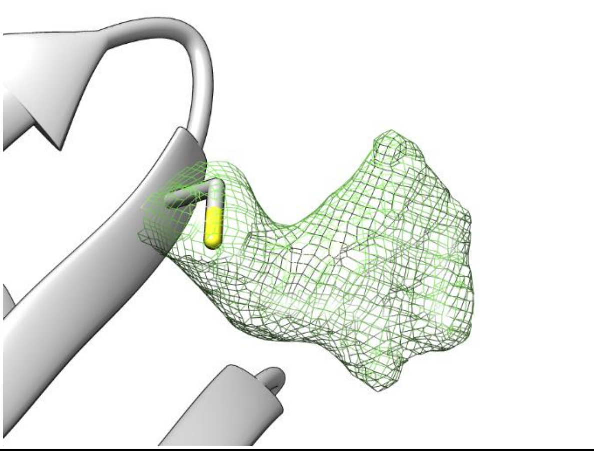

| | |
|---|---|
| **PDB structure code:** 7ARF | **PDB Ligand code:** RVW |
| **Resolution:** 2.0 Å | **$R_{work}$/$R_{free}$:** 0.21/0.25 |
| **Occupancy:** 1.0 | **Ligand RSCC/RSR:** 0.93/0.16 |

**Biochemistry & Cell biology**

| | |
|---|---|
| **Anti viral activity $EC_{50}$:** N.a. | **vRNA yield $EC_{50}$:** N.a. |
| **Anti viral activity $EC_{90}$:** N.a. | **vRNA yield $EC_{90}$:** N.a. |
| **Cytotoxicity $CC_{50}$:** N.a. | **SI CC50/EC50:** N.a. |
| **Native MS adduct saturation:** N.a. | **Native MS adduct size:** N.a. |

### Hit#21: Glutathione isopropyl ester bound to SARS-CoV-2 MPro, PDB: 7AX6

**Synonyms:** Glutathione monoisopropyl ester, Glutathione isopropyl ester, H-Glu(cys-gly-isopropyl ester)-OH

**PubChem CID:** 114894

**CAS:** 97451-46-2

**Molecular weigth:** 349.41 g/mol

**Binding type:** covalent, disulfide bond

**Binding location:** Cysteine 156

**Screen compound ID:** SPE_K58937086

**Original compound:**

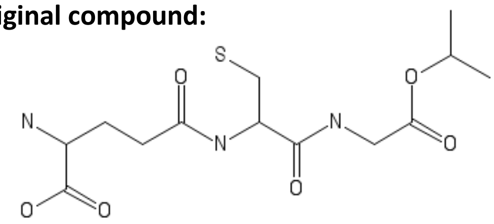

Lig-Plot interaction network

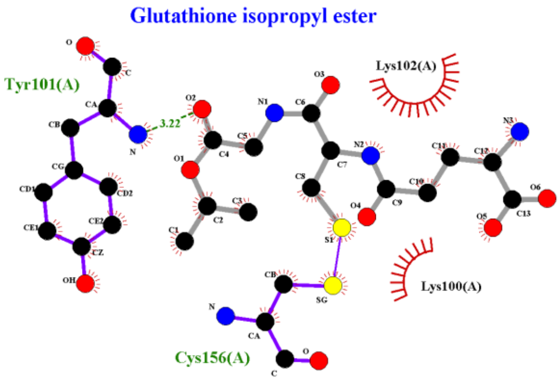


**Isomeric smiles:** CC(C)OC(=O)CNC(=O)C(CS)NC(=O)CCC(C(=O)O)N

**Refinement 2Fo-Fc map**, 0.7 σ-level

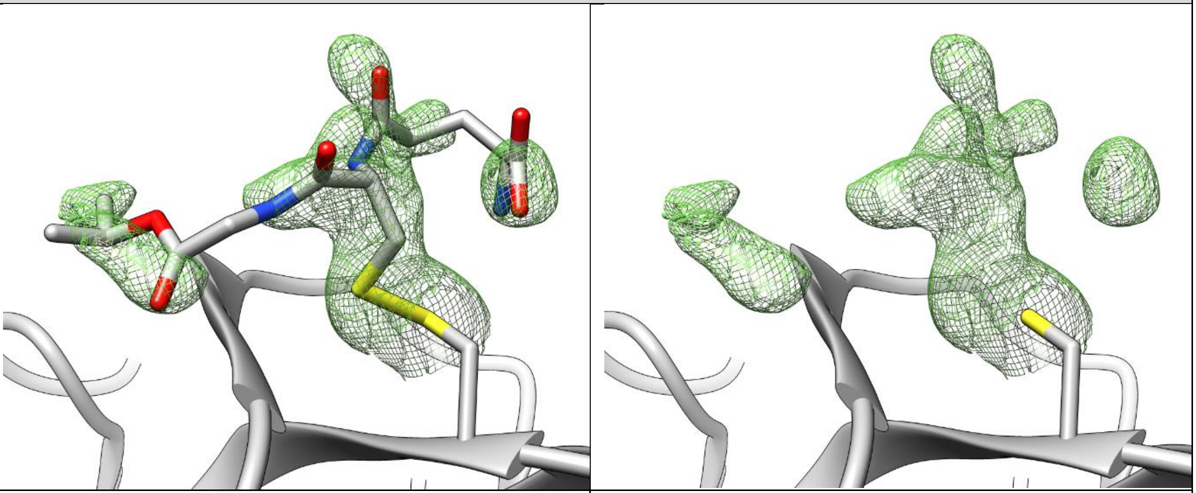

| | |
|---|---|
| **PDB structure code:** 7AX6 | **PDB Ligand code:** S8H |
| **Resolution:** 1.95 Å | **$R_{work}$/$R_{free}$:** 0.19/0.25 |
| **Occupancy:** 0.6 | **Ligand RSCC/RSR:** 0.85/0.24 |

**Biochemistry & Cell biology**

| | |
|---|---|
| **Anti viral activity $EC_{50}$:** N.a. | **vRNA yield $EC_{50}$:** N.a. |
| **Anti viral activity $EC_{90}$:** N.a. | **vRNA yield $EC_{90}$:** N.a. |
| **Cytotoxicity $CC_{50}$:** N.a. | **SI CC50/EC50:** N.a. |
| **Native MS adduct saturation:** 24% (62/28/10) | **Native MS adduct size:** 126/188 Da |

#### Hit#22: LSN-2463359 bound to SARS-CoV-2 MPro, PDB: 7AWU

**Synonyms:** N-Isopropyl-5-(pyridin-4-ylethynyl)picolinamide, LSN2463359, CHEMBL2431212

**PubChem CID:** 72551298

**CAS:** -

**Molecular weigth:** 265.31 g/mol

**Binding type:** Non-covalent

**Binding location:** Active site pocket

**Screen compound ID:** SPE_A06935312

**Original compound:**

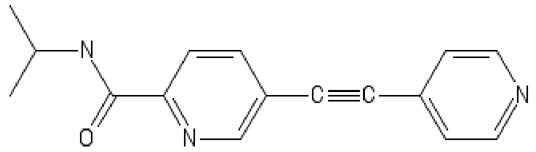


Lig-Plot interaction network

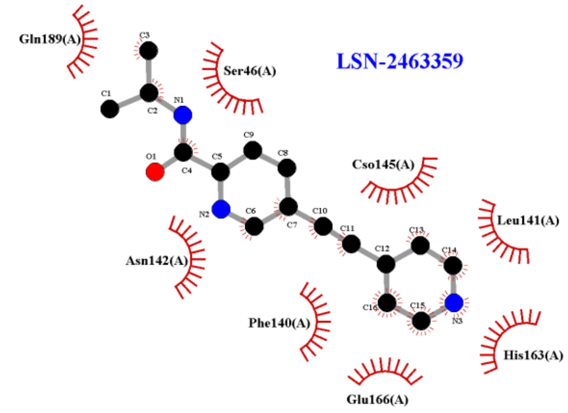


**Isomeric smiles:** CC(C)NC(=O)C1=NC=C(C=C1)C#CC2=CC=NC=C2

**Refinement 2Fo-Fc map**, 0.6 σ-level, **PanDDA event map BDC 0.18** , 1.6 σ-level

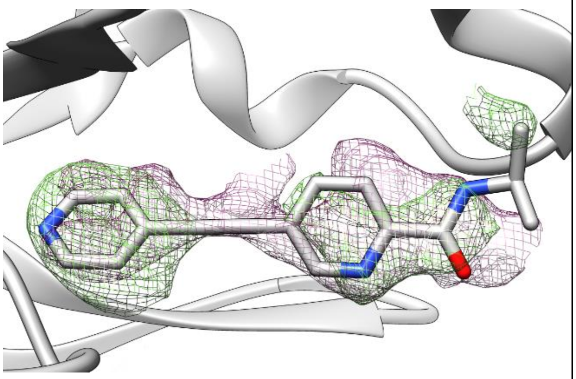

2Fo-Fc map, PanDDA event map

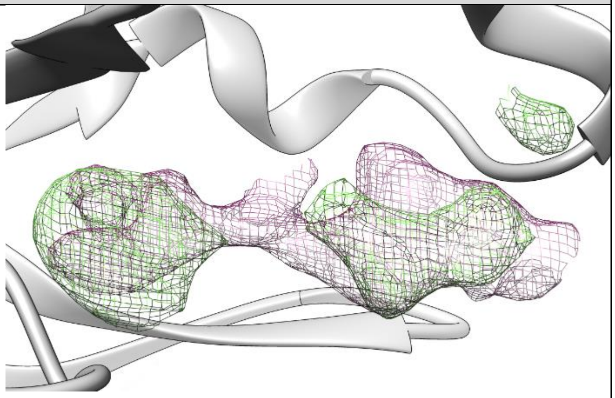

| | |
|---|---|
| **PDB structure code:** 7AWU | **PDB Ligand code:** S8B |
| **Resolution:** 2.07 Å | **$R_{work}$/$R_{free}$:** 0.21/0.26 |
| **Occupancy:** 0.75 | **Ligand RSCC/RSR:** 0.73/0.44 |
| **Biochemistry & Cell biology** | |
| **Anti viral activity $EC_{50}$:** N.a. | **vRNA yield $EC_{50}$:** N.a. |
| **Anti viral activity $EC_{90}$:** N.a. | **vRNA yield $EC_{90}$:** N.a. |
| **Cytotoxicity $CC_{50}$:** N.a. | **SI CC50/EC50:** N.a. |
| **Native MS adduct saturation:** N.a. | **Native MS adduct size:** N.a. |

### Hit#23: SUN-B-8155 bound to SARS-CoV-2 MPro, not deposited

**Synonyms:** SUN-B 8155, 345893-91-6, 5-[1-[(2-Aminophenyl)imino]ethyl]-1,6-dihydroxy-4-methyl-2(1H)-pyridone, ZINC9129

**PubChem CID:** 135484493

**CAS:** 6990-06-3

**Molecular weigth:** 273.29 g/mol

**Binding type:** binding mode undefined

**Binding location:** active site pocket

**Screen compound ID:** SPE_K79239947

**Original compound:**

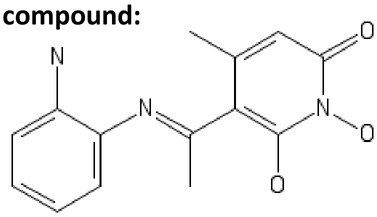


Lig-Plot interaction network

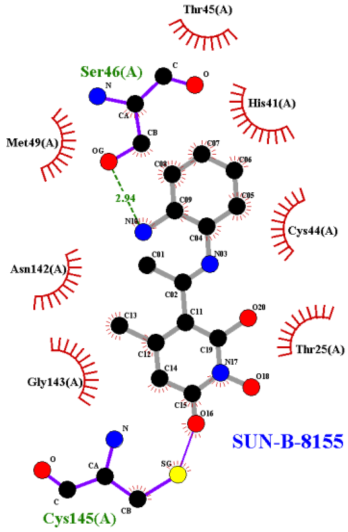


**Isomeric smiles:** CC1=CC(=O)N(C(=C1C(=NC2=CC=CC=C2N)C)O)O

**PanDDA event map BDC 0.18** , 1.2 σ-level

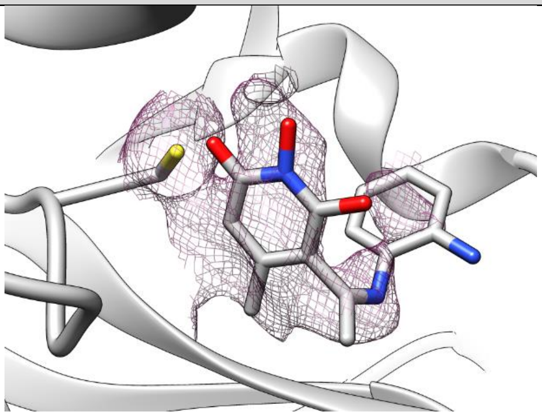

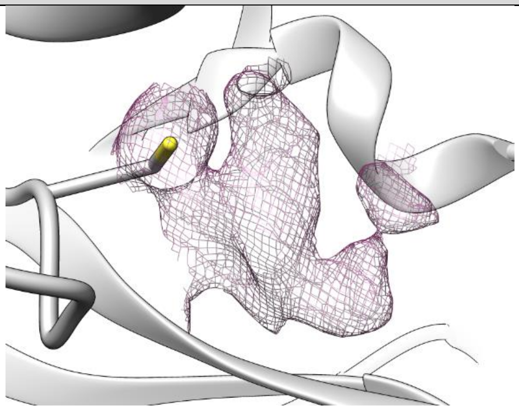

PanDDA event Map, preliminary model

| | |
|---|---|
| **PDB structure code:** - | **PDB Ligand code:** - |
| **Resolution:** 1.7 Å | **$R_{work}$/$R_{free}$:** - |
| **Occupancy:** - | **Ligand RSCC/RSR:** - |
| **Biochemistry & Cell biology** | |
| **Anti viral activity $EC_{50}$:** N.a. | **vRNA yield $EC_{50}$:** N.a. |
| **Anti viral activity $EC_{90}$:** N.a. | **vRNA yield $EC_{90}$:** N.a. |
| **Cytotoxicity $CC_{50}$:** N.a. | **SI CC50/EC50:** N.a. |
| **In vitro inhibition IC50:** N.a. | **Native mass spectrometry:** N.a. |

#### Hit#24: SEN1269 bound to SARS-CoV-2 MPro, PDB: 7AVD

**Synonyms:** SEN1269, 3-(5-(3-(dimethylamino)phenoxy)pyrimidin-2-ylamino)phenol

**PubChem CID:** 46835756

**CAS:** 956128-01-1

**Molecular weigth:** 322.36 g/mol

**Binding type:** Non-covalent

**Binding location:** Surface

**Screen compound ID:** SPE_K70614402

**Original compound:**

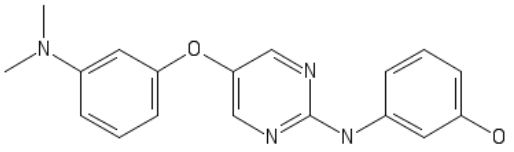

Lig-Plot interaction network

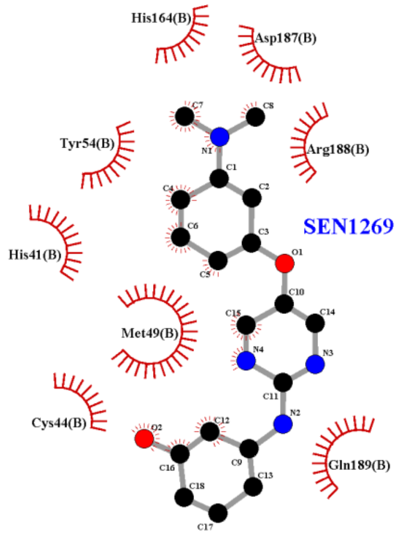


**Isomeric smiles:** CN(C)C1=CC(=CC=C1)OC2=CN=C(N=C2)NC3=CC(=CC=C3)O

**Refinement 2Fo-Fc map**, 0.7 σ-level

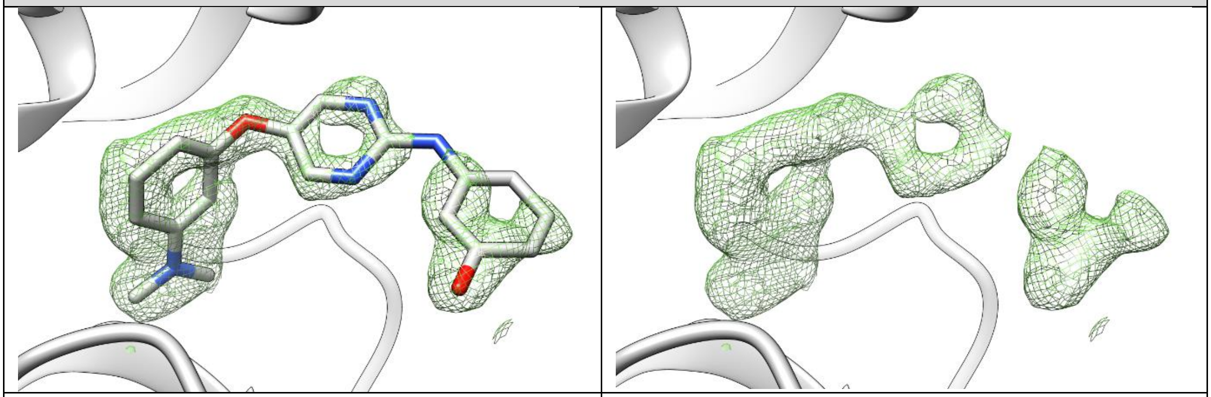

| | |
|---|---|
| **PDB structure code:** 7AVD | **PDB Ligand code:** S1W |
| **Resolution:** 1.8 Å | **$R_{work}$/$R_{free}$:** 0.19/0.24 |
| **Occupancy:** 1.0 | **Ligand RSCC/RSR:** 0.76/0.27 |

**Biochemistry & Cell biology**

| | |
|---|---|
| **Anti viral activity $EC_{50}$:** N.a. | **vRNA yield $EC_{50}$:** N.a. |
| **Anti viral activity $EC_{90}$:** N.a. | **vRNA yield $EC_{90}$:** N.a. |
| **Cytotoxicity $CC_{50}$:** N.a. | **SI CC50/EC50:** N.a. |
| **In vitro inhibition IC50:** N.a. | **Native mass spectrometry:** N.a. |

### Hit#25: PD168568 bound to SARS-CoV-2 MPro, PDB: 7AMJ

**Synonyms:** PD 168568, 210688-56-5
**PubChem CID:** 56972231
**CAS:** 210688-56-5
**Molecular weigth:** 422.4 g/mol
**Binding type:** non-covalent
**Binding location:** allosteric site I
**Screen compound ID:** SPE_A82013137
**Original compound:**

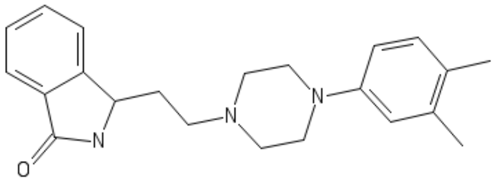

Lig-Plot interaction network

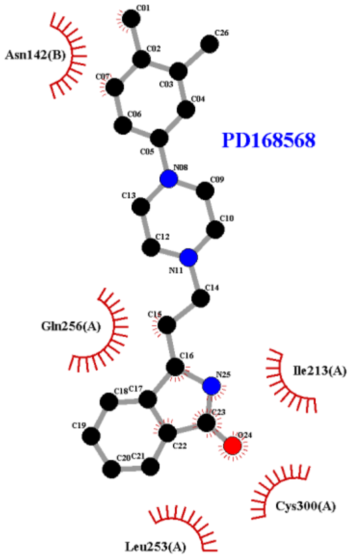


**Isomeric smiles:** CC1=C(C=C(C=C1)N2CCN(CC2)CCC3C4=CC=CC=C4C(=O)N3)C.Cl.Cl

**Refinement 2Fo-Fc map**, 0.5 σ-level

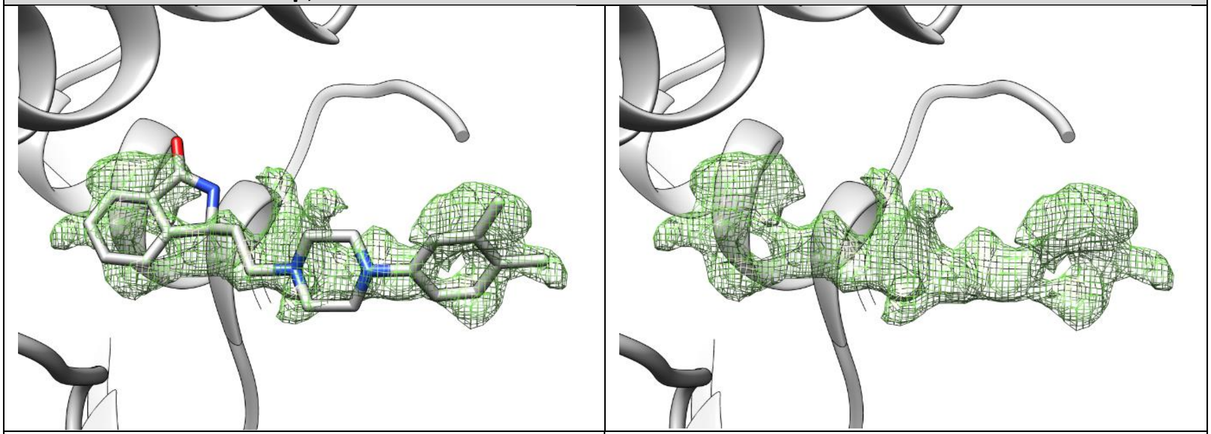

| | |
|---|---|
| **PDB structure code:** 7AMJ | **PDB Ligand code:** RMZ |
| **Resolution:** 1.59 Å | **$R_{work}/R_{free}$:** 0.17/0.21 |
| **Occupancy:** 0.42 + 0.42 | **Ligand RSCC/RSR:** 0.65/0.27 |

**Biochemistry & Cell biology**

| | |
|---|---|
| **Anti viral activity $EC_{50}$:** N.a. | **vRNA yield $EC_{50}$:** N.a. |
| **Anti viral activity $EC_{90}$:** N.a. | **vRNA yield $EC_{90}$:** N.a. |
| **Cytotoxicity $CC_{50}$:** N.a. | **SI CC50/EC50:** N.a. |
| **Native MS adduct saturation:** 11% (81/16/3) | **Native MS adduct size:** 350 Da |

#### Hit#26: Tofogliflozin bound to SARS-CoV-2 MPro, PDB: 7APH

**Synonyms:** Tofogliflozin, 903565-83-3, CSG 452, Tofogliflozin anhydrous, UNII-554245W62T

**PubChem CID:** 46908929

**CAS:** 903565-83-3

**Molecular weigth:** 386.4 g/mol

**Binding type:** non-covalent

**Binding location:** allosteric site I

**Screen compound ID:** SPE_K96344439

**Interaction:**

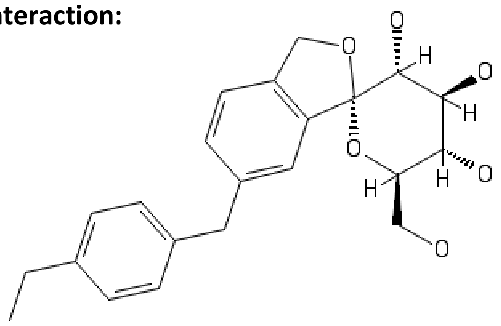

Lig-Plot interaction network

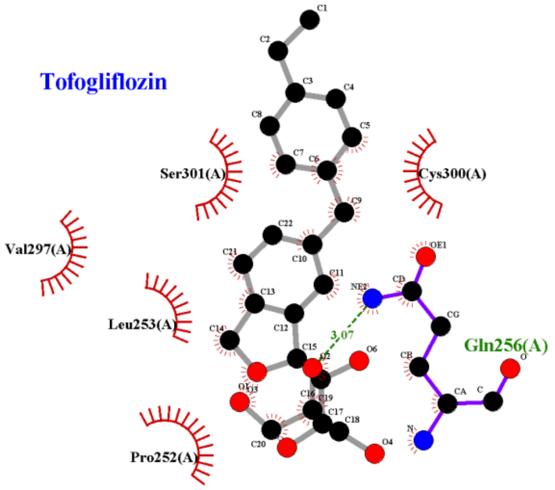


**Isomeric smiles:**
CCC1=CC=C(C=C1)CC2=CC3=C(CO[C@@]34[C@@H]([C@H]([C@@H]([C@H](O4)CO)O)O)O)C=C2

**Refinement 2Fo-Fc map**, 0.7 σ-level

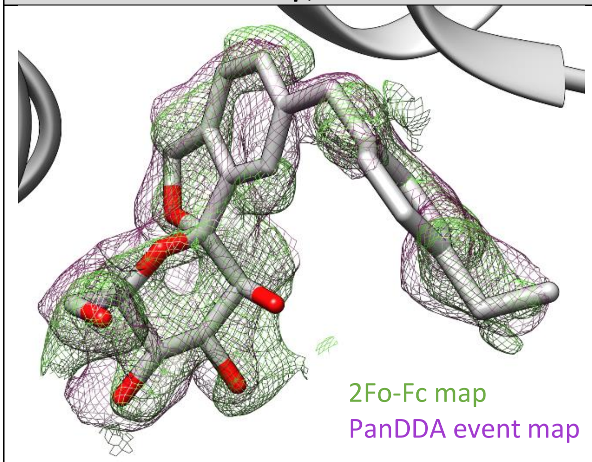


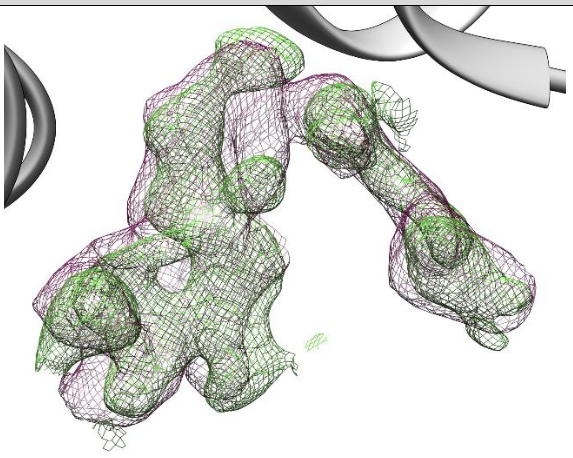

| | |
|---|---|
| **PDB structure code:** 7APH | **PDB Ligand code:** RT2 |
| **Resolution:** 1.65 Å | **$R_{work}$/$R_{free}$:** 0.23/0.27 |
| **Occupancy:** 0.5 | **Ligand RSCC/RSR:** 0.52/0.38 |

**Biochemistry & Cell biology**

| | |
|---|---|
| **Anti viral activity $EC_{50}$:** N.a. | **vRNA yield $EC_{50}$:** N.a. |
| **Anti viral activity $EC_{90}$:** N.a. | **vRNA yield $EC_{90}$:** N.a. |
| **Cytotoxicity $CC_{50}$:** N.a. | **SI CC50/EC50:** N.a. |
| **Native MS adduct saturation:** 9% (82/18/0) | **Native MS adduct size:** 380 Da |

### Hit#27: Adrafinil bound to SARS-CoV-2 MPro, PDB: 7ANS

**Synonyms:** Adrafinil, Olmifon, CRL 40028

**PubChem CID:** 3033226

**CAS:** 63547-13-7

**Molecular weigth:** 289.4 g/mol

**Binding type:** non-covalent

**Binding location:** active site pocket

**Screen compound ID:** DOM_SIM_074 & SPE_A90926615

**Interaction:**

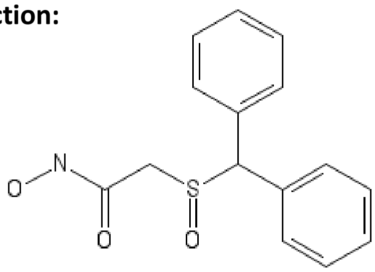


Lig-Plot interaction network

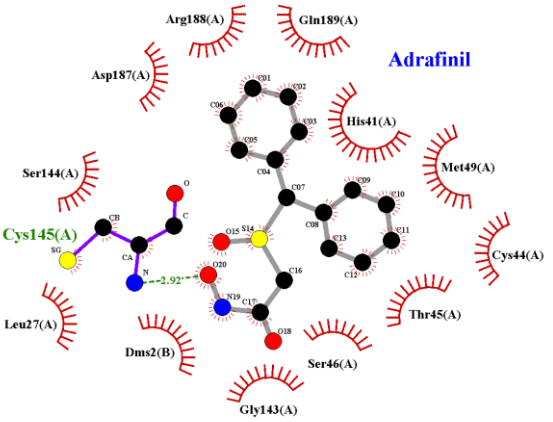


**Isomeric smiles:** C1=CC=C(C=C1)C(C2=CC=CC=C2)[S](=O)CC(=O)NO

**Refinement 2Fo-Fc map**, 0.5 σ-level

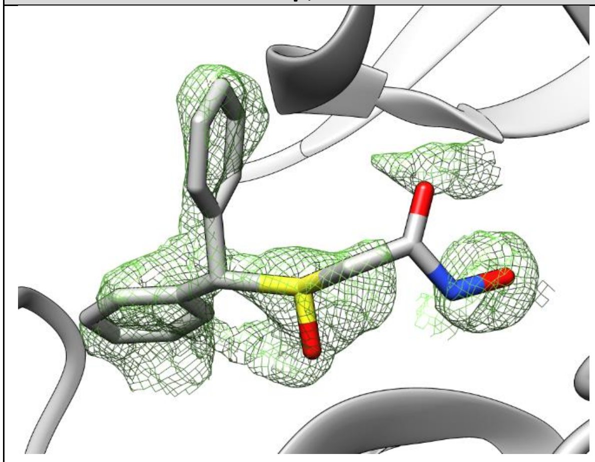

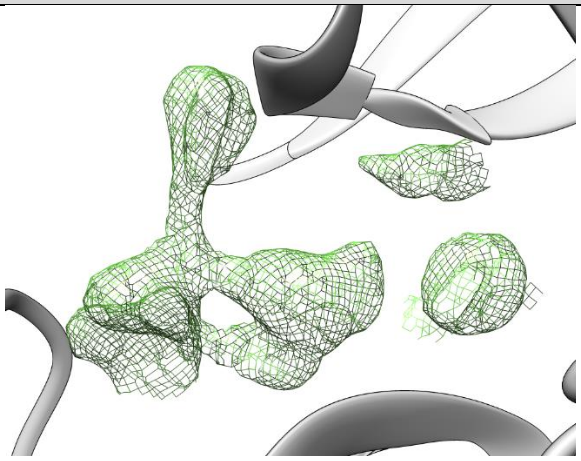

| | |
|---|---|
| **PDB structure code:** 7ANS | **PDB Ligand code:** RNW |
| **Resolution:** 1.7 Å | **$R_{work}$/$R_{free}$:** 0.18/0.21 |
| **Occupancy:** 0.65 | **Ligand RSCC/RSR:** 0.63/0.28 |
| **Biochemistry & Cell biology** | |
| **Anti viral activity $EC_{50}$:** N.a. | **vRNA yield $EC_{50}$:** N.a. |
| **Anti viral activity $EC_{90}$:** N.a. | **vRNA yield $EC_{90}$:** N.a. |
| **Cytotoxicity $CC_{50}$:** N.a. | **SI CC50/EC50:** N.a. |
| **Native MS adduct saturation:** 28% (58/28/14) | **Native MS adduct size:** 289 Da |

#### Hit#28: Tretazicar bound to SARS-CoV-2 MPro, PDB: 7AK4

| | |
|---|---|
| **Synonyms:** Tretazicar, CB 1954, 21919-05-1 | Lig-Plot interaction network |
| **PubChem CID:** 89105 | |
| **CAS:** 21919-05-1 | |
| **Molecular weigth:** 252.18 g/mol | |
| **Binding type:** non-covalent | |
| **Binding location:** active site pocket | |
| **Screen compound ID:** DOM_SIM_234 | |
| **Interaction:** 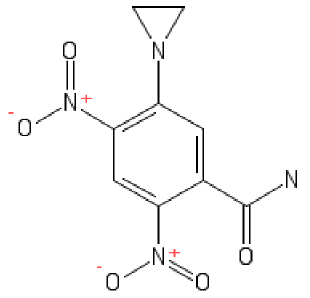 | 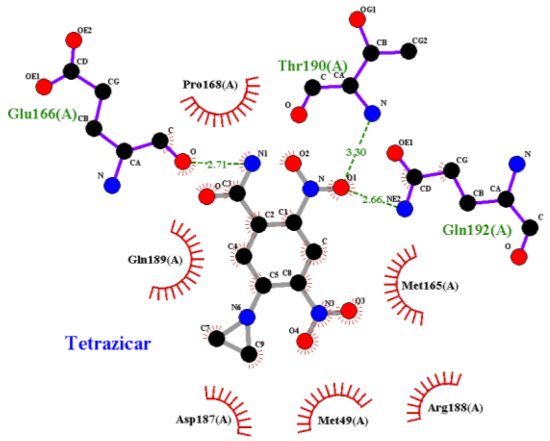 |



**Isomeric smiles:** C1CN1C2=C(C=C(C(=C2)C(=O)N)[N+](=O)[O-])[N+](=O)[O-]

**Refinement 2Fo-Fc map**, 0.7 σ-level

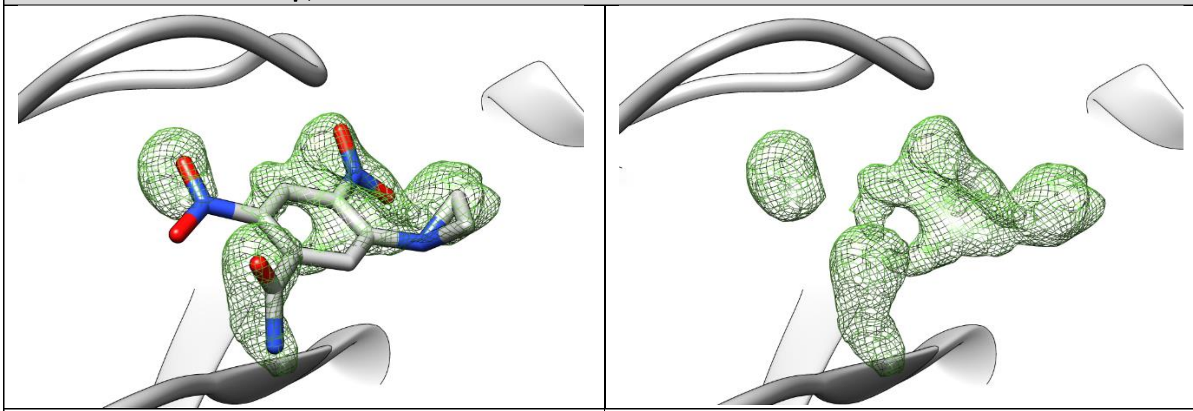

| | |
|---|---|
| **PDB structure code:** 7AK4 | **PDB Ligand code:** CB1 |
| **Resolution:** 1.63 Å | **$R_{work}$/$R_{free}$:** 0.19/0.22 |
| **Occupancy:** 0.71 | **Ligand RSCC/RSR:** 0.62/0.20 |
| **Biochemistry & Cell biology** | |
| **Anti viral activity $EC_{50}$:** N.a. | **vRNA yield $EC_{50}$:** N.a. |
| **Anti viral activity $EC_{90}$:** N.a. | **vRNA yield $EC_{90}$:** N.a. |
| **Cytotoxicity $CC_{50}$:** N.a. | **SI CC50/EC50:** N.a. |
| **Native MS adduct saturation:** - | **Native MS adduct size:** N.a. |

, Deutsches Elektronen Synchrotron, Notkestraße 85, 22607 Hamburg

### Hit#29: Climbazole bound to SARS-CoV-2 MPro, PDB: 7AOL

| | |
|---|---|
| **Synonyms:** Climbazole, 38083-17-9, Baypival, Baysan | Lig-Plot interaction network |
| **PubChem CID:** 37907 | |
| **CAS:** 38083-17-9 | |
| **Molecular weigth:** 292.76 g/mol | |
| **Binding type:** non-covalent | |
| **Binding location:** surface | |
| **Screen compound ID:** DOM_SIM_072 | |
| **Interaction:** 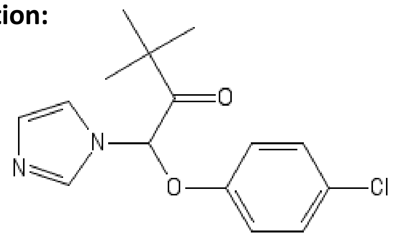 | 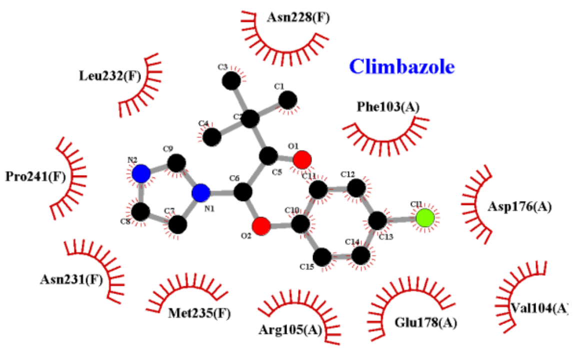  |

**Isomeric smiles:** CC(C)(C)C(=O)C(N1C=CN=C1)OC2=CC=C(C=C2)Cl

**Refinement 2Fo-Fc map**, 0.5 σ-level

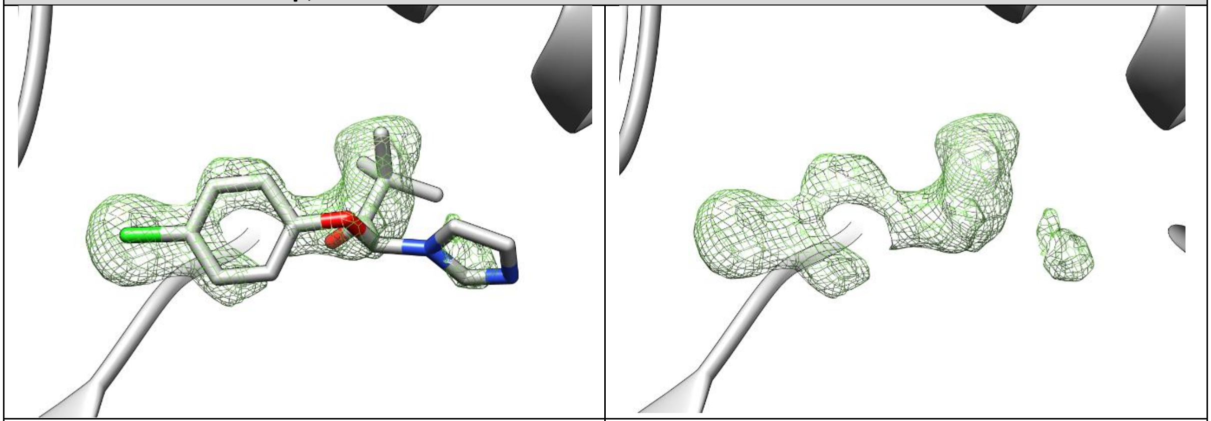

| | |
|---|---|
| **PDB structure code:** 7AOL | **PDB Ligand code:** RQH |
| **Resolution:** 1.47 Å | **$R_{work}$/$R_{free}$:** 0.17/0.19 |
| **Occupancy:** 0.65 | **Ligand RSCC/RSR:** 0.86/0.26 |
| **Biochemistry & Cell biology** | |
| **Anti viral activity $EC_{50}$:** N.a. | **vRNA yield $EC_{50}$:** N.a. |
| **Anti viral activity $EC_{90}$:** N.a. | **vRNA yield $EC_{90}$:** N.a. |
| **Cytotoxicity $CC_{50}$:** N.a. | **SI CC50/EC50:** N.a. |
| **Native MS adduct saturation:** - | **Native MS adduct size:** N.a. |

#### Hit#30: Dexrazoxane bound to SARS-CoV-2 MPro, not deposited

**Synonyms:** Dexrazoxane, 24584-09-6, Zinecard, Cardioxane

**PubChem CID:** 71384

**CAS:** 24584-09-6

**Molecular weigth:** 268.27 g/mol

**Binding type:** binding mode undefined

**Binding location:** active site pocket

**Screen compound ID:** DOM_SIM_486

**Original compound:**

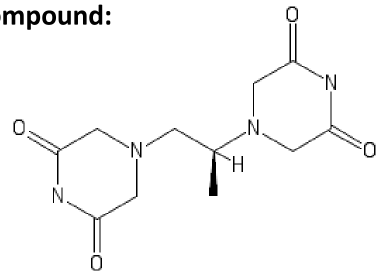

Lig-Plot interaction network

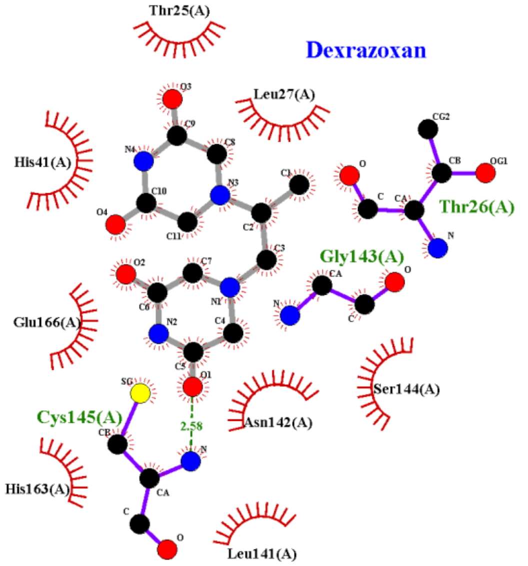


**Isomeric smiles:** C[C@@H](CN1CC(=O)NC(=O)C1)N2CC(=O)NC(=O)C2

**PanDDA event map BDC 0.18** , 1.2 σ-level

PanDDA event map,
Preliminary model

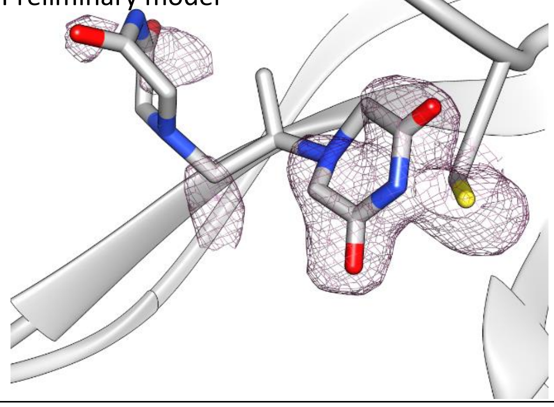

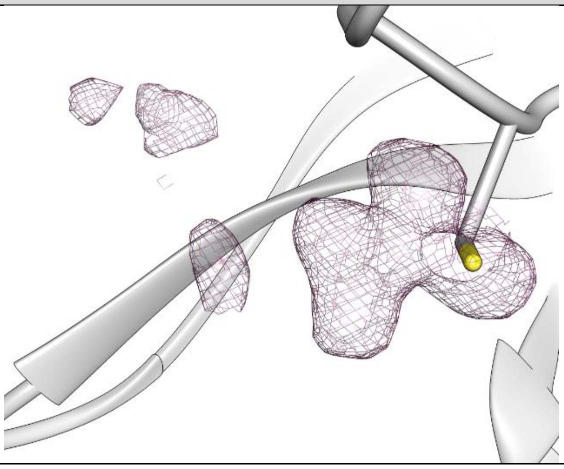

| | |
|---|---|
| **PDB structure code:** - | **PDB Ligand code:** - |
| **Resolution:** 1.7 Å | $\mathbf{R_{work}/R_{free}}$: - |
| **Occupancy:** - | **Ligand RSCC/RSR:** - |
| **Biochemistry & Cell biology** | |
| **Anti viral activity $EC_{50}$:** N.a. | **vRNA yield $EC_{50}$:** N.a. |
| **Anti viral activity $EC_{90}$:** N.a. | **vRNA yield $EC_{90}$:** N.a. |
| **Cytotoxicity $CC_{50}$:** N.a. | **SI CC50/EC50:** N.a. |
| **Native MS adduct saturation:** 30% (56/29/15) | **Native MS adduct size:** 268 Da |

#### Hit#31: AR-42 bound to SARS-CoV-2 MPro, PDB: 7AXO

| | |
|---|---|
| **Synonyms:** AR-42, OSU-HDAC42 | Lig-Plot interaction network |
| **PubChem CID:** 6918848 | |
| **CAS:** 935881-37-1 | |
| **Molecular weigth:** 312.4 g/mol | |
| **Binding type:** non-covalent | |
| **Binding location:** surface | |
| **Screen compound ID:** DOM_SIM_538 | |
| **Interaction:** | |

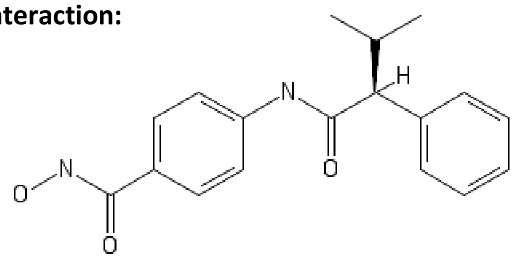


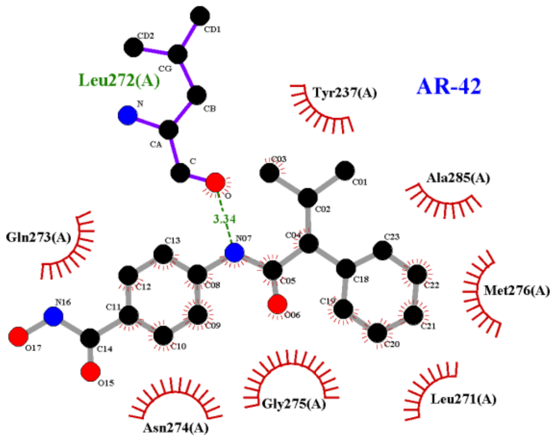


**Isomeric smiles:** CC(C)[C@@H](C1=CC=CC=C1)C(=O)NC2=CC=C(C=C2)C(=O)NO

**Refinement 2Fo-Fc map**, 0.7 σ-level, **PanDDA event map BDC 0.18** , 1.2 σ-level

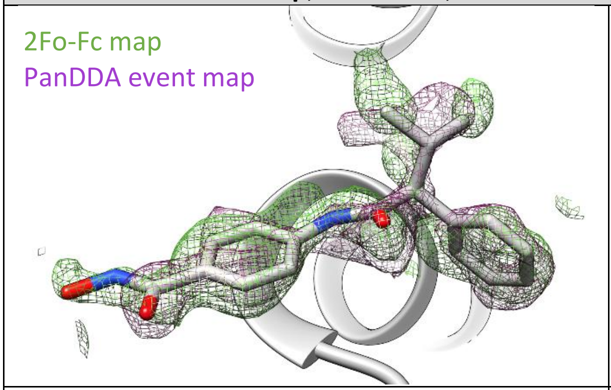


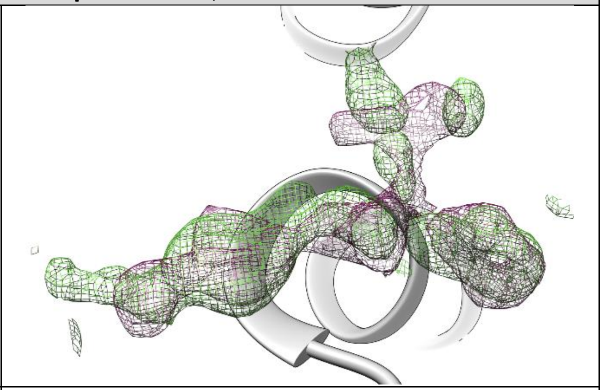

| | |
|---|---|
| **PDB structure code:** 7AXO | **PDB Ligand code:** QCP |
| **Resolution:** 1.65 Å | $\mathbf{R_{work}/R_{free}}$**:** 0.17/0.20 |
| **Occupancy:** 0.79 | **Ligand RSCC/RSR:** 0.67/0.22 |
| **Biochemistry & Cell biology** | |
| **Anti viral activity $EC_{50}$:** N.a. | **vRNA yield $EC_{50}$:** N.a. |
| **Anti viral activity $EC_{90}$:** N.a. | **vRNA yield $EC_{90}$:** N.a. |
| **Cytotoxicity $CC_{50}$:** N.a. | **SI CC50/EC50:** N.a. |
| **Native MS adduct saturation:** 13% (78/18/4) | **Native MS adduct size:** 580 Da |

### Hit#32: Clonidine bound to SARS-CoV-2 MPro, PDB: 7AWW

**Synonyms:** Clonidine

**PubChem CID:** 5358572

**CAS:** 21739-91-3

**Molecular weigth:** 285.09 g/mol

**Binding type:** non-covalent

**Binding location:** active site

**Screen compound ID:** SPE_K98530306

**Interaction:**

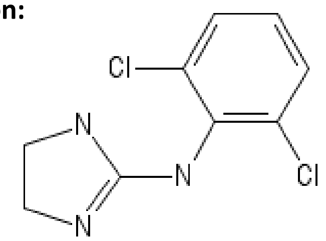


Lig-Plot interaction network

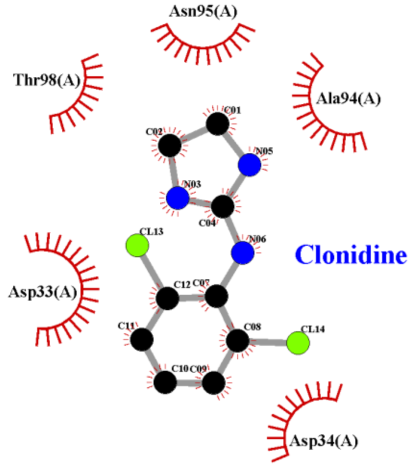


**Isomeric smiles:** C1CN=C(N1)NC2=C(C=CC=C2Cl)Cl

**Refinement 2Fo-Fc map**, 0.7 σ-level, **PanDDA event map BDC 0.18** , 1.6 σ-level

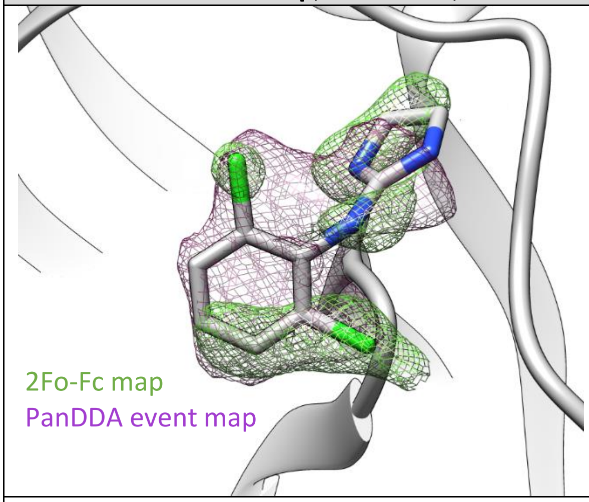


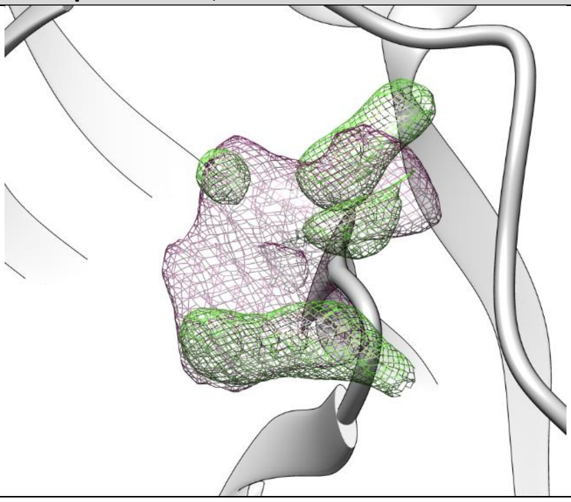

| | |
|---|---|
| **PDB structure code:** 7AWW | **PDB Ligand code:** CLU |
| **Resolution:** 1.65 Å | **$R_{work}$/$R_{free}$:** 0.20/0.24 |
| **Occupancy:** 0.80 | **Ligand RSCC/RSR:** 0.40/0.39 |
| **Biochemistry & Cell biology** | |
| **Anti viral activity $EC_{50}$:** N.a. | **vRNA yield $EC_{50}$:** N.a. |
| **Anti viral activity $EC_{90}$:** N.a. | **vRNA yield $EC_{90}$:** N.a. |
| **Cytotoxicity $CC_{50}$:** N.a. | **SI CC50/EC50:** N.a. |
| **Native MS adduct saturation:** - | **Native MS adduct size:** N.a. |

#### Hit#33: Tegafur bound to SARS-CoV-2 MPro, PDB: 7AWR

**Synonyms:** Tegafur, Ftorafur, 17902-23-7, Futraful, Fluorofur

**PubChem CID:** 5386

**CAS:** 6990-06-3

**Molecular weigth:** 200.17 g/mol

**Binding type:** non-covalent

**Binding location:** surface

**Screen compound ID:** SPE_A06935312

**Original compound:**

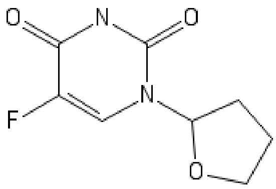


Lig-Plot interaction network

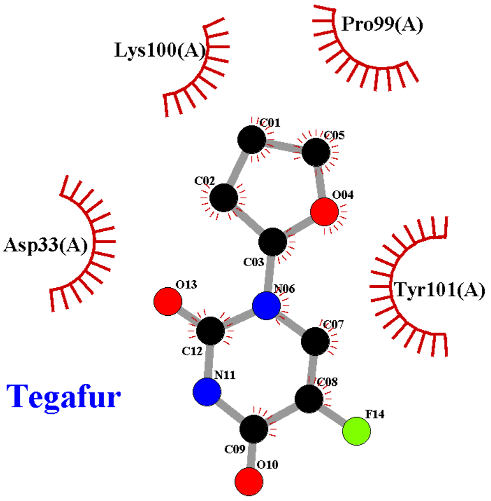


**Isomeric smiles:** C1CC(OC1)N2C=C(C(=O)NC2=O)F

**Refinement 2Fo-Fc map**, 0.6 σ-level, **PanDDA event map BDC 0.18** , 1.7 σ-level

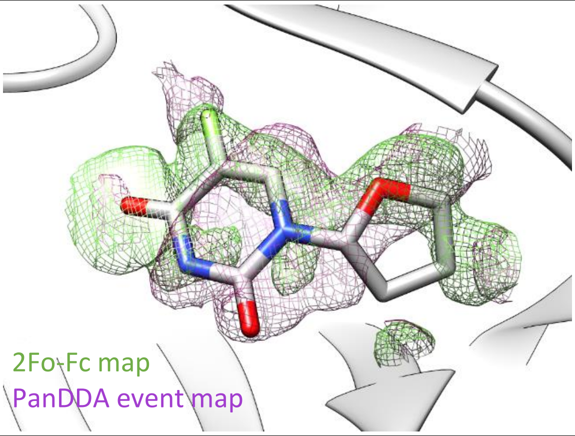


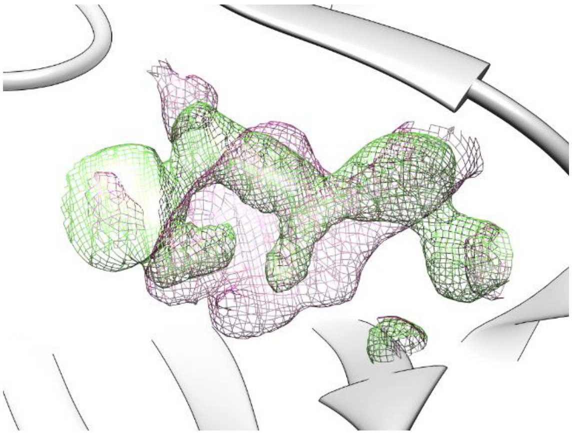

| | |
|---|---|
| **PDB structure code:** 7AWR | **PDB Ligand code:** S7W |
| **Resolution:** 1.65 Å | **$R_{work}$/$R_{free}$:** 0.18/0.22 |
| **Occupanc** | **Ligand RSCC/RSR:** 0.50/0.27 |
| **Biochemistry & Cell biology** | |
| **Anti viral activity $EC_{50}$:** N.a. | **vRNA yield $EC_{50}$:** N.a. |
| **Anti viral activity $EC_{90}$:** N.a. | **vRNA yield $EC_{90}$:** N.a. |
| **Cytotoxicity $CC_{50}$:** N.a. | **SI CC50/EC50:** N.a. |
| **Native MS adduct saturation:** - | **Native MS adduct size:** N.a. |

### Hit#34: Necrostatin-1 bound to SARS-CoV-2 MPro, not deposited

| | |
|---|---|
| **Synonyms:** Necrostatin-1, MTH-DL-Tryptophan, Nec-1 | Lig-Plot interaction network |
| **PubChem CID:** 2828334 | |
| **CAS:** 4311-88-0 | |
| **Molecular weigth:** 259.33 g/mol | |
| **Binding type:** non-covalent | |
| **Binding location:** surface | |
| **Screen compound ID:** SPE_A06935312 | |
| **Original compound:** | |

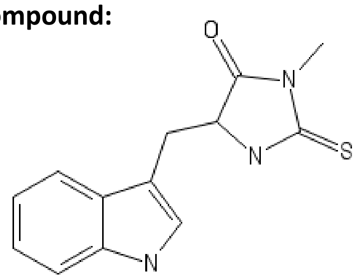


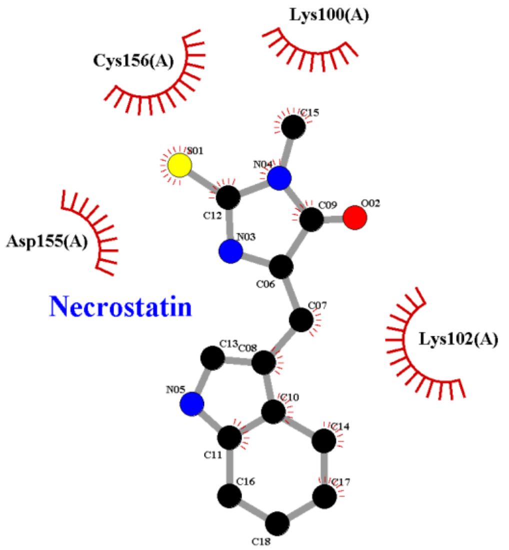


**Isomeric smiles:** CN1C(=O)C(NC1=S)CC2=CNC3=CC=CC=C32

**Refinement 2Fo-Fc map**, 0.7 σ-level

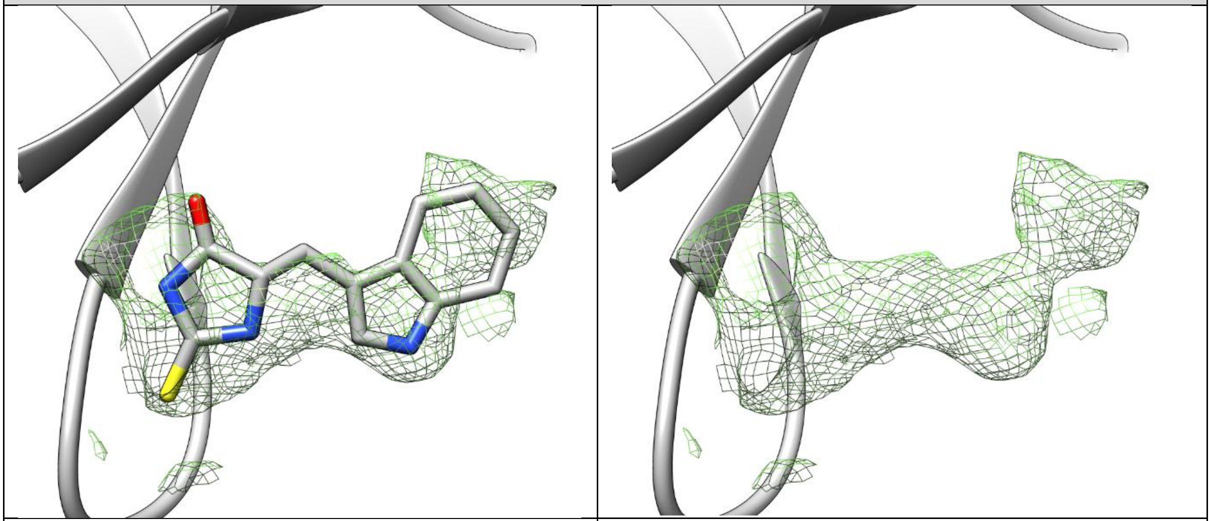

| | |
|---|---|
| **PDB structure code:** - | **PDB Ligand code:** - |
| **Resolution:** 1.7 Å | $\mathbf{R_{work}/R_{free}}$: - |
| **Occupancy:** - | **Ligand RSCC/RSR:** - |

**Biochemistry & Cell biology**

| | |
|---|---|
| **Anti viral activity** $\mathbf{EC_{50}}$: N.a. | **vRNA yield** $\mathbf{EC_{50}}$: N.a. |
| **Anti viral activity** $\mathbf{EC_{90}}$: N.a. | **vRNA yield** $\mathbf{EC_{90}}$: N.a. |
| **Cytotoxicity** $\mathbf{CC_{50}}$: N.a. | **SI CC50/EC50:** N.a. |
| **Native MS adduct saturation:** - | **Native MS adduct size:** N.a. |

#### Hit#35: Ipidacrine bound to SARS-CoV-2 MPro, PDB: 7AF0

**Synonyms:** Ipidacrine, Amiridine, 2,3,5,6,7,8-Hexahydro-1H-cyclopenta[b]quinolin-9-ylamine

**PubChem CID:** 604519

**CAS:** 62732-44-9

**Molecular weigth:** 188.27 g/mol

**Binding type:** non-covalent

**Binding location:** surface

**Screen compound ID:** SPE_K66896231

**Original compound:**

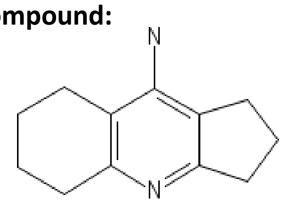


Lig-Plot interaction network

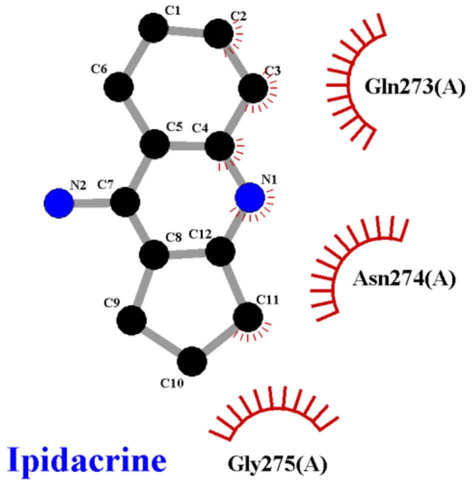


**Isomeric smiles:** C1CCC2=C(C1)C(=C3CCCC3=N2)N

**Refinement 2Fo-Fc map**, 0.8 σ-level

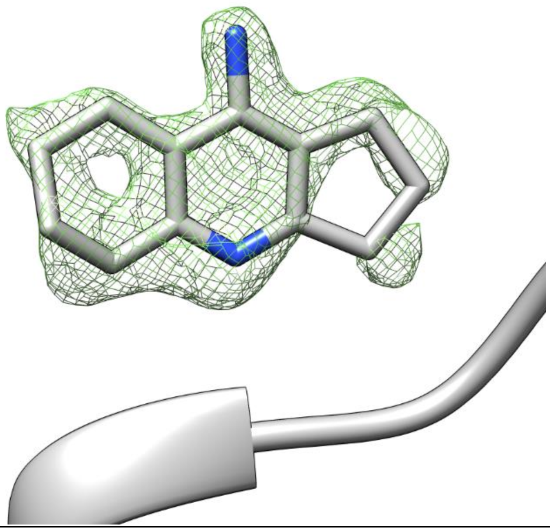

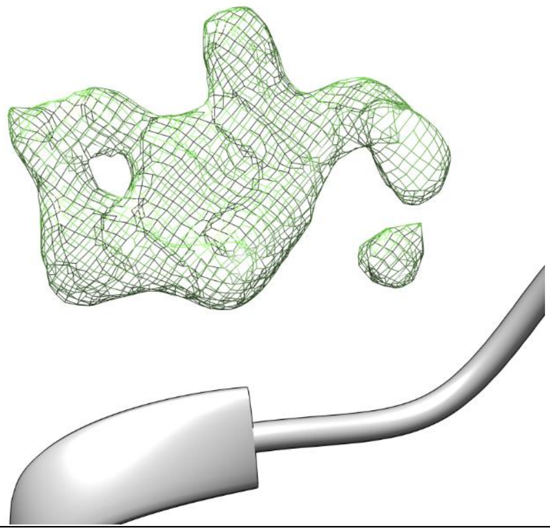

| | |
|---|---|
| **PDB structure code:** 7AF0 | **PDB Ligand code:** R9W |
| **Resolution:** 1.7 Å | $\mathbf{R_{work}/R_{free}}$: 0.18/0.23 |
| **Occupancy:** 0.61 | **Ligand RSCC/RSR:** 0.74/0.22 |

**Biochemistry & Cell biology**

| | |
|---|---|
| **Anti viral activity $EC_{50}$:** N.a. | **vRNA yield $EC_{50}$:** N.a. |
| **Anti viral activity $EC_{90}$:** N.a. | **vRNA yield $EC_{90}$:** N.a. |
| **Cytotoxicity $CC_{50}$:** N.a. | **SI CC50/EC50:** N.a. |
| **Native MS adduct saturation:** - | **Native MS adduct size:** N.a. |

### Hit#36: AZD6482 bound to SARS-CoV-2 MPro, PDB: 6YVF

**Synonyms:** AZD6482, 1173900-33-8
**PubChem CID:** 44137675
**CAS:** 1173900-33-8
**Molecular weigth:** 408.4 g/mol
**Binding type:** non-covalent
**Binding location:** surface
**Screen compound ID:** SPE_K58772419
**Interaction:**

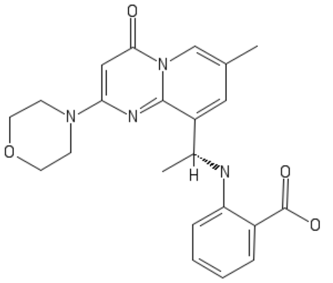

Lig-Plot interaction network

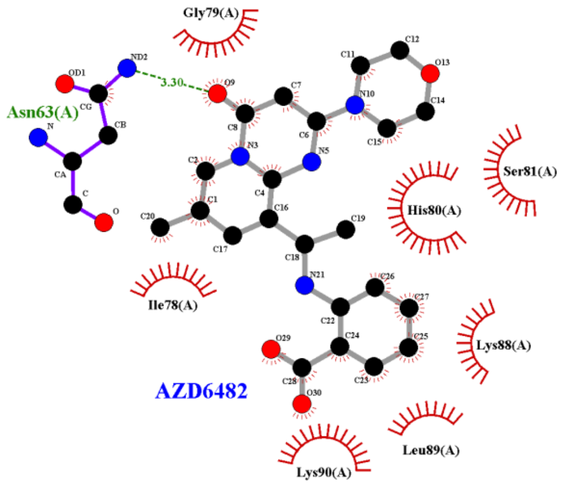


**Isomeric smiles:** CC1=CN2C(=O)C=C(N=C2C(=C1)[C@@H](C)NC3=CC=CC=C3C(=O)O)N4CCOCC4

**Refinement 2Fo-Fc map**, 1.0 σ-level

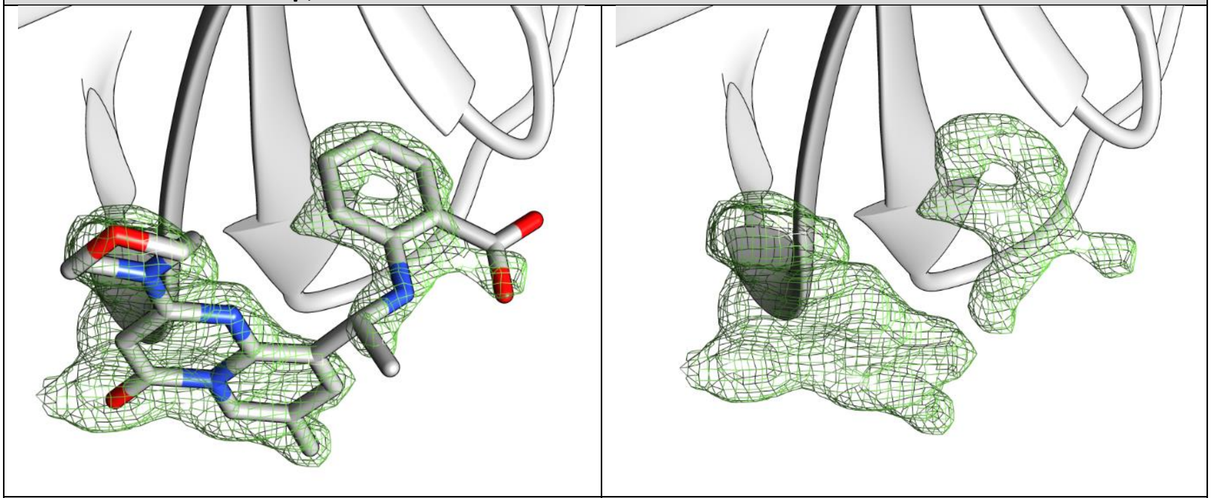

| | |
|---|---|
| **PDB structure code:** 6YVF | **PDB Ligand code:** A82 |
| **Resolution:** 1.63 Å | **$R_{work}$/$R_{free}$:** 0.19/0.21 |
| **Occupancy:** 0.71 | **Ligand RSCC/RSR:** 0.81/0.13 |

**Biochemistry & Cell biology**

| | |
|---|---|
| **Anti viral activity $EC_{50}$:** N.a. | **vRNA yield $EC_{50}$:** N.a. |
| **Anti viral activity $EC_{90}$:** N.a. | **vRNA yield $EC_{90}$:** N.a. |
| **Cytotoxicity $CC_{50}$:** N.a. | **SI CC50/EC50:** N.a. |
| **Native MS adduct saturation:** 24% (62/29/9) | **Native MS adduct size:** 408 Da |

### Hit#37: Polydatin bound to SARS-CoV-2 MPro, not deposited

**Synonyms:** Polydatin, Piceid, Trans-Piceid

**PubChem CID:** 5281718

**CAS:** 27208-80-6

**Molecular weigth:** 390.4 g/mol

**Binding type:** Non-covalent

**Binding location:** surface

**Screen compound ID:** DOM_SIM_161

**Original compound:**

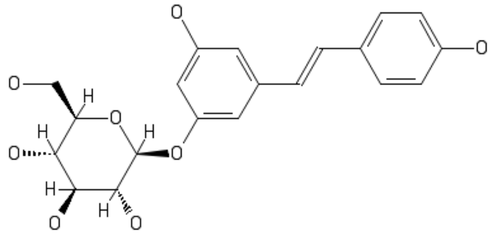

Lig-Plot interaction network

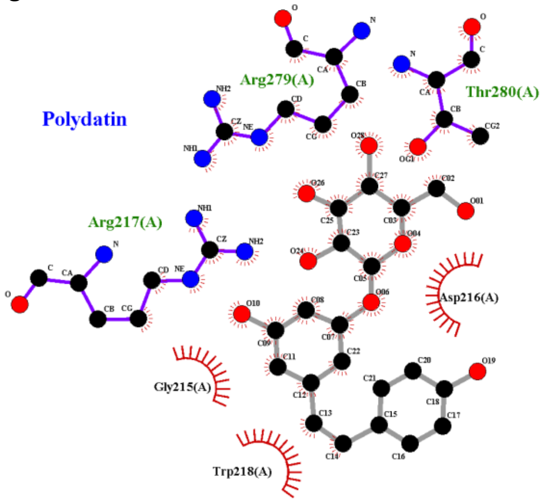


**Isomeric smiles:**
C1=CC(=CC=C1/C=C/C2=CC(=CC(=C2)O[C@H]3[C@@H]([C@H]([C@@H]([C@H](O3)CO)O)O)O)O)O

**PanDDA event map BDC 0.25** , 1.5 σ-level

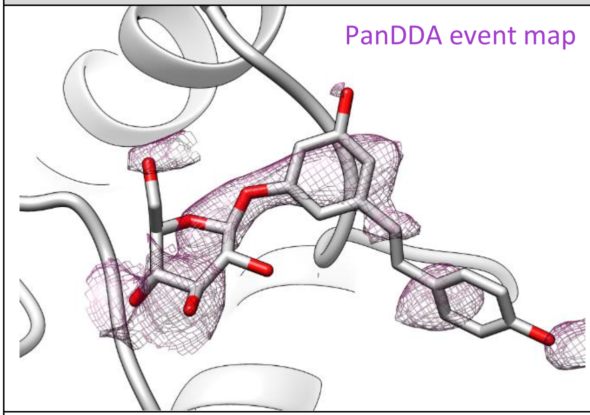


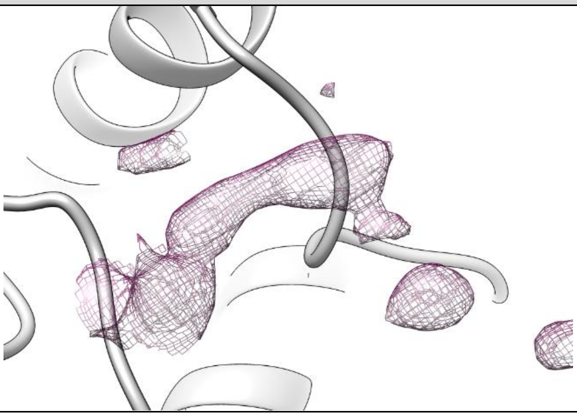

| | |
|---|---|
| **PDB structure code:** - | **PDB Ligand code:** - |
| **Resolution:** 1.7 Å | **$R_{work}$/$R_{free}$:** - |
| **Occupancy:** - | **Ligand RSCC/RSR:** - |

**Biochemistry & Cell biology**

| | |
|---|---|
| **Anti viral activity $EC_{50}$:** N.a. | **vRNA yield $EC_{50}$:** N.a. |
| **Anti viral activity $EC_{90}$:** N.a. | **vRNA yield $EC_{90}$:** N.a. |
| **Cytotoxicity $CC_{50}$:** N.a. | **SI CC50/EC50:** N.a. |
| **Native MS adduct saturation:** 5% (91/9/0) | **Native MS adduct size:** 390 Da |
